# Indole-3-propionic acid nano-prodrugs sustain increased lymphoid exposure and alleviate experimental autoimmune arthritis

**DOI:** 10.64898/2026.10.13.759006

**Authors:** Yitong Guo, Huiqi Wang, Brad Cleveland, Ho Yeung Leung, Dinh Chuong Nguyen, Jack Thomson Smith, J. Emiliano Gómez Medellín, Giulia E. Stewart, Raul Johnson, Lindsey M. Williams, Xiaoxing Wang, Peng Qiu, Tomas Mustelin, Shijie Cao

## Abstract

Chronic inflammatory diseases, including rheumatoid arthritis (RA), currently require sustained control of inflammation. Although long-acting therapies can reduce dosing frequency, those that rely on systemic immunosuppression may still compromise protective immunity. Microbiota-derived metabolites offer a complementary starting point for immune modulation through naturally occurring host–microbe signals. Indole-3-propionic acid (IPA), a gut microbial tryptophan metabolite naturally present in human circulation, has newly recognized immunoregulatory activity, yet its effects across disease contexts and its potential for therapeutic use remain incompletely defined. Here, we developed polymeric IPA nano-prodrugs (pIPA_NP_) by conjugating IPA to a methacrylamide polymer through ester linkages and formulating the polymer into injectable nanoparticles. Following subcutaneous administration of fluorescently-labeled pIPA_NP_, injection-site fluorescence remained detectable for at least 35 days, with labeled material detected in multiple lymph nodes. LC-MS/MS confirmed higher lymph-node IPA concentrations in mice treated with the pIPA_NP_s compared to free IPA control at every sampled time point over 14 days. In collagen antibody-induced arthritis (CAIA), pIPA_NP_s reduced clinical disease severity and paw swelling under both prophylaxis and therapeutic administration. pIPA_NP_ treatment increased the relative representation of CD11c⁺CD103⁺CD11b⁻ cells with a conventional dendritic cell (cDC)1-associated phenotype and reduced costimulatory-marker frequencies in both CD11c⁺CD103⁺ and CD11c⁺CD11b⁺ populations, most prominently in hock-draining lymph nodes. These changes were accompanied by increased PD-1⁺ and IFN-γ⁺ frequencies among CD4⁺ and CD8⁺ T cells.

Fluorescent nanoparticle tracking showed preferential association with several myeloid populations, while single-cell profiling revealed divergent interferon-response signatures across splenic populations. These findings reveal arthritis-associated immunomodulatory activity of polymeric IPA and support its further development as a microbiome-inspired, long-acting approach to inflammatory arthritis.

## 1. Introduction

Rheumatoid arthritis (RA) requires sustained control of inflammation to prevent joint damage and preserve physical function. Current disease-modifying antirheumatic drugs (DMARDs) include conventional agents such as methotrexate; biologics that inhibit TNF or IL-6 signaling, block T-cell costimulation, or deplete B cells; and synthetic Janus kinase (JAK) inhibitors (*1*). These therapies have substantially improved care, but persistent disease activity and treatment-related adverse effects still require dose adjustment, combination therapy, or switching agents. Glucocorticoids can provide additional control, although cumulative toxicity limits prolonged use (*2*). Some therapies also affect protective immune responses: for example, rituximab reduces antibody responses to certain vaccines (*3*). These limitations motivate the development of therapeutic approaches that combine effective inflammatory control with a favorable distribution and duration of drug exposure.

Long-acting injectables are already possible with established RA treatments (*4*). For example, subcutaneous golimumab is administered monthly, and rituximab courses can be repeated at approximately 6-month intervals. However, extending drug exposure does not itself separate control of arthritis from effects on normal immune function. The pathways targeted by these agents also contribute to host immune defense. For example, Golimumab inhibits TNFα and carries a boxed warning for serious infections. Rituximab depletes CD20^+^ B cells and can reduce antibody responses to certain vaccines (*5–7*). A depot formulation can reduce administration frequency, but exposure that persists after dosing also reduces flexibility when treatment must be interrupted. Thus, the opportunity for a new long-acting therapy lies in coupling sustained disease control with a favorable distribution of exposure and a therapeutic window that limits adverse effects while preserving protective immunity.

Microbial metabolites naturally present in the host provide a biologically grounded starting point for exploring such therapies. Short-chain fatty acids, e.g. butyrate, promotes the differentiation of colonic regulatory T cells, demonstrating how a product of microbial metabolism can influence host immune responses (*8*, *9*). Translating these activities into therapies requires delivery strategies that control where and how the metabolites become available. Our previous work showed that polymeric micelles releasing butyrate in the intestine reduced peanut-induced anaphylaxis and ameliorated experimental colitis in mice (*10*). We also developed a serine-conjugated butyrate prodrug with improved oral bioavailability that suppressed autoimmune arthritis and neuroinflammation (*11*). These studies demonstrate that chemical modification and formulation can improve the therapeutic activity of microbial metabolites and support extending this approach to other microbial immunomodulators.

Indole-3-propionic acid (IPA) is a tryptophan metabolite produced by members of the gut microbiota that enters the circulation, linking intestinal microbial metabolism with host physiology (*12*, *13*). Previous studies implicated IPA in the regulation of intestinal barrier function through pregnane X receptor signaling (*14*). More recently, Li and colleagues showed that IPA enhanced mitochondrial respiration in CD4⁺ T cells through changes in fatty acid and amino acid oxidation. This metabolic response involved PPARβ/δ signaling, limited Th1 and Th17 differentiation, and supported protection against experimental intestinal inflammation (*15*). Additionally in a more relevant collagen-induced arthritis (CIA) study, an orally administered mixture of IPA and indole-3-acetic acid reduced arthritis severity and joint damage, although that study did not resolve the contribution of IPA alone (*16*). These findings support investigating IPA as an immunomodulatory therapeutic, while leaving open how its delivery and chemical presentation influence activity in inflammatory arthritis.

Delivery to lymphatic tissues offers an opportunity to bring microbial metabolites into compartments where antigen-presenting cells and lymphocytes interact. Previous studies demonstrated that nanoparticle size influences lymphatic transport, lymph-node retention, and access to dendritic cells after interstitial administration (*17–19*). In this study, we developed polymeric IPA nano-prodrugs (pIPA_NP_) in which IPA is conjugated to a methacrylamide polymer through ester linkages. We characterized the pIPA_NP_, its distribution following subcutaneous injection, and IPA exposure in lymph nodes and plasma. We then evaluated its activity in preventive and therapeutic settings of collagen antibody-induced arthritis (CAIA), including administration at the belly or near the hock site, after disease onset. pIPA_NP_ treatment reduced clinical arthritis severity, paw swelling, and local IL-6 concentrations, accompanied by changes in myeloid activation markers and T-cell phenotypes in lymph nodes and spleen. Cellular distribution studies and exploratory single-cell RNA sequencing further characterized the immune populations associated with treatment. These experiments examine how polymeric delivery of a microbial tryptophan metabolite alters its tissue exposure and supports anti-inflammatory activity in antibody-induced arthritis.

## 2. Results

### Synthesis and characterization of pIPA_NP_

In this study, we developed an injectable nano-prodrug formulation for prolonged IPA delivery to lymphatic tissues. We first synthesized a polymeric IPA prodrug (pIPA) in which IPA is covalently conjugated to a methacrylamide backbone through ester bonds (**Fig. 1a**). An IPA-containing monomer was synthesized and polymerized by reversible addition-fragmentation chain-transfer (RAFT) polymerization. Monomer and polymer structures were characterized by ¹H nuclear magnetic resonance spectroscopy, and gel permeation chromatography (GPC) showed a number-average molecular weight of approximately 15.9 kDa with a dispersity of 1.07 (**Figs. S1** and **S2**).

**Figure 1.**
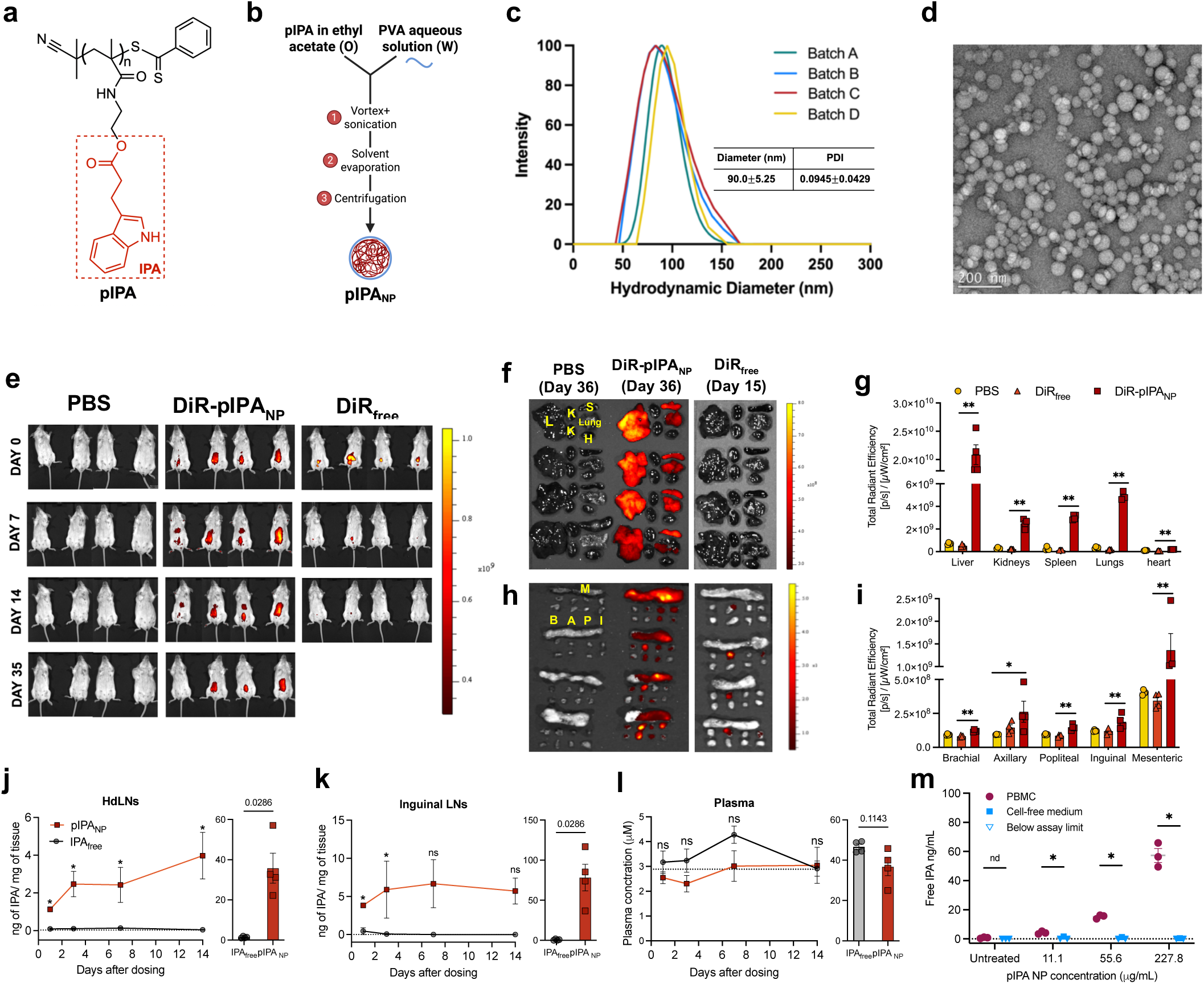
pIPA_NP_ platform formulation, characterization, and biodistribution. **a.** Structure of indole-3-propionic acid polymer (pIPA). **b**. Formulation process of pIPA_NP_ from pIPA. **c.** Hydrodynamic diameter distribution of four independent batches of pIPA_NP_ determined by DLS. **d**. Transmission Electron Microscopy (TEM) image of the pIPA_NP_ at 1:10 dilution. **e**. Live imaging of mice injected subcutaneously with PBS, DiR-pIPA_NP_, or fluorescently matched DiR_free_ on Day 0, 14, and 35 (n = 4) using IVIS. **f.** *Ex vivo* fluorescence images of the mouse liver (L), kidneys (K), spleen (S), lung, and heart (H) taken on Day 36 or Day 15. **g.** *Ex vivo* fluorescence images of the mouse mesenteric (M), inguinal (I), popliteal (P), axillary (A), and brachial (B) lymph nodes taken on Day 36 or Day 15. **h.** Total radiant efficiency ([p/s]/[μW/cm^2^]) of *ex vivo* tissues and **i.** lymph nodes. **j.** IPA concentration (ng IPA/mg tissue) and area under the curve (AUC) in hock-draining lymph nodes (HdLNs; pooled popliteal, axillary, and brachial) and **k.** in inguinal lymph nodes (injection site-draining lymph nodes) at days 1, 3, 7, and 14 following subcutaneous administration of pIPA_NP_ or IPA_free_ (n=4 per timepoint) . **l.** Plasma IPA concentration (μM) in pIPA_NP_- and IPA_free_-treated mice at days 1, 3, 7, and 14 post-administration. Dotted line indicates baseline IPA levels measured in control mice (n=4). Values below the LLOQ but above the LOD were assigned LLOQ/2; concentrations below the zero calibrator were assigned 0. **m.** IPA released from pIPANP after 4 h incubation with human peripheral blood mononuclear cells (PBMC) or in cell-free media at three concentrations (n=3). All statistical comparisons were performed using nonparametric tests. Data are presented as mean ± SEM. \**p* < 0.05, \*\**p* < 0.01.

We proceeded by formulating pIPA into nanoparticles (pIPA_NP_) using a single-emulsion solvent evaporation method (**Fig. 1b**). During the optimization stage, pIPA_NP_ size was tuned to below 100 nm, a size threshold shown to facilitate efficient lymphatic uptake (*20*), which is also our target of interest for this study. Four independent batches showed reproducible and homogeneous hydrodynamic size distributions, with a mean diameter of 90.0 ± 5.25 nm and a mean polydispersity index of approximately 0.095 ± 0.043 (**Fig. 1c**).

Transmission electron microscopy (TEM) showed spherical particles (**Fig. 1d**). Particle diameter remained below 100 nm during 56 days of storage in water at either 4°C or room temperature, and zeta potential measurements showed a near-neutral surface charge (**Fig. S3**).

### pIPA_NP_ sustains increased IPA exposure in lymph nodes after subcutaneous administration

To evaluate the biodistribution and retention of pIPA_NP_ *in vivo*, a longitudinal fluorescence imaging study was conducted. Due to differences in expected retention times, DiR-labeled pIPA_NP_ and free DiR were administered at different time points but sacrificed simultaneously, allowing direct *ex vivo* comparison of tissue fluorescence under equivalent conditions. Longitudinal *in vivo* imaging demonstrated that DiR-labeled pIPA_NP_ produced detectable signal for up to 35 days following a single subcutaneous injection, whereas free DiR signal fell below the detectable threshold by approximately day 14 (**Fig. 1e**), confirming the extended retention conferred by nanoparticle formulation. *Ex vivo* organ analysis at day 36 revealed that pIPA_NP_ were broadly distributed across major organs, with the highest accumulation in the liver, moderate signals in the spleen, kidneys, and lungs, and negligible signal in the heart, consistent with reticuloendothelial system-mediated clearance (*21*) (**Fig. 1f, g**). In contrast, the free DiR group exhibited fluorescence signals comparable to the PBS-treated group at day 15, indicating that the DiR signal had declined to tissue background level by this time point.

*Ex vivo* fluorescence analysis of lymphoid tissues confirmed that DiR-pIPA_NP_ accumulated in all draining lymph nodes (**Fig. 1h, i**), with fluorescence detectable in most lymph nodes at day 36, markedly exceeding free DiR signals at day 15 under equivalent imaging conditions. Notably, fluorescence in the free DiR group was largely restricted to axillary lymph nodes, whereas pIPA_NP_ achieved broader lymph node distribution, including accumulation in hock-draining popliteal lymph nodes. The hock-draining lymph nodes drain the inflamed paws and represent the immune compartment most relevant to joint inflammation in RA.

To quantitatively confirm lymph node drug exposure, IPA tissue concentrations were measured by LC-MS/MS at days 1, 3, 7, and 14 following one subcutaneous administration of pIPA_NP_ or sodium indole-3-propionate (NaIPA; hereafter, free IPA). IPA was detected in both hock-draining and inguinal lymph nodes of pIPA_NP_-treated animals across all four time points, reaching 3.98 ng/mg and 5.71 ng/mg tissue at day 14, respectively (**Fig. 1j, k**). In contrast, IPA remained below the lower limit of quantification (LLOQ) in the free IPA and untreated groups (dotted line) at all time points. Plasma IPA levels were measured to assess systemic exposure and remained comparable to baseline (dotted line) in both the free IPA and pIPA_NP_ groups, with slightly elevated but not statistically significant levels in the free IPA group (**Fig. 1l**). These findings confirm that the pIPA_NP_ nano-prodrug formulation is essential for achieving measurable and sustained drug exposure in secondary lymphoid tissues.

Based on the observation that pIPA_NP_ released IPA in the lymph nodes, we conducted a release kinetics study from the pIPA_NP_s. In cell culture media supplemented with 10% non-heat-inactivated bovine fetal serum at 37 °C, the IPA release was below 0.20% over 14 days (**Fig. S3c**). We hypothesized that the release of IPA might be immune cell-mediated, especially after the pIPA_NP_s are taken up by innate immune cells. To test this hypothesis, we measured IPA release during a 4-hour incubation with human peripheral blood mononuclear cells (PBMCs) isolated from an RA patient. Free IPA concentrations increased with increasing pIPA_NP_ dose in PBMC-containing samples, whereas cell-free controls at matched pIPA_NP_ concentrations remained below the assay limit (**Fig. 1m**). These findings suggest that PBMC-associated processes may facilitate IPA release from pIPA NPs, although the responsible cell populations and release mechanisms remain to be determined.

### pIPA_NP_ prophylaxis reduces arthritis incidence, severity, and joint inflammation

Having established that pIPA_NP_ sustain increased IPA exposure in draining lymph nodes, we next investigated whether pIPA_NP_ prophylaxis could prevent joint inflammation in the collagen antibody-induced arthritis (CAIA) model. CAIA is initiated through passive administration of arthritogenic anti-collagen antibodies, allowing evaluation of the antibody-driven effector phase of arthritis without requiring active immunization against collagen (*22*). Female BALB/c mice received subcutaneous injections of pIPA_NP_, free IPA (IPA_free_), or PBS vehicle control one and seven days prior to arthritogenic antibody injection (**Fig. 2a**). IPA_free_ and pIPA_NP_ were administered at an equivalent IPA dose of 2 mg per mouse per injection, establishing exposure before disease induction. Healthy mice were included as a baseline non-inflammatory reference. Clinical arthritis scores were assessed using a swelling-based scale of 0–4 for each paw, yielding a total score of 0–16 per mouse (**Table S1**). Following disease induction, PBS- and IPA_free_-treated mice developed progressive arthritis with similar clinical scores. In contrast, pIPA_NP_ prophylaxis limited the increase in clinical arthritis scores and the animals exhibited lower disease severity throughout the study (**Fig. 2b**). Additionally, the difference in disease severity was reflected in cumulative arthritis scores over days 0–11 post induction, which were significantly lower in the pIPA_NP_ group than in the PBS and IPA_free_ groups (*P* < 0.01 and *P* < 0.05, respectively; **Fig. 2c**).

**Figure 2.**
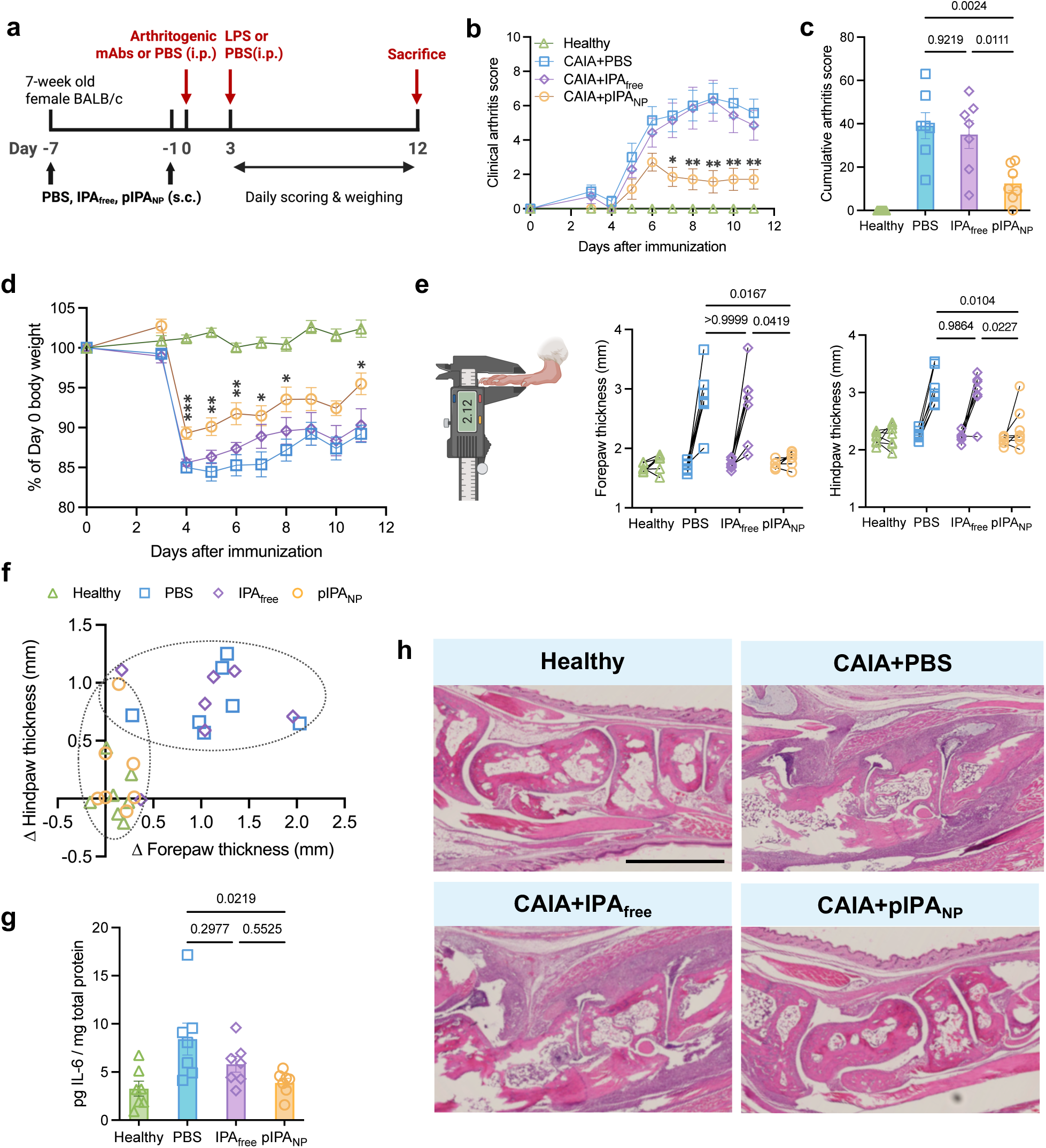
pIPA_NP_ reduced the disease severity of CAIA in a prophylactic setting. **a.** Experimental timeline. **b.** Clinical arthritis scores of mice from Day 0 to Day 11 **c.** The average cumulative arthritis score (n=7) from Day 0 to Day 11. **d.** Daily body weight expressed as a percentage of Day 0 body weight. **e.** Average forepaw and hindpaw thickness at Day 0 vs. Day 11. Statistical comparisons were performed on the change in thickness (Δ=Day 11−Day 0) between groups. **f.** Scatter plot of individual mice showing Δ forepaw vs. Δ hindpaw thickness (Day 11−Day 0), colored by group. **g.** ELISA for IL-6 levels in mouse paw homogenates, normalized by total protein content. h. Representative histological images of mouse joint sections stained with hematoxylin and eosin (H&E), harvested on Day 12 . Statistical comparisons were performed using ordinary one-way ANOVA with Tukey’s multiple comparisons test or the Kruskal-Wallis test with Dunn’s post hoc test, depending on data distribution. *p < 0.05, **p < 0.01, ***p < 0.001, ****p < 0.0001; ns, not significant.

Although all animals in arthritic groups lost weight after disease induction, pIPA_NP_-treated mice exhibited a less pronounced weight loss and a greater recovery during the follow-up period compared to the PBS and IPA_free_ groups (**Fig. 2d**). To quantitatively corroborate joint swelling using an independent measurement besides clinical scores, forepaw and hindpaw thickness were measured before disease induction and on day 11 post induction. PBS- and IPA_free_-treated mice exhibited substantial increases in paw thickness, whereas pIPA_NP_-treated mice had no swelling or reduced increases in both forepaws and hindpaws (**Fig. 2e**). Graphing the increases in forepaw and hindpaw thickness for individual mice further showed that the pIPA_NP_ group clustered with the baseline, non-inflamed mice, while the IPA_free_-treated group clustered with the mock-treated PBS group generally exhibiting increased swelling (**Fig. 2f**). Normalized IL-6 concentrations in paw homogenates were significantly lower in pIPA_NP_-treated mice compared to PBS-treated arthritic mice (P < 0.05; Fig. 2g), whereas IPA_free_-treated mice exhibited a moderate, not significant change from the mock-treated control group (**Fig. 2g**). Histological analysis provided an additional assessment of inflammation within the affected joints.

Hemaetoxylin and eosin (H&E)-stained sections collected on day 12 post arthritogenic antibody injection showed prominent inflammatory-cell infiltration in mock PBS- and IPA_free_-treated mice, while sections from pIPA_NP_-treated mice showed reduced infiltration (**Fig. 2h**, and **Fig. S4**). Overall, these findings demonstrate that pIPANP prophylaxis attenuates the clinical and tissue manifestations of antibody-induced arthritis, with greater protection than dose-matched free IPA and concordant reductions in disease severity, paw swelling, and local joint inflammation.

### Disease protection is accompanied by changes in dendritic cell and T cell phenotypes

To examine immune changes associated with pIPA_NP_-mediated disease protection, we analyzed myeloid and T-cell populations in the spleen, hock-draining lymph nodes (hdLNs) draining the affected paws, and inguinal lymph nodes (ingLNs) draining the injection site from the CAIA prophylaxis study. Flow cytometry revelaed (**Fig. S5) that** pIPA_NP_ prophylaxis significantly reduced the frequencies of CD86^+^, CD80^+^, CD40^+^, and MHC-II^+^ cells within CD11b^+^f4/80^+^, CD11b^+^CD11c^-^, CD11b^+^CD11c^+^, CD11c^+^, and CD103^+^CD11c^+^ populations in hdLNs compared with both PBS and IPA_free_ (**Fig. 3a and Fig. S8**). These coordinated reductions indicated a less activated dendritic cell phenotype in the lymph nodes associated with joint inflammation. Changes in the spleen and ingLNs were more selective. In the spleen, pIPA_NP_ reduced CD40⁺ and MHC-II⁺ frequencies within the CD11b⁺CD11c⁺ population compared with PBS, while CD103⁺CD11c⁺ cells showed reductions in several costimulatory markers and in MHC-II (**Fig. S7**). In ingLNs, CD103⁺CD11c⁺ cells exhibited lower CD80⁺ and MHC-II⁺ frequencies compared with both arthritic control groups (**Fig. S9**). CD11b⁺CD11c⁻ cells showed few marker reductions across the three tissues, whereas splenic CD11b⁺F4/80⁺ cells exhibited increased CD86⁺ and CD80⁺ frequencies (**Figs. S7–S9**).

**Figure 3.**
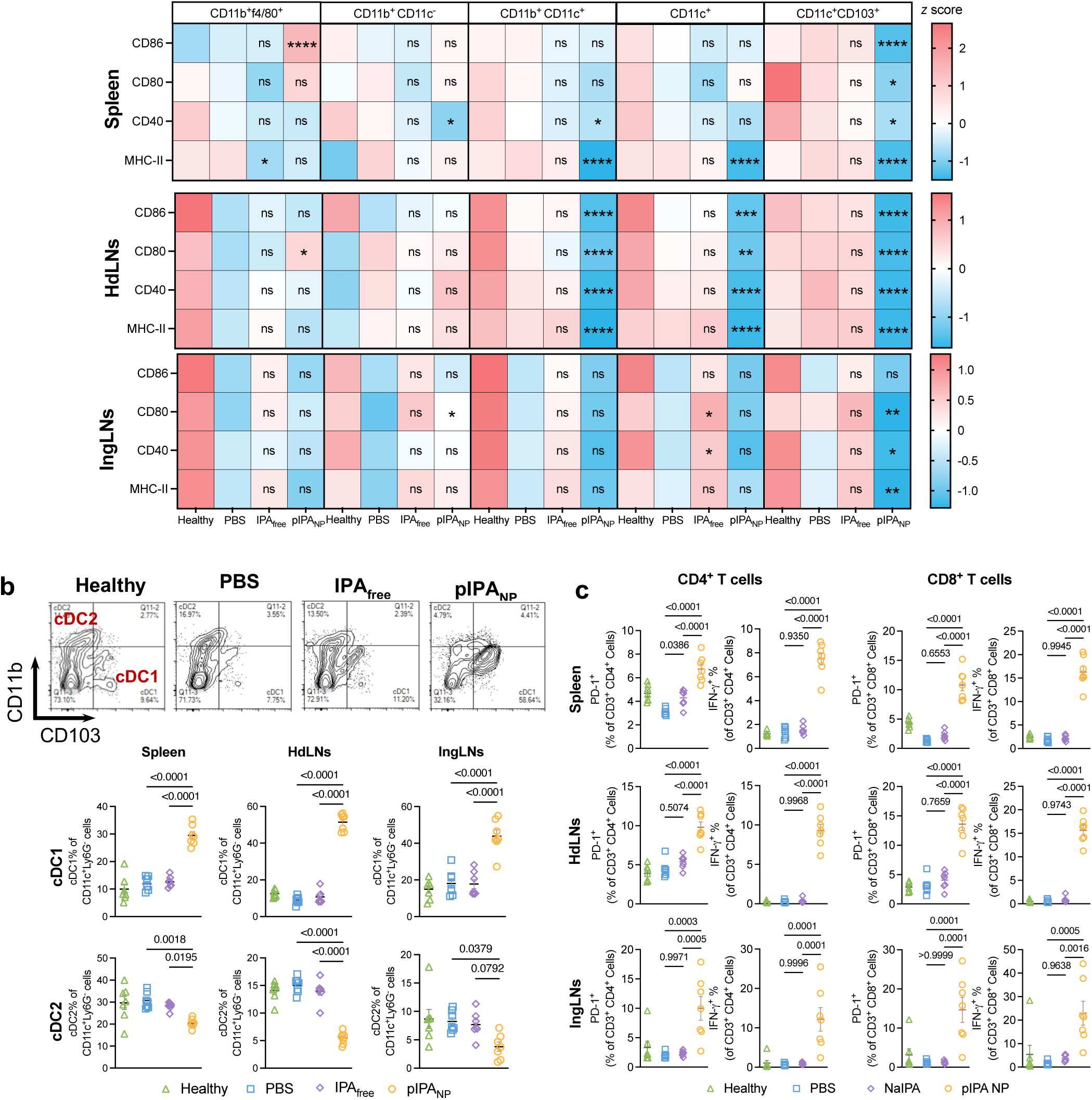
pIPA_NP_ treatment promotes a tolerogenic phenotype in both myeloid and T cell compartments across immune tissues. a. Heat map of z-score-normalized % of CD86+, CD80+, CD40+, and MHC-II+ cells within CD11b+f4/80+, CD11b+CD11c-, CD11b+CD11c+, CD11c+ CD11b-, and CD103+CD11c+ cells in spleen, hock-draining lymph nodes (hdLNs), and inguinal lymph nodes (ingLNs) (z=(x−mean)/SD). **b.** Representative flow cytometry plots with quadrant gating on CD11b and CD103, showing the percentages of CD11b^-^CD103^+^ (cDC1) and CD11b⁺CD103^-^ (cDC2) cells (left), and quantification of each population (right) in spleen, hock-draining lymph nodes (hdLNs), inguinal lymph nodes (ingLNs) **c.** Percentage of PD-1+ and IFN-γ+ cells among CD4+ and CD8+ cells in spleen, hdLNs, and ingLNs. Statistical significance was determined by ordinary one-way ANOVA with Tukey’s multiple comparisons test: *p < 0.05, **p < 0.01, ***p < 0.001, ****p < 0.0001.

We further evaluated dendritic cell composition by quantifying CD11b and CD103 expression within the CD11c⁺Ly6G⁻ compartment (**Fig. S6**). pIPA_NP_ significantly increased the proportion of CD11b⁻CD103⁺ cells, corresponding to the cDC1, in the spleen, hdLNs, and ingLNs compared with PBS and IPA_free_, with the largest proportional shift occurring in the hdLNs (**Fig. 3b**). The CD11b⁺CD103⁻ fraction, corresponding to cDC2 populations, decreased significantly in both the spleen and the hdLNs.

T-cell profiling showed that pIPA_NP_ increased PD-1⁺ frequencies among both CD4⁺ and CD8⁺ T cells in the spleen, hdLNs, and ingLNs compared with PBS and IPA_free_. Intracellular IFN-γ⁺ frequencies were also increased in both T-cell populations across these tissues (**Fig. 3c**; **gating in Fig. S10**). Increased PD-1 expression can accompany T-cell activation and does not independently establish exhaustion or functional tolerance (*23*). IFN-γ has been shown to restrain IL-17-associated responses in collagen-induced arthritis (*24*), providing a possible context for its increase in protected animals, although whether this mechanism contributes to protection in the present CAIA study remains to be investigated.

### Therapeutic administration of pIPA_NP_ reduced symptoms in models of established arthritis and altered myeloid phenotypes

After establishing the efficacy of pIPA_NP_ in a prophylactic setting, we proceeded by asking whether pIPA_NP_ could improve symptoms in animals with established arthritis. To that end, we administered PBS or DiD-loaded pIPA_NP_ (2 mg IPA-equivalent per injection) subcutaneously to CAIA arthritic mice on days 6 and 7 post arthritogenic antibody injection, after clinical arthritis and paw swelling had developed (**Fig. 4a**). DiD loading allowed us to track the cellular biodistribution of the pIPA_NP_s. In a separate study with unlabeled pIPA_NP_s, we observed similar clinical and flow cytometry outcomes (Supplementary Fig. S11). Following treatment, clinical arthritis scores declined in pIPA_NP_-treated mice, while scores in PBS-treated animals continued to increase.

**Figure 4.**
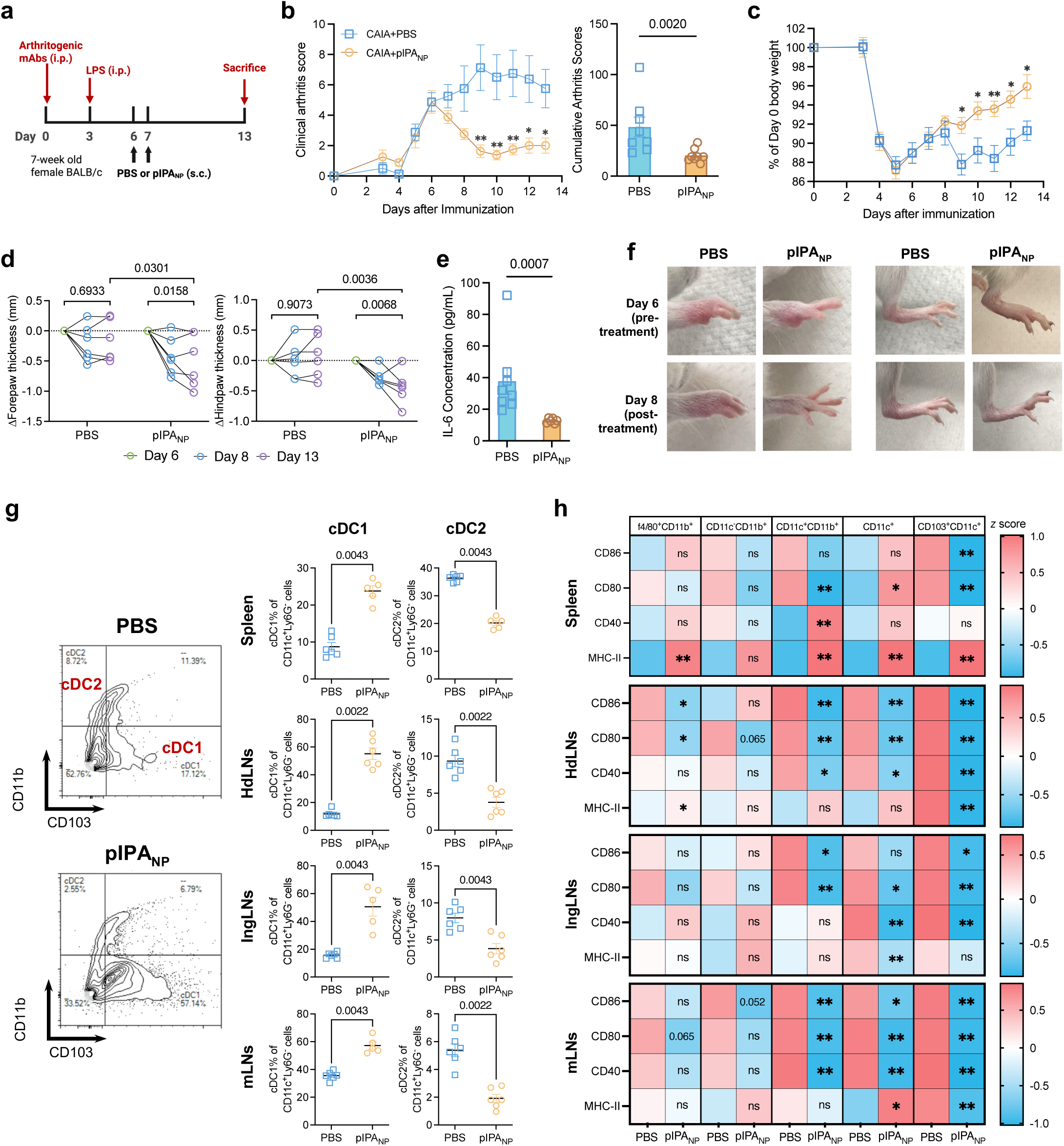
pIPA_NP_ reversed CAIA after disease induction and reduced myeloid activation. **a.** Experimental timeline. **b**. Clinical arthritis scores of mice from Day 0 to Day 13 and the average cumulative arthritis score of each subject (n=8). **c**. Daily body weight expressed as a percentage of Day 0 body weight. **d**. ΔForepaw and hindpaw thickness on Day 8, and Day 13 with reference to Day 6 thickness. Two-way ANOVA was used for statistical testing. **e.** ELISA for IL-6 levels in mouse paw homogenates. **f.** Representative photos of matched mouse front and back paws before and after treatment. **g.** Representative flow cytometry plots with quadrant gating on CD11b and CD103, showing the percentages of CD11b^-^ CD103^+^ (cDC1) and CD11b⁺CD103^-^ (cDC2) cells (left), and quantification of each population (right) in spleen, hock-draining lymph nodes (hdLNs), inguinal lymph nodes (ingLNs), and mesenteric lymph nodes (mLNs) (n=6). **h.** Heat map of z-score-normalized % of CD86^+^, CD80^+^, CD40^+^, and MHC-II^+^ cells within CD11b^+^f4/80^+^, CD11b^+^CD11c^-^, CD11b^+^CD11c^+^, CD11c^+^ CD11b^-^, and CD103^+^CD11c^+^ cells in spleen, hdLNs, ingLNs, and mLNs (*z*=(x−mean)/SD). Data are presented as mean ± SEM. Statistical comparisons were performed using the two-tailed Mann-Whitney U test. *p < 0.05, **p < 0.01, ***p < 0.001, ****p < 0.0001; ns, not significant.

Although scores increased modestly late in the observation period, pIPA_NP_-treated mice maintained lower disease severity through day 14 and had significantly lower cumulative arthritis scores (**Fig. 4b**). The improvement in clinical scores was accompanied by better body-weight recovery (**Fig. 4c**). Both forepaw and hindpaw thickness decreased significantly from their pretreatment measurement on day 6 in pIPA_NP_-treated mice, and these reductions persisted through the final measurement. As expected, mock PBS-treated animals showed no significant improvement over time (**Fig. 4d**). Matched mouse paws showed reduced visible paw swelling following pIPA_NP_ therapeutic administration, whereas the swelling persisted or worsened in mock PBS-treated animals (**Fig. 4f and Fig. S12**). Paw homogenates from pIPA_NP_-treated mice also contained significantly lower IL-6 concentrations (**Fig. 4e**), similar to what was observed in pIPA_NP_ prophylaxis.

Flow cytometry revealed changes in dendritic cell composition across the spleen, hdLNs, ingLNs, and mesenteric lymph nodes (mLNs). Within the CD11c⁺Ly6G⁻ compartment, pIPA_NP_ treatment significantly increased the proportion of CD11b⁻CD103⁺ cells, and decreased the CD11b⁺CD103⁻ fraction, in all four lymphoid tissues compared with mock PBS treatment (**Fig. 4g**; **gating in Fig. S6**). These DC population shifts were similar to what was observed under pIPA_NP_ prophylaxis and therefore remained evident when treatment was initiated after disease onset. Similarly, changes in costimulation-associated markers were particularly apparent in dendritic cell populations. In hdLNs and mLNs, pIPA_NP_ reduced CD86⁺, CD80⁺, and CD40⁺ frequencies within CD11b⁺CD11c⁺ cells, while CD103⁺CD11c⁺ cells showed reductions in all four measured markers, including MHC-II (**Fig. 4h and Figs. S14–S15**). In ingLNs, CD103⁺CD11c⁺ cells also exhibited lower CD86⁺, CD80⁺, and CD40⁺ frequencies, although the difference in MHC-II positivity did not reach statistical significance (**Fig. S16**). The splenic response was more mixed. CD103⁺CD11c⁺ cells exhibited lower CD86⁺ and CD80⁺ frequencies but increased MHC-II⁺ frequencies following pIPA_NP_ treatment (**Fig. 4h and Fig. S13**). Splenic CD11b⁺F4/80⁺ cells also showed increased MHC-II expression. Across tissues, CD11b⁺CD11c⁻ populations displayed fewer consistent changes in costimulatory-marker frequencies (**Figs. S13–S16**).

### pIPA_NP_ associate preferentially with myeloid populations in lymphoid tissues

The use of DiD-loaded pIPA_NP_ allowed the examination of cellular biodistribution of pIPA_NP_ in the context of arthritis. Here, we analyzed cell-associated DiD fluorescence in lymphoid tissues collected previously(**Fig. 4**). DiD-positive frequencies and mean fluorescence intensity (MFI) were quantified in hdLNs, ingLNs, mLNs, and spleen to assess the fraction of labeled cells and the intensity of fluorescence within each population (**Fig. 5a, b**; myeloid gating in **Fig. S17**).

**Figure 5.**
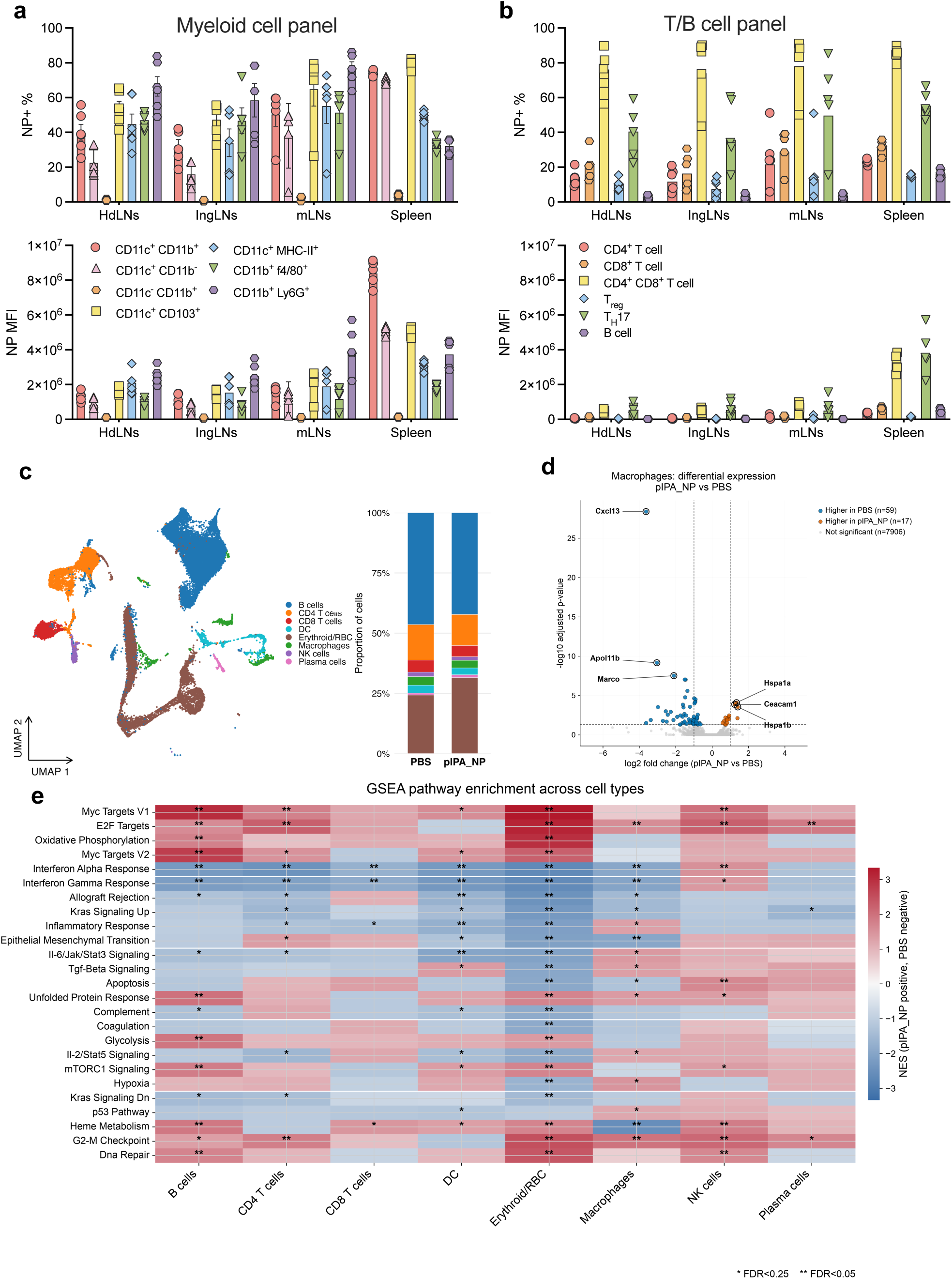
Cellular distribution of pIPA NPs and exploratory splenic transcriptional profiling in CAIA. **a,b.** Cellular distribution of fluorescently labeled pIPA NPs in myeloid populations (**a**) and T- and B-cell populations (**b**) from the therapeutic CAIA study shown in Fig. 4. The percentage of NP-positive cells (top) and mean fluorescence intensity (MFI) of NP-associated fluorescence (bottom) are shown for hock-draining lymph nodes (HdLNs), inguinal lymph nodes (IngLNs), mesenteric lymph nodes (mLNs), and spleen. **c.** UMAP of splenic scRNA-seq data colored by broad cell-type annotation and relative cell-type composition in PBS- and pIPA NP-treated mice (33,616 annotated cells; n=2 per group). Unresolved clusters were excluded. Composition was calculated from cells pooled within each treatment group and is descriptive. **d.** Volcano plot of macrophage differential gene expression using counts aggregated per mouse and analyzed by DESeq2 (Wald test with Benjamini-Hochberg correction). Orange and blue indicate genes with higher expression in pIPA_NP_ and PBS groups, respectively, at adjusted *P* < 0.05; gray indicates nonsignificant genes. Dashed lines indicate adjusted *P* = 0.05 and log₂ fold change = ±1, with the latter shown as reference lines. **e.** Pre-ranked gene set enrichment analysis (GSEA) of selected MSigDB Hallmark pathways across cell types, with genes ranked by the DESeq2 Wald statistic. Color indicates the normalized enrichment score for pIPA NP versus PBS (red, positive; blue, negative). *FDR < 0.25; **FDR < 0.05.

Several myeloid populations exhibited substantial NP-associated fluorescence, including CD11c⁺CD11b⁺, CD11c⁺CD103⁺, CD11c⁺MHC-II⁺, CD11b⁺F4/80⁺, and CD11b⁺Ly6G⁺ cells. CD11c⁺CD103⁺ cells and CD11b⁺Ly6G⁺ neutrophils were among the populations with the highest DiD-positive frequencies in lymph nodes. In contrast, CD11c⁻CD11b⁺ cells showed consistently low DiD-positive frequencies and MFI across the tissues examined (**Fig. 5a**). This limited association coincided with the fewer costimulatory-marker changes observed in this population during pIPA_NP_ therapeutic intervention (**Fig. 4h and Figs. S13–S16**). The distribution of DiD fluorescence showed similar trends among cell populations across tissues. A few exceptions included several CD11c^+^ populations exhibited greater MFI in the spleen than in lymph nodes, with particularly strong fluorescence in splenic CD11c⁺CD11b⁺ cells (**Fig. 5a**).

Conventional CD4⁺ and CD8⁺ T cells, regulatory T cells, and B cells generally exhibited lower DiD-positive frequencies and MFI than the strongly labeled myeloid populations (**Fig. 5b**, **gating in Fig. S18**). However, there are certain T cell populations including CD4⁺CD8⁺ and Th17 T cells also contained an appreciable DiD-positive fraction. Both populations exhibited greater fluorescence intensity in the spleen than in lymph nodes (**Fig. 5b**), demonstrating that association with the fluorescent formulation also occurred in selected T-cell populations.

### Exploratory single-cell profiling reveals distinct transcriptional responses across splenic populations

We collected spleens from additional PBS- and pIPA_NP_-treated arthritic mice at CAIA endpoint following therapeutic administration. To examine transcriptional changes associated with pIPA_NP_ treatment, we performed exploratory single-cell RNA sequencing of splenocytes. After exclusion of unresolved clusters, 33,616 cells were assigned to eight broad populations using canonical marker expression: B cells, plasma cells, CD4^+^ T cells, CD8^+^ T cells, NK cells, macrophages, dendritic cells, and erythroid cells (**Fig. 5c and Fig. S19**). We did not observe significant changes in cell population fractions from pooled cell proportions, indicating that the pIPA_NP_ did not change the major immune cell populations (**Fig. 5c**).

Mouse-level pseudobulk analysis using DESeq2 identified the largest number of differentially expressed genes in macrophages (annotation showed in **Fig. S19**), with 59 genes expressed at lower levels and 17 at higher levels following pIPA_NP_ treatment at adjusted *P* < 0.05 (**Fig. 5d and Fig. S20f**). Lower-expressed genes included *Cxcl13* (log₂ fold change = −3.63), *Apol11b*, and *Marco*, while higher-expressed genes included *Hspa1a*, *Ceacam1*, and *Hspa1b* (**Table S2**). B cells had 25 differentially expressed genes, whereas fewer genes met this threshold in dendritic cells, CD4 T cells, and CD8 T cells, with six, two, and one detected differences, respectively. No genes met the adjusted significance threshold in NK cells (**Fig. S19**).

To assess coordinated changes beyond individually significant genes, preranked gene set enrichment analysis was performed using MSigDB Hallmark gene sets. Both interferon-α and interferon-γ response signatures were negatively enriched in B cells, CD4 and CD8 T cells, dendritic cells, macrophages, and erythroid cells following pIPA_NP_ treatment (FDR < 0.05; **Fig. 5e**). NK cells showed the opposite pattern, with positive enrichment of the interferon-α response signature at FDR < 0.05 and the interferon-γ response signature at the exploratory FDR < 0.25 threshold. Additional pathway changes involved E2F target gene sets, which were positively enriched in B cells, CD4 T cells, macrophages, NK cells, erythroid cells, and plasma cells, and oxidative phosphorylation, which was positively enriched in B cells and erythroid cells (FDR < 0.05; **Fig. 5e**).

IL-6-JAK-STAT3 signaling signatures were negatively enriched in B cells, CD4⁺ T cells, and DCs, while inflammatory-response signatures were negatively enriched in CD4⁺ and CD8⁺ T cells and DCs. In macrophages, however, both signatures were positively enriched at the exploratory FDR < 0.25 threshold, despite negative enrichment of interferon-response signatures (**Fig. 5e**).

### Local pIPA_NP_ treatment reduces established arthritis and reproduces key immune phenotypes

Finally, rather than pIPA_NP_ subcutaneous administration near the flank, we injected pIPA_NP_ near the paws to model local treatment injections performed in rheumatoid arthritis patients. Mice with established CAIA received subcutaneous injections of PBS, IPA_free_, or pIPA_NP_ near all four hocks at 2 mg per mouse on days 6 and 7 following disease induction (**Fig. 6a**). Clinical arthritis scores declined progressively in the pIPA_NP_ group, while disease activity remained elevated in the PBS and IPA_free_ groups. The post-treatment cumulative arthritis score was significantly lower with pIPA_NP_ than with PBS, although the reduction relative to IPA_free_ did not reach statistical significance (**Fig. 6b**). pIPA_NP_-treated mice also showed a more complete recovery of body weight during follow-up (**Fig. 6c**). Average paw thickness decreased significantly between days 6 and 14 in the pIPA_NP_ group, whereas the paired changes in the PBS and IPA_free_ groups were not significant (**Fig. 6d**).

**Figure 6.**
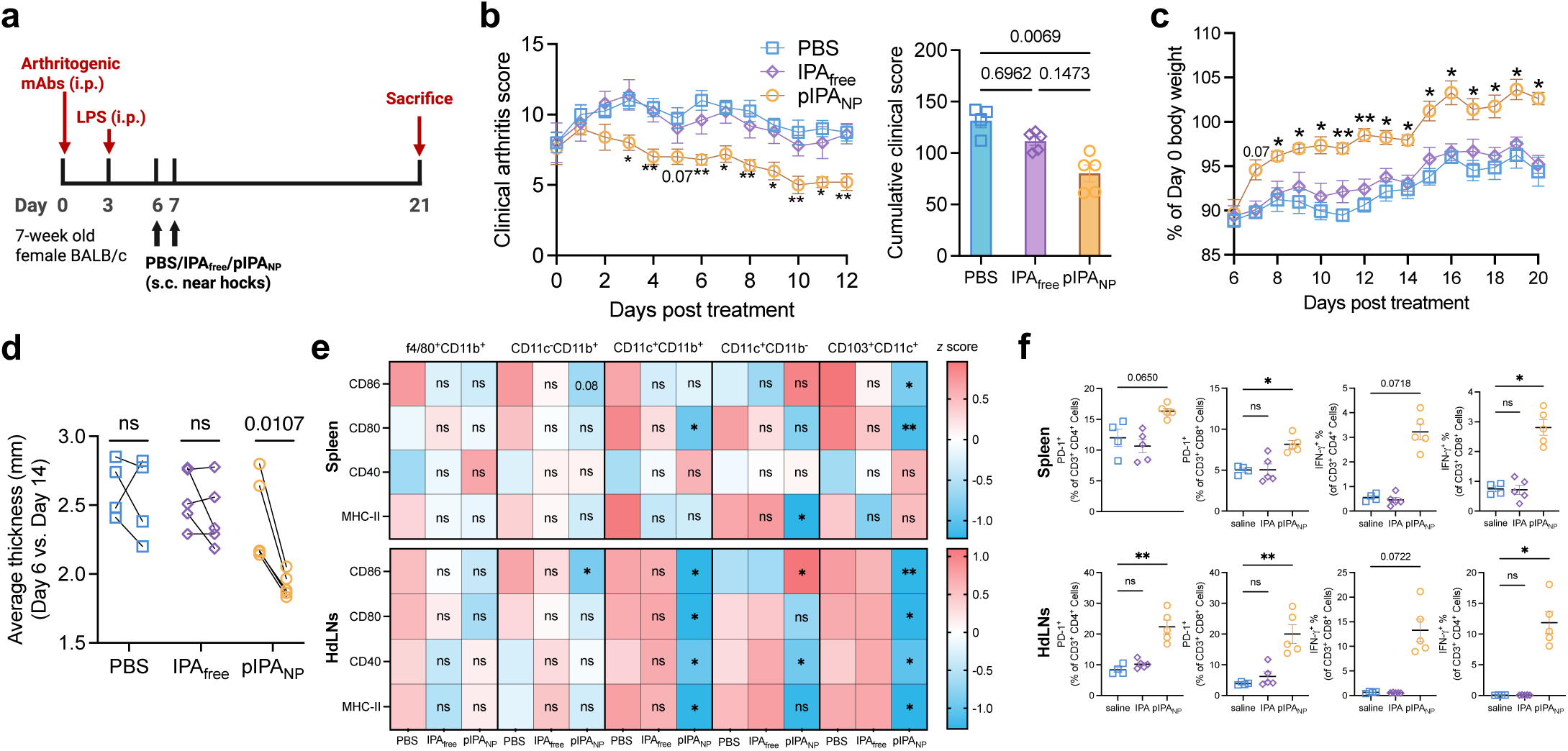
Subcutaneous injection of pIPA_NP_ near the hocks recapitulates therapeutic efficacy while inducing local and systemic immunomodulation. a. Experimental timeline. b. Clinical arthritis scores of mice from Day 0 to Day 12 post-treatment and the average post-treatment cumulative arthritis score of each subject (n=4 for PBS group, n=5 for IPA_free_ and pIPA_NP_ group). c. Daily body weight expressed as a percentage of Day 0 body weight from day 0 to day 12 post-treatment. d. Average thickness of all paws at Day 6 vs. Day 14. Paired t test was used for statistical analysis. e. Heat map of z-score-normalized % of CD86^+^, CD80^+^, CD40^+^, and MHC-II^+^ in CD11b^+^f4/80^+^, CD11b^+^CD11c^-^, CD11b^+^CD11c^+^, CD11c^+^CD11b^-^, and CD103^+^CD11c^+^ cells in spleen and hock-draining lymph nodes (hdLNs) (*z*=(x−mean)/SD). f. Percentage of PD-1^+^ and IFN-γ_+_ cells among CD4+ and CD8+ cells in spleen and hdLNs. Data are presented as mean ± SEM. Statistical comparisons were performed using a paired t test (d, [other panels]) or the Kruskal-Wallis test. *p < 0.05, **p < 0.01, ***p < 0.001, ****p < 0.0001; ns, not significant.

Representative forepaw and hindpaw showed early reductions in visible swelling between days 6 and 8 in pIPA_NP_-treated animals, with variation in the extent of improvement among animals (**Figs. S20 and S21**).

In hdLNs collected at the study endpoint, pIPA_NP_ treatment reduced the frequencies of CD86⁺, CD80⁺, CD40⁺, and MHC-II⁺ cells within both CD11b⁺CD11c⁺ and CD103⁺CD11c⁺ populations compared with mock PBS treatment (**Fig. 6e**). These findings were consistent with the prominent reductions in hdLN costimulatory-marker frequencies observed in the earlier prophylactic and therapeutic study. Splenic changes were more selective, including lower CD80⁺ frequencies within CD11b⁺CD11c⁺ cells, lower CD86⁺ and CD80⁺ frequencies within CD103⁺CD11c⁺ cells, and lower MHC-II⁺ frequencies within CD11b⁻CD11c⁺ cells. Other hdLN subsets showed marker-specific responses: CD86⁺ frequencies decreased within CD11b⁺CD11c⁻ cells, whereas CD11b⁻CD11c⁺ cells showed increased CD86⁺ and decreased CD40⁺ frequencies. In the T-cell compartment, pIPA_NP_ treatment increased PD-1⁺ frequencies among both CD4⁺ and CD8⁺ T cells in hdLNs and among CD8⁺ T cells in the spleen compared with PBS (**Fig. 6f**). IFN-γ⁺ frequencies also increased among CD8⁺ T cells in both tissues. Splenic PD-1⁺ CD4⁺ T-cell frequencies and IFN-γ⁺ CD4⁺ T-cell frequencies in both tissues were numerically higher with pIPA_NP_, but these differences did not reach statistical significance.

## 4. Discussion

Microbiota-derived metabolites offer opportunities for immune modulation, but translating their biological activity into treatment requires control over their exposure in relevant tissue. Here, we developed a polymeric IPA nano-prodrug formulation that sustained measurable IPA exposure in lymph nodes and reduced arthritis severity when administered both before or after disease onset. We observed immunological changes in dendritic cell costimulatory marker frequencies, relative DC subset representation, and T cell PD-1 and IFNγ phenotypes. These findings support further development of pIPA_NP_ as an injectable approach for delivering a microbial metabolite to immune compartments involved in inflammation.

Following a single subcutaneous administration, DiR-labeled pIPA_NP_ produced detectable fluorescence for at least 35 days, while LC-MS/MS measurements demonstrated lymph node IPA exposure throughout the 14-day sampling period (**Fig. 1**). Nanoparticle access to lymph nodes is consistent with previous studies showing that particle properties influence lymphatic transport and exposure of lymph-node antigen-presenting cells (*20*).

However, longer pharmacokinetic and efficacy studies will therefore be needed to define the relationship between carrier retention, IPA exposure, and duration of disease control. IPA’s microbial origin provides a biological rationale and the formulation allows this existing host-microbe signal to be administered at a defined dose with a controlled pattern of tissue exposure.

The PBMC incubation study provides an initial indication that cellular processes may contribute to IPA release from pIPA_NP_, especially given our observation that no IPA release was detected in cell-free media (**Fig. 1m**). However, this short incubation does not define the longer-term release kinetics or identify the responsible cell population. Release could involve extracellular hydrolysis, intracellular processing after particle uptake, or both. Whether esterases or other enzymes participate remains unresolved. Studies in purified immune populations, combined with measurements of extracellular and intracellular IPA and polymer-associated material, will help distinguish these possibilities. Particularly, the cellular biodistribution findings identify DC-containing populations as candidates for further mechanistic investigation. Nanoparticle-associated fluorescence was generally greater in myeloid populations than in conventional T- and B-cell populations, and prominent reductions in costimulatory marker frequencies occurred within CD11b⁺CD11c⁺ and CD103⁺CD11c⁺ populations in hdLNs (**Figs. 3–6**). This overlap supports the hypothesis that exposure of antigen-presenting cells contributes to the immune response to pIPA_NP_. Nevertheless, fluorescence association does not establish internalization, and lower frequencies of marker-positive cells do not directly measure antigen presentation or the capacity to activate T cells. This might require further studies such as functional DC–T-cell coculture studies to connect these phenotypes with immune responsiveness.

Which populations are necessary for disease protection remains an open question. The increased relative representation of CD11b⁻CD103⁺ cells in both prophylactic and therapeutic studies is of interest. Conventional DC subsets have distinct functional specializations: cDC1s are associated with efficient antigen cross-presentation, while cDC2 populations can support CD4⁺ T-cell differentiation, including Th17 responses demonstrated in mucosal settings (*25*, *26*). A change in their relative representation could therefore alter the balance of signals available to T cells. However, cDC1 enrichment cannot by itself be interpreted as induction of tolerance. Recent work demonstrated that erythropoietin-receptor signaling can promote a tolerogenic cDC1 state, whereas another study found that Batf3-dependent cDC1s were required for initiation of CIA (*27*, *28*).

CD103⁺ DCs can include tissue-derived migratory populations (*29*, *30*), raising questions about their contribution to particle transport and immune modulation.

Changes in T cell PD-1 expression provide another possible connection between lymphoid exposure and disease protection. PD-1 can restrain T cell responses when its inhibitory signaling pathway is engaged, and a phase 2a trial of the PD-1 agonist peresolimab demonstrated activity in patients with RA (*31*). However, increased PD-1 expression alone does not establish enhanced inhibitory signaling or T cell exhaustion. Petrelli et al. showed that PD-1⁺ CD8⁺ T cells in inflammatory arthritis could retain proliferative and effector functions, including IFNγ production (*23*). Our findings may therefore reflect activation-associated induction of an inhibitory receptor, potentially increasing the opportunity for feedback regulation. Defining PD-1 and IFN-γ coexpression, correlation, and responsiveness to PD-1 engagement would help determine the functional significance of these changes. The increased frequencies of IFN-γ⁺ T cells accompanying reduced arthritis severity require a similarly contextual interpretation. Although IFN-γ can participate in inflammatory responses, its role in experimental arthritis depends on the model and immune setting. In CIA, endogenous IFN-γ was shown to limit disease susceptibility through suppression of IL-17 responses (*24*). This provides a possible explanation for the association between increased IFN-γ⁺ frequencies and disease protection, but an IFN-γ–IL-17 mechanism has not been demonstrated in our CAIA experiments. Increased IFN-γ positivity among CD4⁺ T cells also raises the possibility of a Th1-associated response, which would require confirmation through T-bet expression and broader functional profiling.

Recent studies on IPA have demonstrated that its immune effects can differ across cellular, tissue, and disease contexts. Li et al. showed that IPA enhanced mitochondrial respiration in CD4⁺ T cells and conferred protection against intestinal inflammation (*15*). Fernandes et al. reported that IPA derived from helminth-altered microbiota promoted lung epithelial type I interferon responses and protected offspring against respiratory viral infections (*32*), although the epithelial type I interferon mechanism does not directly explain the increased T-cell IFN-γ frequencies we observed in lymphoid tissues. Differences in formulation, exposure, cell population, and inflammatory environment may contribute to these distinct responses.

Future translation will require extending these findings beyond the acute antibody-induced mouse model. CAIA can develop without adaptive immune priming, although collagen-reactive T cells can influence disease progression (*22*). Evaluation in CIA, including treatment after onset and longer follow-up, would allow assessment of adaptive immune responses and sustained joint protection. The hock injection study also supports further investigation of injection-site selection, with comparisons at matched doses and volumes to determine how administration near inflamed joints affects drug exposure and therapeutic efficacy. These studies could help establish whether local administration of this long-acting platform offers a clinically relevant approach to sustained disease control. Clinical translation will also require evaluation of repeat-dose tolerability, injection-site reactions, and polymer clearance, together with functional assessments of vaccine responses and antimicrobial host defense. Additionally, physiological exposure to IPA does not establish the safety of sustained pharmacological exposure or of the synthetic polymer. A pharmacokinetic framework that distinguishes released IPA, polymer-associated IPA, and residual carrier would support selection of dosing intervals while accounting for both therapeutic exposure and material persistence.

## 5. Materials and Methods

### Synthesis of [2-(methacryloylamino)ethyl]indole-3-propionate monomer (MA-IPA)

To synthesize the MA-IPA monomer, 3-IPA (10 g, 52.9 mmol), DMAP (0.65 g, 5.3 mmol), and 60 mL of DCM were added to a 200 mL vacuum-dried flask. The mixture was cooled over ice for 15 minutes, and a solution of DCC (12 g, 58.2 mmol) in a minimum volume of DCM was added. Subsequently, HEMAm (7.5 g, 58.2 mmol) was added dropwise, and the reaction was proceeded with stirring at room temperature overnight. The reaction mixture was filtered and sequentially washed with saturated ammonium chloride, sodium bicarbonate, and brine solution (2x each) before being dried with anhydrous magnesium sulfate. The reaction mixture was then concentrated via rotary evaporation and purified by silica gel flash chromatography on Teledyne ISCO, CombiFlash^®^ NextGen 300+ system using a gradient of CHCl_3_/MeOH (MeOH fraction v/v, 0-5%). The product was obtained as a pale yellow solid (13.6g, 86% yield). 1 H NMR (500 MHz, DMSO-d 6 , δ, ppm): 1.85 (s, 3H, −CCH 3 ), 2.67 (t, 2H, −CH 2 C=O), 2.96 (t, 2H, −CH 2 NH), 3.36 (t, 2H, C 8 H 5 NH-CH 2 ), 4.08(t, 2H, −OCH 2 ), 5.35 and 5.65 (s, 2H, −C=CH 2 ), 6.96–7.52 (m, 5H, indole ring), 8.03 (s, 1H, −NHC=O), 10.79 (s, 1H, −NHCH) (**Fig. S1**).

### Synthesis of poly([2-(methacryloylamino)ethyl]indole-3-propionate) (pIPA)

The pIPA was prepared using 2-cyano-2-propyl benzodithioate (CPBDT) as the RAFT chain transfer agent at 70 °C. Briefly, MA-IPA (1.5 g, 5.0 mmol), CPBDT (14.2mg, 0.067 mmol), and AIBN (3.2 mg, 0.02 mmol) were added to a dry glass vial and dissolved in 5 mL of DMF. The reaction mixture was purged with nitrogen for 30 minutes and then allowed to polymerize at 70 °C for 16 h. After polymerization, the crude polymer was purified by precipitation into petroleum ether/methanol (1:1, v/v) three times, redissolving in acetone between each cycle. The resulting solid was dried under vacuum overnight. The pIPA was obtained as a pink solid. 1 H NMR (500 MHz, DMSO-d_6_ , δ, ppm): 0.87-1.14 (m, 3H, backbone CH 3 ), 1.67 (m, 2H, backbone CH 2 ), 2.64 (m, 2H, CH 2 C=O), 2.93(m, 2H, CH 2 NH), 3.18(m, 2H, C 8 H 5 NH-CH 2 ), 3.97(m, 2H, OCH 2 ), 6.91-7.45(m, 5H, indole ring), 7.96(m, 1H, NHC=O), 10.76(m, 1H, NHCH) (**Fig. S2**).

### Formulation of pIPA Nanoparticles (pIPA_NP_)

The pIPA_NP_ was synthesized from pIPA using the single emulsion solvent evaporation method. A 50 mg/mL solution of pIPA was prepared in ethyl acetate, and 1 mL of this solution was added dropwise to 2 mL of 5% (w/v) polyvinyl alcohol aqueous solution while vortexing in a glass tube. The mixture was then probe-sonicated for 20 seconds at 50% amplitude using a Fisherbrand CL-18 system coupled with an FB120 ultrasonic processor (120 W, 20 kHz; Fisher Scientific), followed by 10 seconds of vortexing. This coupled process was repeated three times. The emulsified mixture was transferred to 40 mL of ultrapure water in a round-bottom flask and depleted of ethyl acetate by rotary evaporation. The suspension was then filtered with Millipore^®^ Steriflip^®^ Vacuum Tube Top Filter (50 mL, 0.22 um) to ensure sterility for *in vivo* administration. The filtered content was divided into two 50-mL centrifuge tubes and centrifuged at 20,000 ×g for 40 min. Following centrifugation, the supernatant was carefully discarded. The resulting pellets containing pIPA-NP were resuspended in ultrapure water.

### Dynamic Light Scattering (DLS) and Zeta Potential

The hydrodynamic diameter and polydispersity index (PDI) of pIPA_NP_ were measured by DLS at 25°C (Brookhaven Instruments). 10 µL of pIPA_NP_ suspension was diluted in 2 mL of ultrapure water in a disposable cuvette. Zeta potential was measured at 25°C (Malvern Panalytical) using disposable capillary zeta cells, with samples diluted 1:150 in 10 mM sodium chloride solution. Samples were equilibrated for 90 s before analysis, and each measurement was performed in triplicate, with an acquisition time of 60 s per replicate.

### Transmission Electron Microscopy (TEM)

Carbon-coated copper grids (CF200-Cu-50, Electron Microscopy Sciences) were plasma glow-discharged (O_2_/Ar, 30 s) before sample deposition. pIPA_NP_ suspensions (40 µL) were incubated on the grids for 2 min, followed by removal of excess liquid using Whatman filter paper. Grids were negatively stained with UranyLess for 1 min, blotted dry, and air-dried for 15–30 min before imaging. TEM images were acquired using a FEI Tecnai G2 F20 S-Twin TEM operated at 200 kV.

### Liquid Chromatography-Tandem Mass Spectrometry (LC-MS/MS)

Standards of IPA (Sigma, Cat#106-31-0) were prepared in 1:1 acetonitrile-water mixture and diluted to create standard curves ranging from 1-1000 ng/mL. The internal standard used was indole-2,4,5,6,7-d5-3-acetic Acid (IAA-d5) (CDN Isotopes, Cat#D-2203), and 5 ng of the IAA-d5 was spiked into each sample. The LC-MS/MS analysis was performed on an ACQUITY UPLC system coupled with a Waters Xevo TQ-S mass spectrometer in positive electrospray ionization mode. Quantification was carried out using multiple reaction monitoring (MRM) transitions at m/z 190.18 > 130.12 for 3-IPA and m/z 181.23 > 133.92 for IAA-d5. The cone voltage was optimized at 64 V for the 3-IPA transition and 4 V for the IAA-d5 transition. Collision energies were optimized at 18 V for 3-IPA and 16 V for IAA-d5. The analysis used an Ascentis^®^ Express 90 Å RP-Amide HPLC Columns (15 cm × 2.1 mm, 2.7 μm), maintained at 30°C. The mobile phase consisted of 0.4% formic acid in water (A) or in acetonitrile (B), with a flow rate of 0.3 mL/min, a total run time of 9 minutes, and an injection volume of 10 µL. To quantify the IPA content of pIPA_NP_, samples were diluted 1:100 in 1 M NaOH and hydrolyzed overnight following 1 h of bath sonication.

### Plasma sample preparation

Plasma samples (20 µL) were extracted with 100 µL of ice-cold acetonitrile containing IAA-d_5_ as an internal standard. Samples were vortexed for 30 seconds, incubated at 4 °C for 15 min, and centrifuged at 18,000 × g for 15 min at 4 °C. The supernatant was diluted with water to a final composition of 50% water and filtered through 0.2 µm syringe filters prior to LC-MS analysis.

### Lymph node samples preparation

Saline (100 µL) was added to pre-weighed lymph node samples, which were then homogenized using an Omni Bead Ruptor Elite (Revvity). The tissue homogenate was extracted with 300 µL of ice-cold acetonitrile, vortexed for 30 seconds, incubated at 4 °C for 15 min to allow efficient extraction, and centrifuged at 18,000 × g for 15 min at 4 °C. The supernatant was diluted with water to a final composition of 50% water and filtered through 0.2 µm syringe filters prior to LC-MS analysis.

### Mice

All procedures involving animals were approved by the Institutional Animal Care and Use Committee (IACUC) at the University of Washington (Protocol #4581-01), and adhered to the Animal Welfare Act, the Guide for the Care and Use of Laboratory Animals, and Public Health Service Policy on Humane Care and Use of Laboratory Animals. BALB/c mice, aged 8-12 weeks, were purchased from The Jackson Laboratory and used in this study. Mice were housed in a specific pathogen-free facility at the University of Washington and maintained on a 12-hour light/dark cycle at 20°C to 26°C.

### Biodistribution of pIPA-NP

Eight-week-old female BALB/c mice (The Jackson Laboratory) were used to study *in vivo* biodistribution of pIPA_NP_. 1% (w/w) of 1,1′-dioctadecyl-3,3,3′,3′-tetramethylindotricarbocyanine iodide (DiR) dye was mixed with a 50 mg/mL IPA polymer stock and synthesized into DiR-pIPA-NP following the same procedure described in the *Formulation of pIPA_NP_* section. The DiR-pIPA_NP_, PBS, or free DiR was injected subcutaneously around the mice’s abdominal area. Live imaging was taken on days 0, 1, 3, 7, 14 for the free DiR group and on days 0, 1, 3, 7, 14, 21, 28, and 35, for the PBS and DiR-pIPA_NP_ group, using IVIS^®^ Spectrum Imaging System (XENOGEN^®^). On day 15 (free DiR) or 36 (PBS and DiR-pIPA-NP), mice were euthanized, and their tissues and lymph nodes were collected for *ex vivo* imaging.

### Human blood sample collection

Fresh whole blood was drawn from an adult patient with RA. Study participants were recruited through the University of Washington, Division of Rheumatology Biorepository to participate in research studies at the University of Washington Medical Center, Seattle, WA. The study was approved by the institutional ethics board (STUDY00006196), and informed written consent was obtained from all participants according to the Declaration of Helsinki.

### Human PBMC isolation

Polymorphonuclear (PMN) cells and peripheral blood mononuclear cells (PBMC) were isolated from freshly collected whole blood using PolymorphPrep (Cosmo Bio, USA) according to the manufacturer’s instructions. Briefly, whole blood was layered onto an equal volume of PolymorphPrep and centrifuged at 625 x g for 30 minutes without stopping the brake. PBMC and PMN layers were transferred to individual tubes. Red blood cells were removed with RBC lysis buffer (BioLegend, USA). Cells were then washed in 1× PBS and counted on a hemocytometer before downstream application.

### pIPA_NP_ Release in Human PBMCs

pIPA_NP_ at various concentrations was incubated with human PBMCs in serum-free RPMI 1640 medium for 4 h at 37°C and 5% CO₂. Cells were pelleted by centrifugation at 300 × g for 5 min. The supernatant was collected, and the cell pellet was lysed in 100% acetonitrile (ACN). The lysate was then combined with the supernatant. Negative controls consisted of pIPA_NP_ incubated in cell-free RPMI under identical conditions. IPA concentrations were quantified by LC-MS.

### Collagen-Antibody Induced Arthritis (CAIA) Model

To induce CAIA, eight-week-old female BALB/c mice (The Jackson Laboratory) were injected intraperitoneally with 1 mg of anti-collagen antibody cocktail (Chondrex, Inc., Cat#53040) on Day 0, followed by 25 μg of LPS (Chondrex, Inc.) on Day 3. Starting from Day 3, clinical arthritis score and body weights were recorded daily until the end of the study by one person who was blinded to the group assignment. The scoring criteria are detailed in **Supplementary Table 1**. Mice were euthanized at the study endpoints, and their spleens, inguinal, popliteal, axillary, and brachial lymph nodes were collected and processed into fresh single-cell suspensions for flow cytometry. The right paws were homogenized for cytokine analysis, while the left paws were fixed in formalin for histology.

### Prophylactic Treatment Study in the CAIA Model

For the prophylactic CAIA study, mice received subcutaneous injections of PBS, pIPA_NP_, or free IPA (IPA_free_) on Day -7 and Day -1 before disease induction. The dosing schedule was selected to establish IPA exposure before CAIA induction. The concentration of IPA contained within pIPA-NP formulations was quantified by LC-MS/MS and adjusted to 10 mg/mL to match the IPA concentration of the free IPA control formulation. Each mouse received an equivalent IPA dose of 2 mg per injection (10 mg/mL IPA, as in IPA_free_ or pIPA_NP_, in 200 μL PBS). CAIA was induced according to the protocol described above. Body weight and clinical arthritis scores were monitored daily from Day 3 until study termination. Paw thickness was measured using a digital caliper on Day 0 and Day 11 to evaluate disease progression and treatment efficacy.

### Therapeutic Treatment Study in the CAIA Model

For the therapeutic CAIA study, disease progression was monitored daily following CAIA induction. Treatment was initiated once the average clinical arthritis score reached >5, corresponding to mild swelling across all paws or severe swelling in multiple paws, indicating established disease onset. Mice were grouped to ensure comparable clinical arthritis scores between treatment groups before therapeutic dosing without redistribution of mice between cages. Mice that did not develop arthritis were excluded from the study. For the Figure 4 study, mice were injected subcutaneously in the abdomen with 200 µL of PBS or pIPA_NP_ (10 mg/mL IPA) on Days 6 and 7. In the **Fig. S11** study, mice were dosed with pIPA_NP_ twice at 10 mg/mL IPA (1x) on Days 6 and 7 or once at 20 mg/mL IPA (2x) on Day 6. Body weight and clinical arthritis scores were recorded daily from Day 3 until the study endpoint (Day 13). Paw thickness measurements were obtained using a digital caliper on Day 0, Day 6, Day 8 and Day 13 to monitor arthritis progression and therapeutic response. Representative photographs of paws were taken on Day 6, Day 8, and Day 13 for evaluation of disease severity and treatment-induced changes.

### Hock injection therapeutic CAIA study

Disease induction, grouping, exclusion criteria and monitoring were as described for the therapeutic CAIA study above. Hock subcutaneous injections were performed as previously described (*19*). Mice received a single treatment on Day 6, consisting of 25 µL of PBS, free IPA or pIPA_NP_ injected subcutaneously at each of four sites around the hocks (100 µL total; 20 mg/mL IPA for free IPA and pIPANP). Body weight and clinical arthritis scores were recorded daily, paw thickness was measured on Days 0, 6, 14 and 21, and mice were euthanized on Day 21.

### Flow Cytometry and Antibodies

Flow cytometry was done using Agilent NovoCyte Flow Cytometer Systems, and data were analyzed in NovoExpress Software. Single-cell suspensions of each subject were plated into the corresponding well of a 96-well plate and were pelleted via centrifugation. Dead cells were stained using a Live/Dead Aqua viability dye (1:500 in PBS) and incubated for 13 min at room temperature, followed by anti-CD16/32 (1:100 in PBS) for 13 min. Cells were then stained with surface antibody cocktails for T cell or myeloid panels and incubated for 30 min at 4°C in the dark. After washing, cells were fixed and permeabilized with Foxp3/Transcription Factor Staining Buffer Set (Invitrogen, Cat#50-112-8857) overnight at 4 °C according to manufacturer’s instructions.

Cells were stained the next day with intracellular markers for 1 h at room temperature. Fluorescence minus one (FMO) controls, dead cell controls, and unstained controls were included for both spleen and lymph nodes. Cells were washed and resuspended in 2% FBS in PBS buffer before running flow cytometry.

### Flow cytometry antibodies

Antibodies against the following markers were used in the CAIA studies: CD3 (BUV737, Cat#741788, BD Biosciences), CD4 (BV786, Cat#740844, BD Biosciences), CD8 (BV421, Cat#100730, BioLegend), CD11b (BV711, Cat#101242, BioLegend), CD11c (BV421, Cat#562782, BD Biosciences), CD25 (APC, Cat#162105, BioLegend), CD40 (APC, Cat#558695, BD Biosciences), CD80 (BUV737, Cat#612773, BD Biosciences), CD86 (FITC, Cat#105006, BioLegend), CD206 (PE, Cat# 141706, BioLegend), Ly6G (BV605, Cat# 127639, BioLegend), F4/80 (BUV563, Cat# 365-4801-82, Thermo Fisher Scientific), I-A/I-E (APC/Cy7, Cat# 107628, BioLegend), PD-1 (BV711, Cat#135231, BioLegend), CTLA-4 (PE-Cy7, Cat#25–1522-80, Invitrogen), IFN-*γ* (BV 605, Cat#505840, BioLegend), IL-10 (PE-Cy7, Cat# 505036, BioLegend), Foxp3(FITC, Cat# 50-112-8844, Fisher Scientific), RORγt(PE, Cat#562607, BD Biosciences).

### Paw Homogenization

Mouse paws were cut into small fragments and transferred into 2-mL homogenization tubes containing 1.4 mm zirconium-silicate beads and 800 µL of T-PER protein extraction buffer supplemented with protease inhibitor. Tissue homogenization was performed using a Revvity Omni Bead Ruptor Elite according to the manufacturer’s instructions. The resulting homogenates were first centrifuged at 2,000 × g for 10 min at 4°C to remove large tissue debris. The supernatant was subsequently transferred to 1.5-mL microcentrifuge tubes and centrifuged at 14,000 × g for 15 min at 4°C. Following centrifugation, the upper lipid layer and bottom cellular pellet were carefully avoided, and the clarified middle aqueous layer was collected, aliquoted, and stored at −80°C until further analysis. This protocol is adapted from (*33*).

### Histology

Harvested paws were fixed in 10% formalin, followed by decalcification in Leica Biosystems Decalcifier II for 72 hours. After decalcification, samples were then rinsed with PBS and transferred to 70% ethanol prior to submission to the histology core facility. Paraffin embedding, sectioning, and staining with hematoxylin and eosin (H&E) were performed by the histology core facility at the University of Washington School of Medicine Comparative Medicine Histology and Imaging Center.

### Enzyme-Linked Immunosorbent Assay (ELISA)

Levels of IL-6 in paw homogenates were quantified using mouse uncoated ELISA kits (Invitrogen, Mouse IL-6 ELISA, Cat. No. 88-7064) according to the manufacturer’s instructions. Briefly, 96-well high-binding flat-bottom ELISA plates were coated overnight at 4°C with capture antibody diluted in coating buffer. Plates were washed with PBS containing 0.05% Tween-20, blocked with ELISA diluent for 1 h at room temperature, and incubated with standards (1-500 pg/mL) and diluted samples for 2 h at room temperature or overnight at 4°C. Following incubation, plates were washed and incubated sequentially with avidin-HRP. Plates were then developed using TMB substrate solution, and the reaction was stopped with 1 N HCl. Absorbance was measured at 450 nm with background correction at 570 nm.

### Single-cell RNA Sequencing (scRNAseq)

#### Single-cell suspension and sequencing

Freshly harvested spleens from four mice were sent to Novogene (Sacramento, CA, USA) for single-cell RNA sequencing. Tissue dissociation, single-cell library preparation using the 10x Genomics Chromium GEM-X Single Cell 3′ Gene Expression kit (v4), and sequencing were performed by Novogene according to their standard protocols. Libraries were sequenced on an Illumina NovaSeq X Plus (paired-end 150 bp), targeting 10,000 cells per sample at approximately 20,000 reads per cell. Reads were aligned to the mouse reference genome (GRCm39), and gene expression matrices were generated using the Cell Ranger pipeline (10x Genomics). In total, 38,972 cells were detected across the four samples prior to quality filtering.

#### Quality control, normalization and integration

Cells with fewer than 200 or more than 6,000 detected genes, or with more than 15% of counts from mitochondrial genes (genes prefixed "mt-"), were removed. Genes detected in fewer than three cells were removed. Counts were normalized to 10,000 counts per cell and log1p-transformed. The 2,000 most variable genes were selected with sample as the batch key, and the data were scaled and reduced by principal component analysis. Samples were integrated with Harmony (harmonypy v2.0.0) in Scanpy v1.11.5 (Python 3.11.15, anndata v0.12.19). Cells were additionally scored against the ImmGen reference (SingleR, singler v0.5.0; celldex v0.3.0, 2024-02-26 release, fine labels), and cells whose SingleR delta score fell in the lowest 5% were excluded as low-confidence assignments. SingleR labels were used for this filter only and not to define any cell type reported here. No separate doublet-detection step was applied.

#### Clustering and dimensionality reduction

Clustering and visualization used Seurat v5.5.1 (SeuratObject v5.4.0; R 4.5.0). The data were log-normalized (scale factor 10,000), the 2,000 most variable features were selected, and the data were scaled and reduced to 30 principal components. Batch effects between samples were corrected with Harmony v2.0.5 (group.by.vars = "sample", theta = 2; converged after 9 iterations). A shared-nearest-neighbor graph was built on the 30 Harmony dimensions, and cells were clustered by the Louvain algorithm at resolution 1.0, giving 35 clusters across 36,141 cells. UMAP embeddings were computed on the same 30 dimensions (cosine metric, uwot).

#### Cell-type annotation

Cell types were assigned manually, cluster by cluster, from the expression of canonical lineage markers (Supplementary Table 1). Clusters were assigned to B cells, CD4 T cells, CD8 T cells, NK cells, macrophages, dendritic cells (DC), erythroid cells, or plasma cells. Four clusters (13, 17, 19 and 24; 2,525 cells, 7.0%) lacked a coherent lineage signature and were labeled "Unresolved". They are omitted from the UMAP and composition displays and were not tested in downstream analyses. Cluster annotations from Seurat were transferred to the Scanpy object by cell barcode. Pseudobulk differential expression, GSEA, GO and LIANA analyses used the Scanpy object (n = 31,399 cells).

#### Cell composition

For each condition, cells from both samples were pooled, and cell-type proportions were calculated as the fraction of annotated cells (non-annotated cells are excluded; 33,616 cells in the Seurat object). Composition is descriptive; no statistical test was applied.

#### Pseudobulk differential expression

Raw counts were summed per sample within each cell type to generate pseudobulk profiles. A cell type was tested only if it had at least 30 cells overall, and a sample–cell-type profile was included only if it contained at least 5 cells. Genes with fewer than 10 total counts across the retained profiles were removed. Differential expression between pIPA_NP and PBS was tested with PyDESeq2 v0.5.4 using the design ∼ condition and the contrast (condition, pIPA_NP, PBS). Statistical significance was assessed with the Wald test. Cook’s-distance outlier filtering and independent filtering were left at their defaults (on), and p-values were adjusted by the Benjamini–Hochberg procedure. Log2 fold changes are unshrunken maximum-likelihood estimates. Genes with adjusted P < 0.05 were called differentially expressed (DEGs). With n = 2 per condition, these tests have limited power and should be interpreted accordingly.

For the DE heatmap (Fig. 2), up to 10 DEGs (adjusted P < 0.05) per cell type with the lowest adjusted P (ties broken by |log2FC|) were selected, and the union of these genes (38 unique genes across 46 cell-type–gene pairs) was displayed. NK cells had no DEGs and are shown without selected genes. Panel A shows log2FC (*adjusted P < 0.05; ** adjusted P < 0.01), and black boxes mark the cell type for which a gene was selected. Panel B shows mean log-normalized expression, averaged per sample and then per condition, z-scored per gene and clipped to ±2.5. Genes were hierarchically clustered by average linkage on correlation distance.

#### Gene set enrichment analysis (GSEA)

Genes were ranked by the DESeq2 Wald statistic, and preranked GSEA was run with gseapy v1.3.1 against MSigDB Hallmark 2020 (minimum gene-set size 10, maximum 500, 1,000 permutations, seed 0). Mouse gene symbols were converted to upper case to match the human gene-set symbols, and duplicate symbols were collapsed by retaining the entry with the largest |statistic|. Pathways significant (FDR < 0.25) in most cell types were displayed as normalized enrichment scores (NES) (Fig. 4). * marks FDR < 0.25 and ** marks FDR < 0.05. Pathways for display (Fig. 4) were chosen from Hallmark sets whose names matched a pre-specified list of immune, stress-response, metabolic and proliferation terms (interferon, TNFα, inflammatory, IL-6/JAK/STAT, complement, apoptosis, p53, unfolded protein, hypoxia, glycolysis, oxidative phosphorylation, mTORC1, allograft rejection, coagulation, reactive oxygen species, KRAS, TGF-β, epithelial–mesenchymal, MYC, E2F, G2M, DNA repair and UV response), ranked by the lowest FDR in any cell type, then the largest absolute NES; the 25 top-ranked pathways were displayed.

#### Gene Ontology over-representation analysis

Up- and down-regulated genes in each cell type were analyzed separately with Enrichr (gseapy v1.3.1; GO_Biological_Process_2023, mouse). Where a direction had at least 10 genes with adjusted P < 0.05, that list was used. For the 13 of the 16 direction-specific lists (8 cell types × 2 directions) that did not meet this criterion, the 200 genes with the lowest adjusted P in that direction were used instead, and these results are exploratory. Terms with FDR ≤ 0.25, at least three overlapping genes and presence in at least two cell types were retained. The top 25 terms per direction were plotted (**Fig. 5**).

#### Cell-cell communication analysis

Ligand–receptor interactions were inferred with LIANA v1.8.1 for mouse consensus database separately for each condition. Only cell types with at least 100 cells in both conditions were included. Interactions detected in both conditions (6,578) were compared, and those with a magnitude rank ≥ 0.99 in either condition were removed (4,700 retained). For each interaction, the rank-enrichment score was defined as magnitude rank (PBS) − magnitude rank (pIPA_NP), so positive values indicate stronger inferred signaling in pIPA_NP; 2,173 interactions were enriched in pIPA_NP and 2,527 in PBS. Chord diagrams (Fig. 6) were drawn with circlize v0.4.18 and show the 20 interactions most enriched in each condition, ranked by |rank-enrichment score|.

#### Statistics and software

All analyses used biological replicates (n = 2 per condition) as the unit of replication. Multiple testing was controlled by the Benjamini–Hochberg procedure. Analyses were performed in Python 3.11.15 (Scanpy 1.11.5, anndata 0.12.19, harmonypy 2.0.0, PyDESeq2 0.5.4, gseapy 1.3.1, LIANA 1.8.1, singler 0.5.0, celldex 0.3.0, numpy 2.4.6, pandas 2.3.3, scipy 1.17.1, matplotlib 3.11.0.) and R 4.5.0 (Seurat 5.5.1, SeuratObject 5.4.0, harmony 2.0.5, circlize 0.4.18, ggplot2 4.0.3, patchwork 1.3.2, dplyr 1.2.1), with R called from Python through rpy2 3.6.8.

## Data Analysis

Statistical analyses and data visualization were performed using GraphPad Prism 11 (GraphPad Software). Normality was assessed before selecting statistical tests. For comparisons between two groups, the Mann-Whitney test was used for non-normally distributed data, and the paired t-test was used to assess repeated measurements from the same animal. For comparisons among more than two groups, one-way ANOVA with Tukey’s or Dunnett’s post hoc test was used for normally distributed data, while the Kruskal-Wallis test with Dunn’s post hoc test was used for non-normally distributed data. Data are presented as mean±SEM unless otherwise specified, and sample sizes are indicated in the corresponding figure legends. A p-value < 0.05 was considered statistically significant.

## Author contribution statement

Y.G. and S.C. wrote the manuscript. Y.G., H.W., B.C., S.C., G.S., R.J., and L.W. performed the animal experiments, Y.G., H.Y.L., J.E.M. and S.C. contributed to data analysis and figure preparation. H.W. synthesized all materials used in this study and assisted with flow staining across all experiments. B.C. collected data for a CAIA study and performed TEM. H.Y.L. processed and analyzed all scRNA-seq data.

D.C.N. performed initial synthesis and characterization of pIPA. J.T.S. performed the cell-mediated release study and optimized the protocol. X. W. isolated human PBMCs. S.C. conceptualized the manuscript and supervised the project. P.Q. supervised the single-cell RNA-seq data analysis. T.M. provided scientific guidance and supplied human PBMCs. All authors participated in critical revision and approved the final version.

## Use of AI statement

ChatGPT and Claude were used to assist with grammar and language editing. The authors take full responsibility for the accuracy and integrity of the manuscript.

## Acknowledgements

This work was supported by start-up fund from the University of Washington.

## References

1. L. Fraenkel, J. M. Bathon, B. R. England, E. W. St. Clair, T. Arayssi, K. Carandang, K. D. Deane, M. Genovese, K. K. Huston, G. Kerr, J. Kremer, M. C. Nakamura, L. A. Russell, J. A. Singh, B. J. Smith, J. A. Sparks, S. Venkatachalam, M. E. Weinblatt, M. Al-Gibbawi, J. F. Baker, K. E. Barbour, J. L. Barton, L. Cappelli, F. Chamseddine, M. George, S. R. Johnson, L. Kahale, B. S. Karam, A. M. Khamis, I. Navarro-Millán, R. Mirza, P. Schwab, N. Singh, M. Turgunbaev, A. S. Turner, S. Yaacoub, E. A. Akl, 2021 American College of Rheumatology Guideline for the Treatment of Rheumatoid Arthritis. Arthritis Care Res. 73, 924–939 (2021).

2. R. S. Hardy, K. Raza, M. S. Cooper, Therapeutic glucocorticoids: mechanisms of actions in rheumatic diseases. Nat. Rev. Rheumatol. 16, 133–144 (2020).

3. C. O. Bingham III, R. J. Looney, A. Deodhar, N. Halsey, M. Greenwald, C. Codding, B. Trzaskoma, F. Martin, S. Agarwal, A. Kelman, Immunization responses in rheumatoid arthritis patients treated with rituximab: Results from a controlled clinical trial. Arthritis Rheum. 62, 64–74 (2010).

4. Y. Guo, R. Johnson, S. Cao, Programmable long-acting injectable nanomedicines for chronic inflammation. Nano Res., doi: 10.26599/NR.2026.94909206 (2026).

5. M. E. Husni, A. Deodhar, S. Schwartzman, S. D. Chakravarty, E. C. Hsia, J. H. Leu, Y. Zhou, K. H. Lo, A. Kavanaugh, Pooled safety results across phase 3 randomized trials of intravenous golimumab in rheumatoid arthritis, psoriatic arthritis, and ankylosing spondylitis. Arthritis Res. Ther. 24, 73 (2022).

6. S. van Assen, A. Holvast, C. A. Benne, M. D. Posthumus, M. A. van Leeuwen, A. E. Voskuyl, M. Blom, A. P. Risselada, A. de Haan, J. Westra, C. G. M. Kallenberg, M. Bijl, Humoral responses after influenza vaccination are severely reduced in patients with rheumatoid arthritis treated with rituximab. Arthritis Rheum. 62, 75–81 (2010).

7. C. O. Bingham, R. J. Looney, A. Deodhar, N. Halsey, M. Greenwald, C. Codding, B. Trzaskoma, F. Martin, S. Agarwal, A. Kelman, Immunization responses in rheumatoid arthritis patients treated with rituximab: results from a controlled clinical trial. Arthritis Rheum. 62, 64–74 (2010).

8. Y. Furusawa, Y. Obata, S. Fukuda, T. A. Endo, G. Nakato, D. Takahashi, Y. Nakanishi, C. Uetake, K. Kato, T. Kato, M. Takahashi, N. N. Fukuda, S. Murakami, E. Miyauchi, S. Hino, K. Atarashi, S. Onawa, Y. Fujimura, T. Lockett, J. M. Clarke, D. L. Topping, M. Tomita, S. Hori, O. Ohara, T. Morita, H. Koseki, J. Kikuchi, K. Honda, K. Hase, H. Ohno, Commensal microbe-derived butyrate induces the differentiation of colonic regulatory T cells. Nature 504, 446–450 (2013).

9. P. V. Chang, L. Hao, S. Offermanns, R. Medzhitov, The microbial metabolite butyrate regulates intestinal macrophage function via histone deacetylase inhibition. Proc. Natl. Acad. Sci. 111, 2247– 2252 (2014).

10. R. Wang, S. Cao, M. E. H. Bashir, L. A. Hesser, Y. Su, S. M. C. Hong, A. Thompson, E. Culleen, M. Sabados, N. P. Dylla, E. Campbell, R. Bao, E. B. Nonnecke, C. L. Bevins, D. S. Wilson, J. A. Hubbell, C. R. Nagler, Treatment of peanut allergy and colitis in mice via the intestinal release of butyrate from polymeric micelles. Nat. Biomed. Eng., 1–18 (2022).

11. S. Cao, E. Budina, M. M. Raczy, A. Solanki, M. Nguyen, T. N. Beckman, J. W. Reda, K. Hultgren, P. S. Ang, A. J. Slezak, L. A. Hesser, A. T. Alpar, K. C. Refvik, L. S. Shores, I. Pillai, R. P. Wallace, A. Dhar, E. A. Watkins, J. A. Hubbell, A serine-conjugated butyrate prodrug with high oral bioavailability suppresses autoimmune arthritis and neuroinflammation in mice. Nat. Biomed. Eng., 1–17 (2024).

12. D. Dodd, M. H. Spitzer, W. Van Treuren, B. D. Merrill, A. J. Hryckowian, S. K. Higginbottom, A. Le, T. M. Cowan, G. P. Nolan, M. A. Fischbach, J. L. Sonnenburg, A gut bacterial pathway metabolizes aromatic amino acids into nine circulating metabolites. Nature 551, 648–652 (2017).

13. D. Dodd, M. H. Spitzer, W. Van Treuren, B. D. Merrill, A. J. Hryckowian, S. K. Higginbottom, A. Le, T. M. Cowan, G. P. Nolan, M. A. Fischbach, J. L. Sonnenburg, A gut bacterial pathway metabolizes aromatic amino acids into nine circulating metabolites. Nature 551, 648–652 (2017).

14. M. Venkatesh, S. Mukherjee, H. Wang, H. Li, K. Sun, A. P. Benechet, Z. Qiu, L. Maher, M. R. Redinbo, R. S. Phillips, J. C. Fleet, S. Kortagere, P. Mukherjee, A. Fasano, J. Le Ven, J. K. Nicholson, M. E. Dumas, K. M. Khanna, S. Mani, Symbiotic Bacterial Metabolites Regulate Gastrointestinal Barrier Function via the Xenobiotic Sensor PXR and Toll-like Receptor 4. Immunity 41, 296–310 (2014).

15. Q. Li, R. de Oliveira Formiga, V. Puchois, L. Creusot, A. H. Ahmad, S. Amouyal, M. A. Campos-Ribeiro, Y. Zhao, D. M. M. Harris, F. Lasserre, S. Ellero-Simatos, H. Guillou, Z. Huang, L. Brot, Y. Hu, L. Chollet, C. Danne, C. Scandola, T. Ledent, G. Chevreux, R. J. Argüello, M. De Carvalho Bittencourt, J. Bettinger, M. D’Aveni-Piney, D. Moulin, S. Schreiber, K. Aden, N. Rolhion, M.-L. Michel, T. Wai, H. Sokol, Microbial metabolite indole-3-propionic acid drives mitochondrial respiration in CD4+ T cells to confer protection against intestinal inflammation. Nat. Metab. 7, 2510–2530 (2025).

16. C. Yang, L. Xie, Z. Deng, H. Ai, T. Xiang, X. Yan, Z. Ling, S. Xiao, Y. Tang, G. Huang, F. Luo, Y. Chen, An orally-administered nanotherapeutics with gold nanospheres supplying for rheumatoid arthritis therapy by re-shaping gut microbial tryptophan metabolism. J. Nanobiotechnology 23, 376 (2025).

17. S. T. Reddy, A. J. van der Vlies, E. Simeoni, V. Angeli, G. J. Randolph, C. P. O’Neil, L. K. Lee, M. A. Swartz, J. A. Hubbell, Exploiting lymphatic transport and complement activation in nanoparticle vaccines. Nat. Biotechnol. 25, 1159–1164 (2007).

18. G. P. Howard, G. Verma, X. Ke, W. M. Thayer, T. Hamerly, V. K. Baxter, J. E. Lee, R. R. Dinglasan, H.-Q. Mao, Critical size limit of biodegradable nanoparticles for enhanced lymph node trafficking and paracortex penetration. Nano Res. 12, 837–844 (2019).

19. S. Cao, E. Budina, R. Wang, M. Sabados, A. Mukherjee, A. Solanki, M. Nguyen, K. Hultgren, A. Dhar, J. A. Hubbell, Injectable butyrate-prodrug micelles induce long-acting immune modulation and prevent autoimmune arthritis in mice. J. Controlled Release 372, 281–294 (2024).

20. S. T. Reddy, A. J. van der Vlies, E. Simeoni, V. Angeli, G. J. Randolph, C. P. O’Neil, L. K. Lee, M. A. Swartz, J. A. Hubbell, Exploiting lymphatic transport and complement activation in nanoparticle vaccines. Nat. Biotechnol. 25, 1159–1164 (2007).

21. E. Blanco, H. Shen, M. Ferrari, Principles of nanoparticle design for overcoming biological barriers to drug delivery. Nat. Biotechnol. 33, 941–951 (2015).

22. K. S. Nandakumar, J. Bäcklund, M. Vestberg, R. Holmdahl, Collagen type II (CII)-specific antibodies induce arthritis in the absence of T or B cells but the arthritis progression is enhanced by CII-reactive T cells. Arthritis Res Ther 6, R544 (2004).

23. A. Petrelli, G. Mijnheer, D. P. H. van Konijnenburg, M. M. van der Wal, B. Giovannone, E. Mocholi, N. Vazirpanah, J. C. Broen, D. Hijnen, B. Oldenburg, P. J. Coffer, S. J. Vastert, B. J. Prakken, E. Spierings, A. Pandit, M. Mokry, F. van Wijk, PD-1+CD8+ T cells are clonally expanding effectors in human chronic inflammation. J. Clin. Invest. 128, 4669–4681.

24. C.-Q. Chu, D. Swart, D. Alcorn, J. Tocker, K. B. Elkon, Interferon-gamma regulates susceptibility to collagen-induced arthritis through suppression of interleukin-17. Arthritis Rheum. 56, 1145–1151 (2007).

25. M. Guilliams, C.-A. Dutertre, C. L. Scott, N. McGovern, D. Sichien, S. Chakarov, S. Van Gassen, J. Chen, M. Poidinger, S. De Prijck, S. J. Tavernier, I. Low, S. E. Irac, C. N. Mattar, H. R. Sumatoh, G. H. L. Low, T. J. K. Chung, D. K. H. Chan, K. K. Tan, T. L. K. Hon, E. Fossum, B. Bogen, M. Choolani, J. K. Y. Chan, A. Larbi, H. Luche, S. Henri, Y. Saeys, E. W. Newell, B. N. Lambrecht, B. Malissen, F. Ginhoux, Unsupervised High-Dimensional Analysis Aligns Dendritic Cells across Tissues and Species. Immunity 45, 669–684 (2016).

26. E. K. Persson, H. Uronen-Hansson, M. Semmrich, A. Rivollier, K. Hägerbrand, J. Marsal, S. Gudjonsson, U. Håkansson, B. Reizis, K. Kotarsky, W. W. Agace, IRF4 Transcription-Factor-Dependent CD103+CD11b+ Dendritic Cells Drive Mucosal T Helper 17 Cell Differentiation. Immunity 38, 958–969 (2013).

27. X. Zhang, C. S. McGinnis, G. Yu, S. Chen, P. Zheng, C. M. Schürch, K. J. Hiam-Galvez, N. E. Reticker-Flynn, W. Guo, W. Yao, J. Qiu, A. Muselman, I. L. Linde, J. W. Hickey, H. Yan, V. M. Tran, W. Qiu, D. Brichart-Vernos, T. Hirai, B. Yu, X. An, Y. Xiao, H. Paidassi, T. C. Scharschmidt, M. Angelo, D. Sheppard, H. Chi, A. T. Satpathy, S. S. Way, B. Malissen, S. Strober, E. G. Engleman, Erythropoietin receptor on cDC1s dictates immune tolerance. Nature 650, 470–480 (2026).

28. M. I. Ramos, S. Garcia, B. Helder, S. Aarrass, Kris, A. Reedquist, S. E. Jacobsen, P. P. Tak, M. C. Lebre, cDC1 are required for the initiation of collagen-induced arthritis. J. Transl. Autoimmun. 3, 100066 (2020).

29. Z. Jiao, S. Bedoui, J. L. Brady, A. Walter, M. Chopin, E. M. Carrington, R. M. Sutherland, S. L. Nutt, Y. Zhang, H.-J. Ko, L. Wu, A. M. Lew, Y. Zhan, The Closely Related CD103+ Dendritic Cells (DCs) and Lymphoid-Resident CD8+ DCs Differ in Their Inflammatory Functions. PLOS ONE 9, e91126 (2014).

30. B. T. Edelson, W. KC, R. Juang, M. Kohyama, L. A. Benoit, P. A. Klekotka, C. Moon, J. C. Albring, W. Ise, D. G. Michael, D. Bhattacharya, T. S. Stappenbeck, M. J. Holtzman, S.-S. J. Sung, T. L. Murphy, K. Hildner, K. M. Murphy, Peripheral CD103+ dendritic cells form a unified subset developmentally related to CD8α+ conventional dendritic cells. J. Exp. Med. 207, 823–836 (2010).

31. J. Tuttle, E. Drescher, J. A. Simón-Campos, P. Emery, M. Greenwald, A. Kivitz, H. Rha, P. Yachi, C. Kiley, A. Nirula, A Phase 2 Trial of Peresolimab for Adults with Rheumatoid Arthritis. N. Engl. J. Med. 388, 1853–1862 (2023).

32. K. A. Fernandes, F. Sanders, T. R. Cafiero, C. Huang, J. Reyes, A. Biswas, A. I. Lim, Maternal helminths rewire the microbiota to promote offspring antiviral immunity. Cell Host Microbe 34, 1050–1066.e9 (2026).

