## Supplementary Figures and Tables for "Indole-3-propionic acid nano-prodrugs sustain increased lymphoid exposure and alleviate experimental autoimmune arthritis"

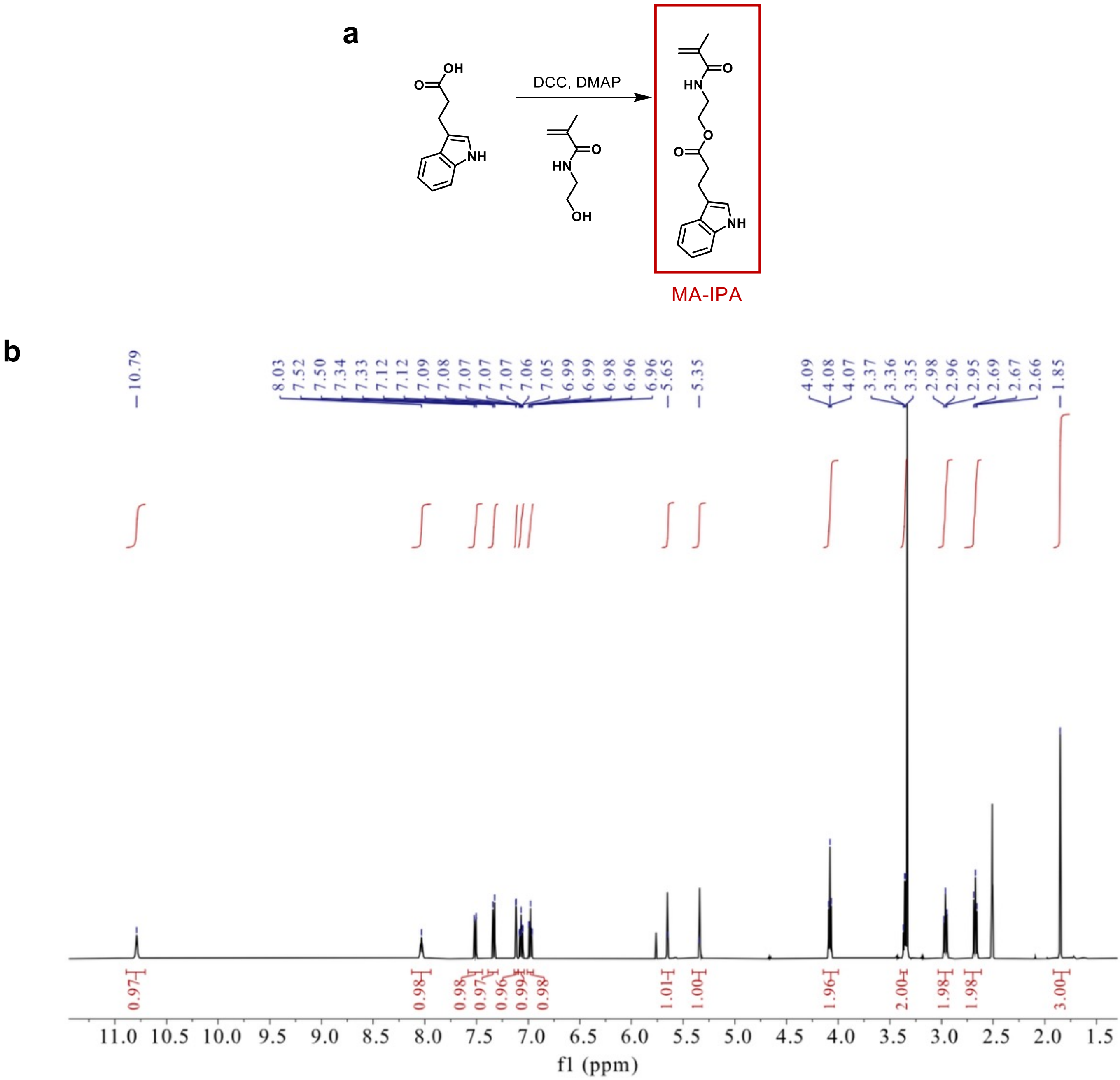


**Supplementary Figure S1. Synthesis and characterization of MA-IPA. a.** reaction scheme for the synthesis of [2-(methacryloylamino)ethyl]indole-3-propionate monomer (MA-IPA) from IPA. **b.** $^{1}$H NMR Spectrum for IPA monomer. 1 H NMR (500 MHz, DMSO-d 6 , δ, ppm): 1.85 (s, 3H, −CCH 3 ), 2.67 (t, 2H, −CH 2 C=O), 2.96 (t, 2H, −CH 2 NH), 3.36 (t, 2H, C 8 H 5 NH-CH 2 ), 4.08(t, 2H, −OCH 2 ), 5.35 and 5.65 (s, 2H, −C=CH 2 ), 6.96–7.52 (m, 5H, indole ring), 8.03 (s, 1H, −NHC=O), 10.79 (s, 1H, −NHCH).

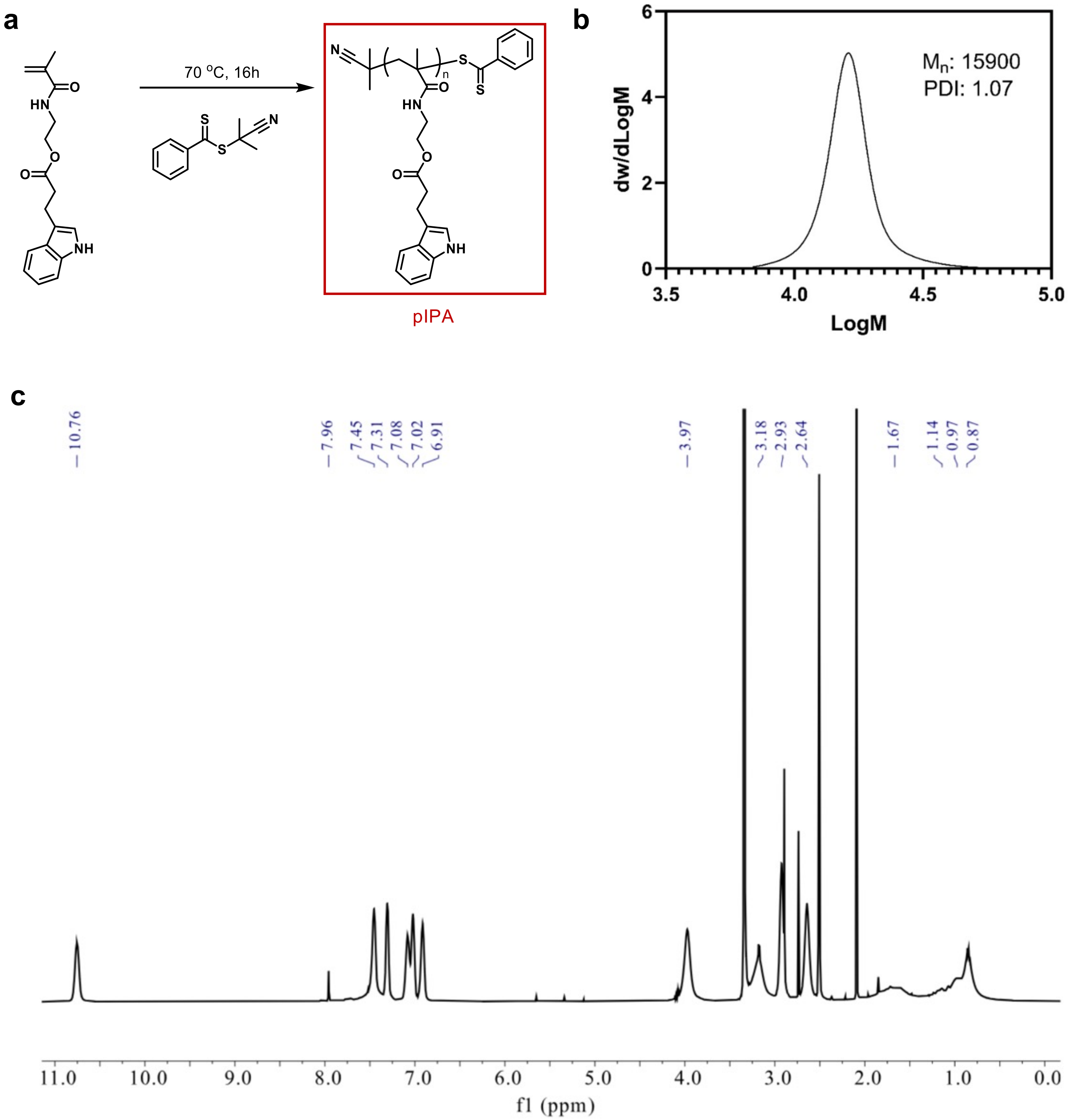


**Supplementary Figure S2. Synthesis and characterization of pIPA. a.** Reaction scheme of pIPA from MA-IPA monomers. **b.** GPC Spectrum for pIPA. **c.** $^{1}$H NMR Spectrum for IPA polymer. 1 H NMR (500 MHz, DMSO-d6 , δ, ppm): 0.87-1.14 (m, 3H,backbone CH 3 ), 1.67 (m, 2H, backbone CH 2 ), 2.64 (m, 2H, CH 2 C=O), 2.93(m, 2H, CH 2 NH), 3.18(m, 2H, C 8 H 5 NH-CH 2 ), 3.97(m, 2H, OCH 2 ), 6.91-7.45(m, 5H, indole ring), 7.96(m, 1H, NHC=O), 10.76(m, 1H, NHCH) (Appendix Figure 2).

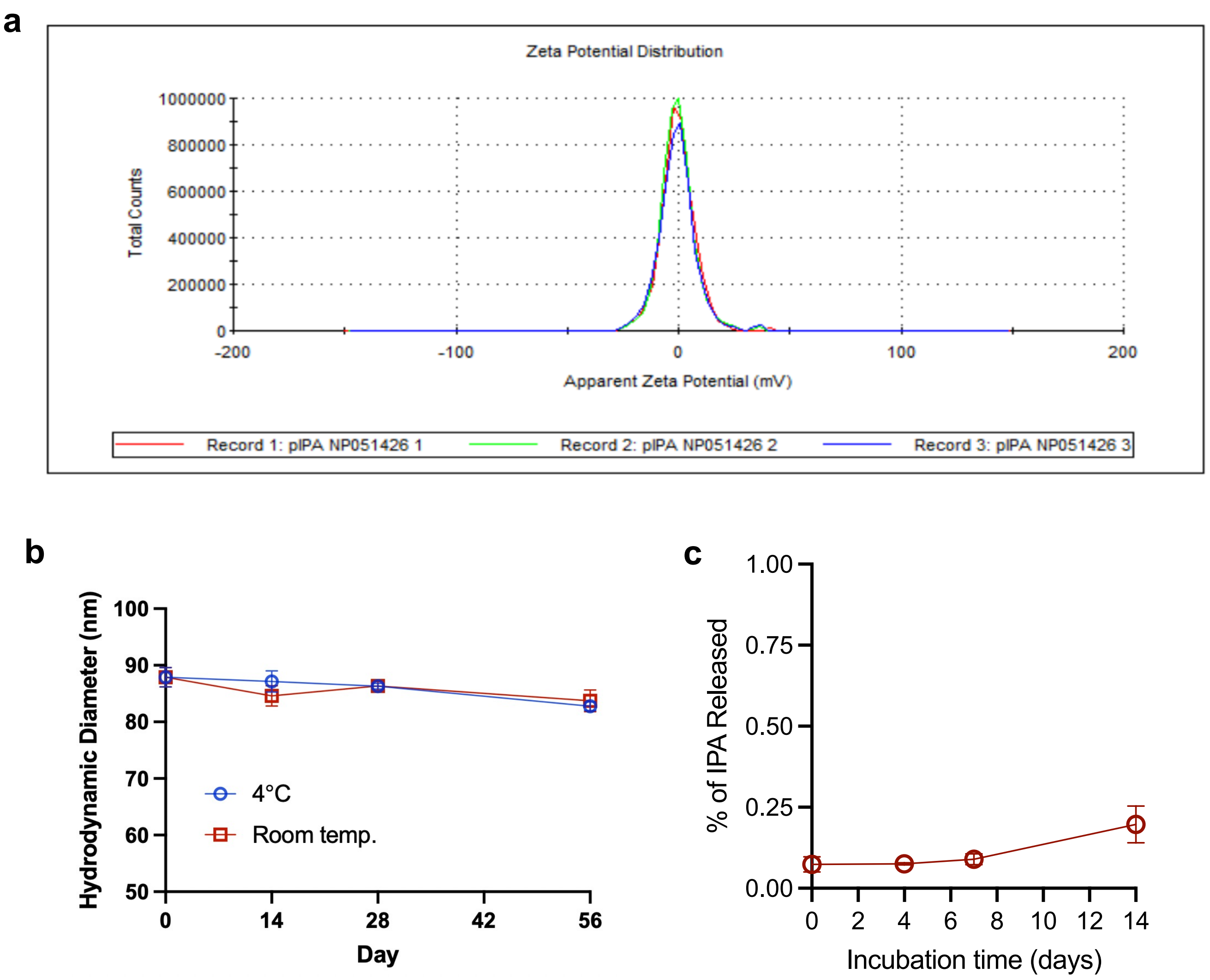


**Supplementary Figure S3. Additional characterization of pIPA$_{NP}$. a.** Zeta potential distribution of pIPA$_{NP}$ measured by DLS. **b.** Long-term size stability of pIPA$_{NP}$ at 4 ˚C and at room temperature in water, n=3 for each time point. c. Cumulative IPA release from pIPA$_{NP}$ over 14 days at 37 ˚C in DMEM media containing 10% non-heat-inactivated fetal bovine serum, quantified by LC-MS/MS.

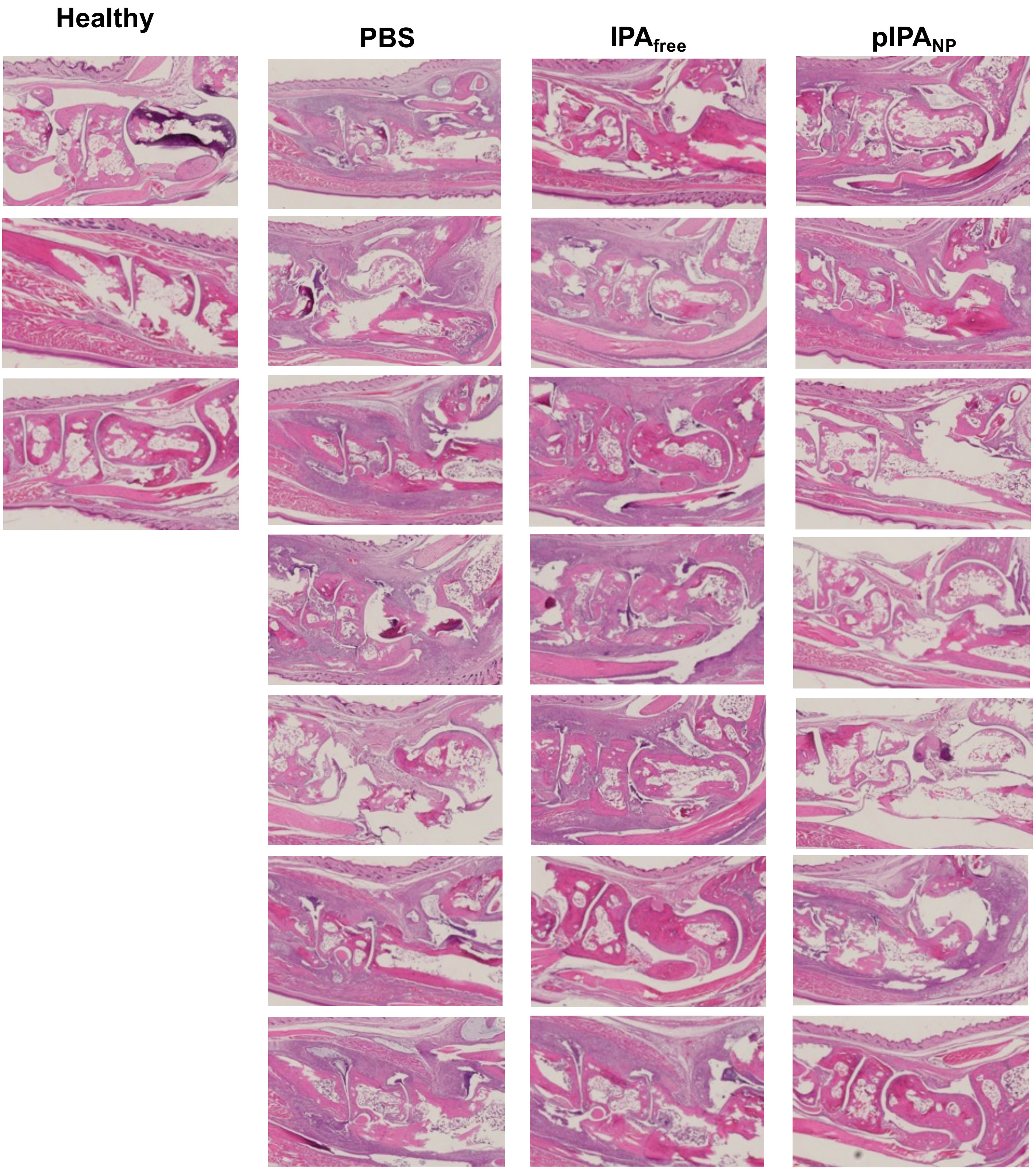


**Supplementary Figure S4. Histology of paw sections from the prophylactic CAIA study.** Hematoxylin and eosin (H&E)-stained joint sections from all mice in the prophylactic study (Figure 2), harvested on Day 12.

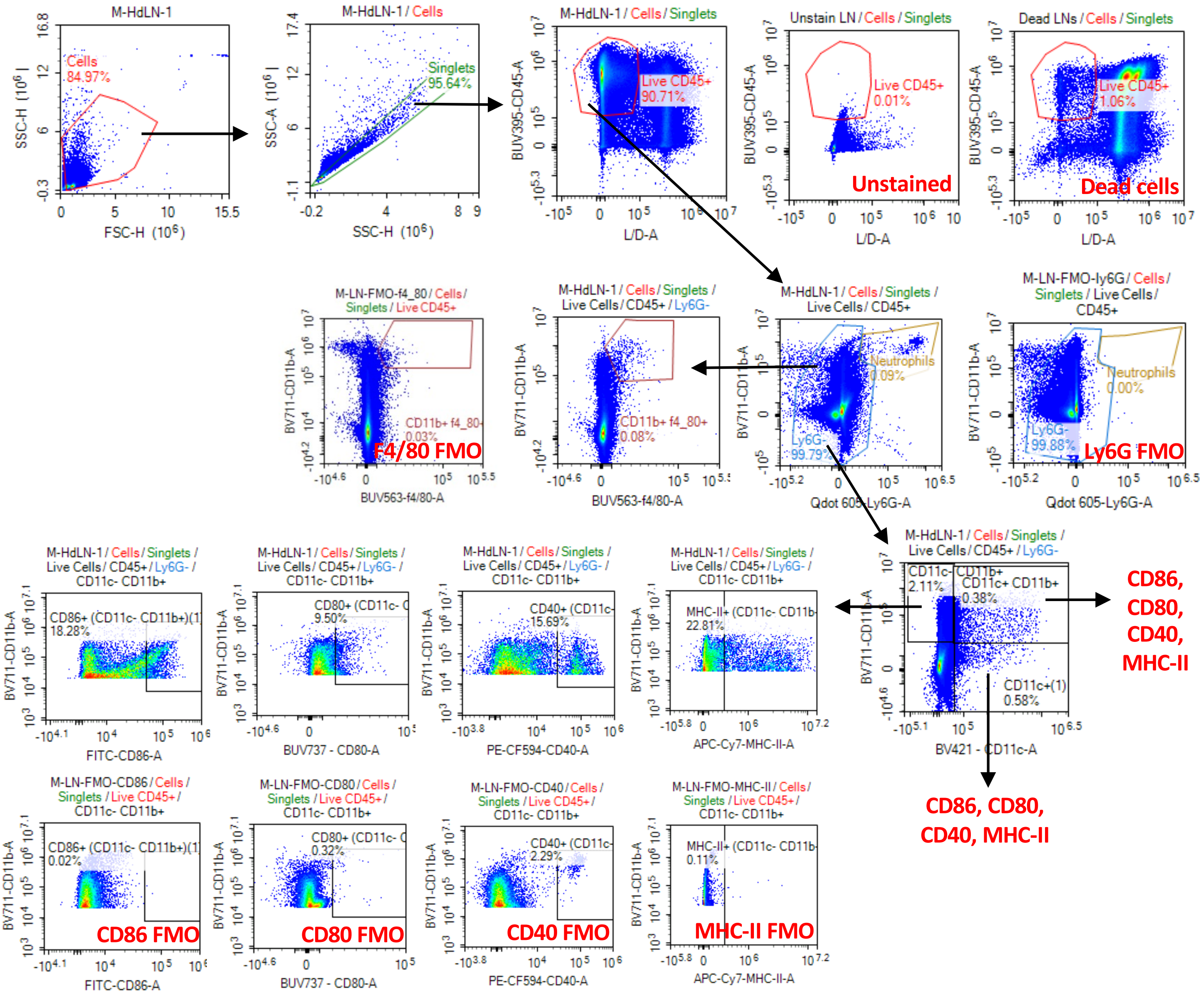


**Supplementary Figure S5. Flow cytometry gating strategy for myeloid subsets and costimulatory markers.** Myeloid subsets were defined as CD11b$^+$F4/80$^+$, CD11b$^+$CD11c$^-$, CD11b$^+$CD11c$^+$, CD11c$^+$CD11b$^-$, and CD103$^+$CD11c$^+$ cells, within which the percentages of CD40$^+$, CD80$^+$, CD86$^+$ and MHC-II$^+$ cells were determined. This gating strategy applies to the data in Figures 3a, 4h and 6e.

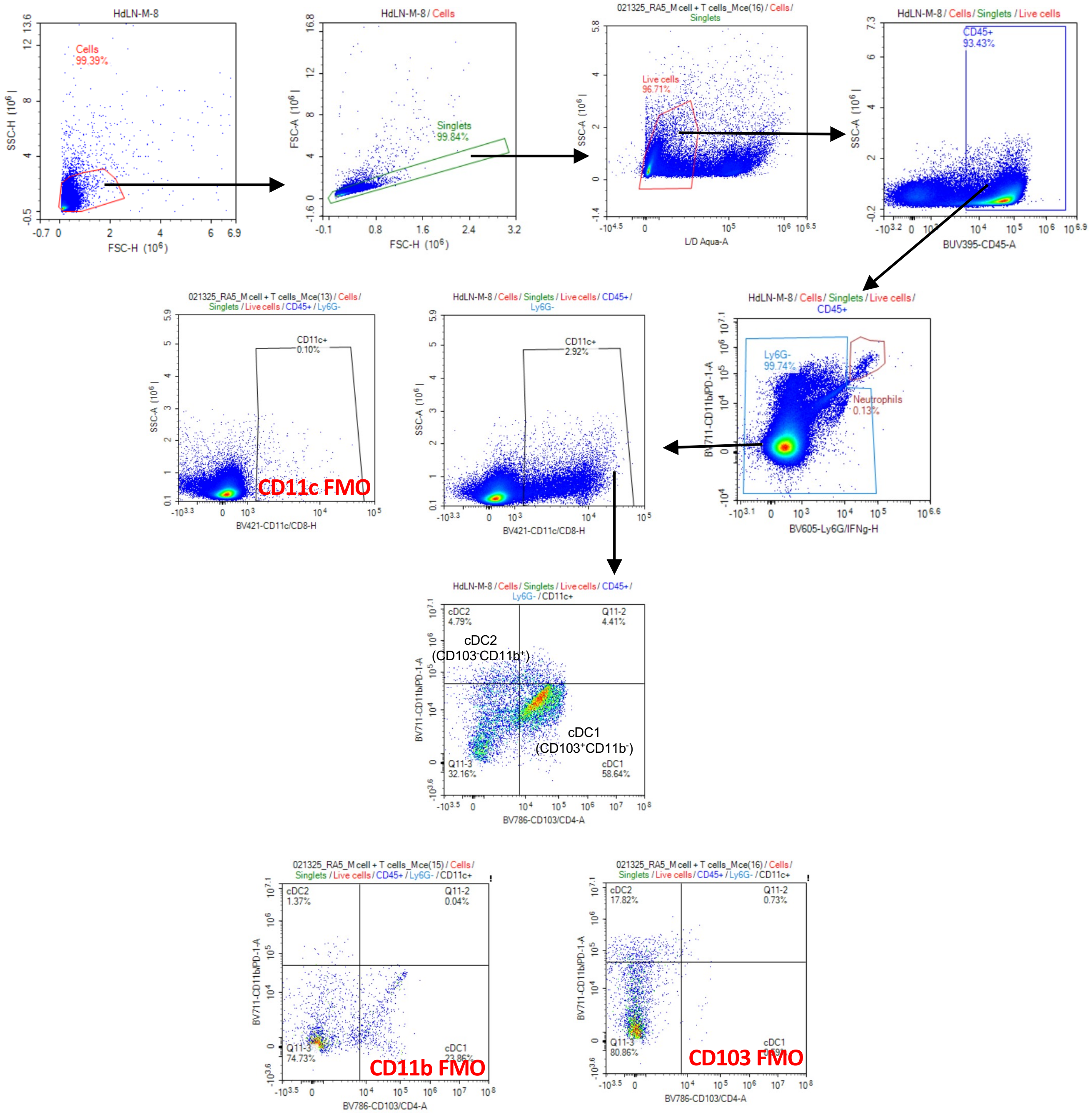


**Supplementary Figure S6. Flow cytometry gating strategy for cDC1 and cDC2.** Conventional dendritic cell 1 and 2 were defined as $CD11b^{-}CD103^{+}$ (cDC1) and $CD11b^{+}CD103^{-}$ (cDC2) cells. This gating strategy applies to the data in Figures 3b and 4g.

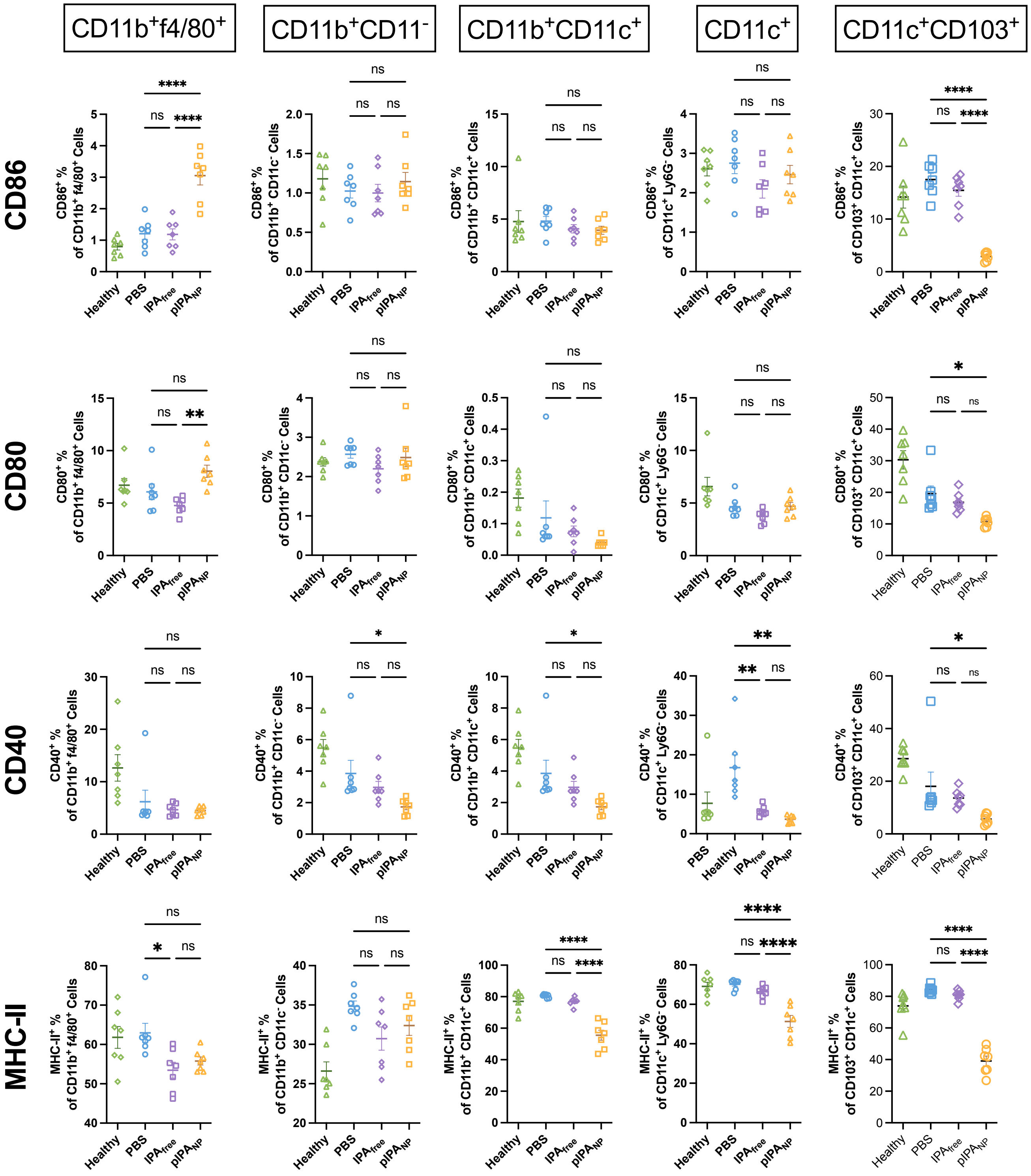


**Supplementary Figure S7. Percentages of costimulatory marker- and MHC-II-positive cells among splenic myeloid subsets in the prophylactic CAIA study.** Percentage of CD86$^{+}$, CD80$^{+}$, CD40$^{+}$ and MHC-II$^{+}$ cells within CD11b$^{+}$F4/80$^{+}$, CD11b$^{+}$CD11c$^{-}$, CD11b$^{+}$CD11c$^{+}$, CD11c$^{+}$CD11b$^{-}$ and CD103$^{+}$CD11c$^{+}$ cells (n=7). The data correspond to the heatmap in Figure 3a. Data are presented as mean ± SEM. Groups were compared by one-way ANOVA with Tukey's multiple comparisons test. *p < 0.05, **p < 0.01, ***p < 0.001, ****p < 0.0001; ns, not significant.

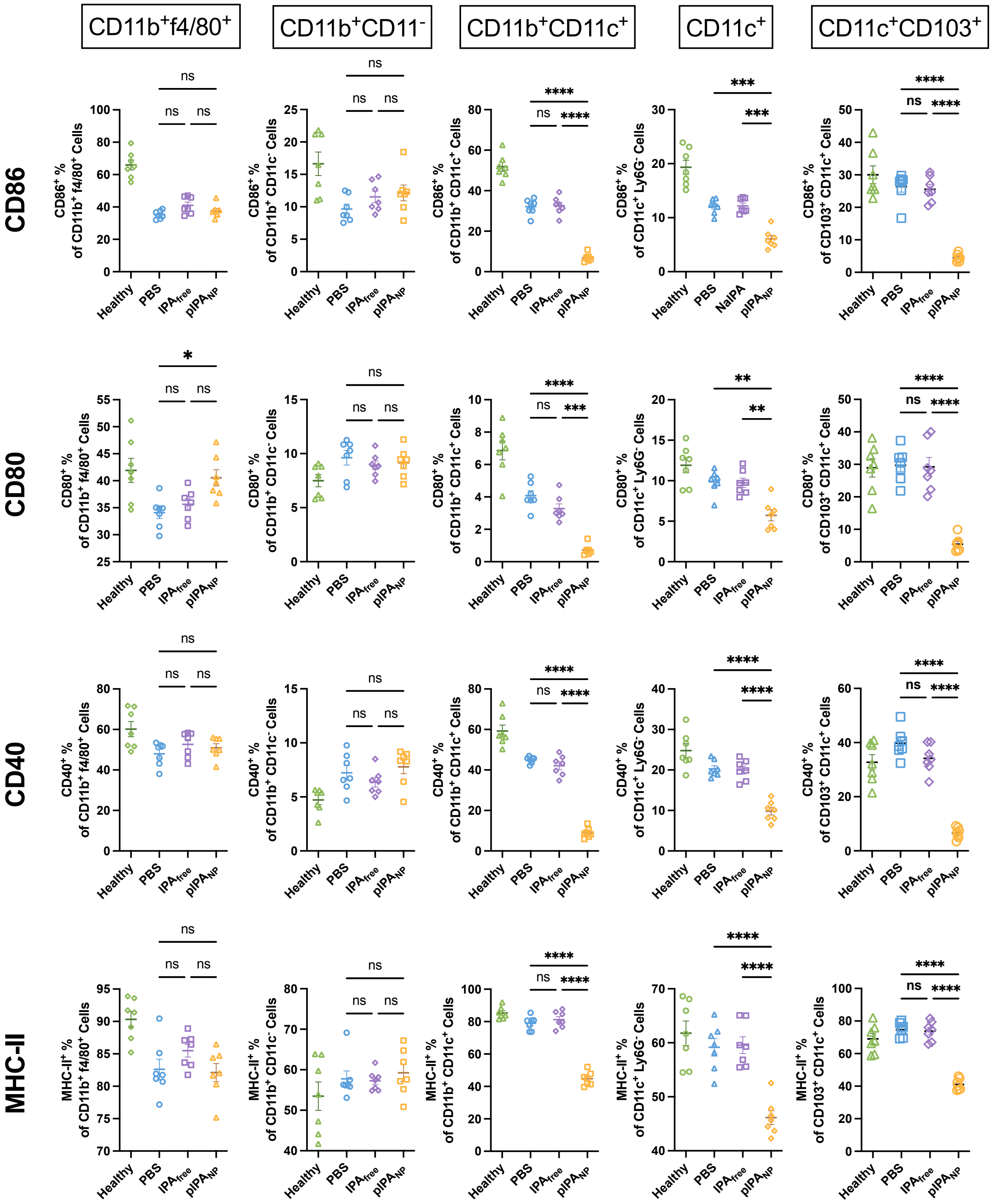


**Supplementary Figure S8. Percentages of costimulatory marker- and MHC-II-positive cells among hock-draining lymph node myeloid subsets in the prophylactic CAIA study.** Percentage of CD86$^{+}$, CD80$^{+}$, CD40$^{+}$ and MHC-II$^{+}$ cells within CD11b$^{+}$F4/80$^{+}$, CD11b$^{+}$CD11c$^{-}$, CD11b$^{+}$CD11c$^{+}$, CD11c$^{+}$CD11b$^{-}$ and CD103$^{+}$CD11c$^{+}$ cells (n=7). The data correspond to the heatmap in Figure 3a. Data are presented as mean ± SEM. Groups were compared by one-way ANOVA with Tukey's multiple comparisons test. *p < 0.05, **p < 0.01, ***p < 0.001, ****p < 0.0001; ns, not significant.

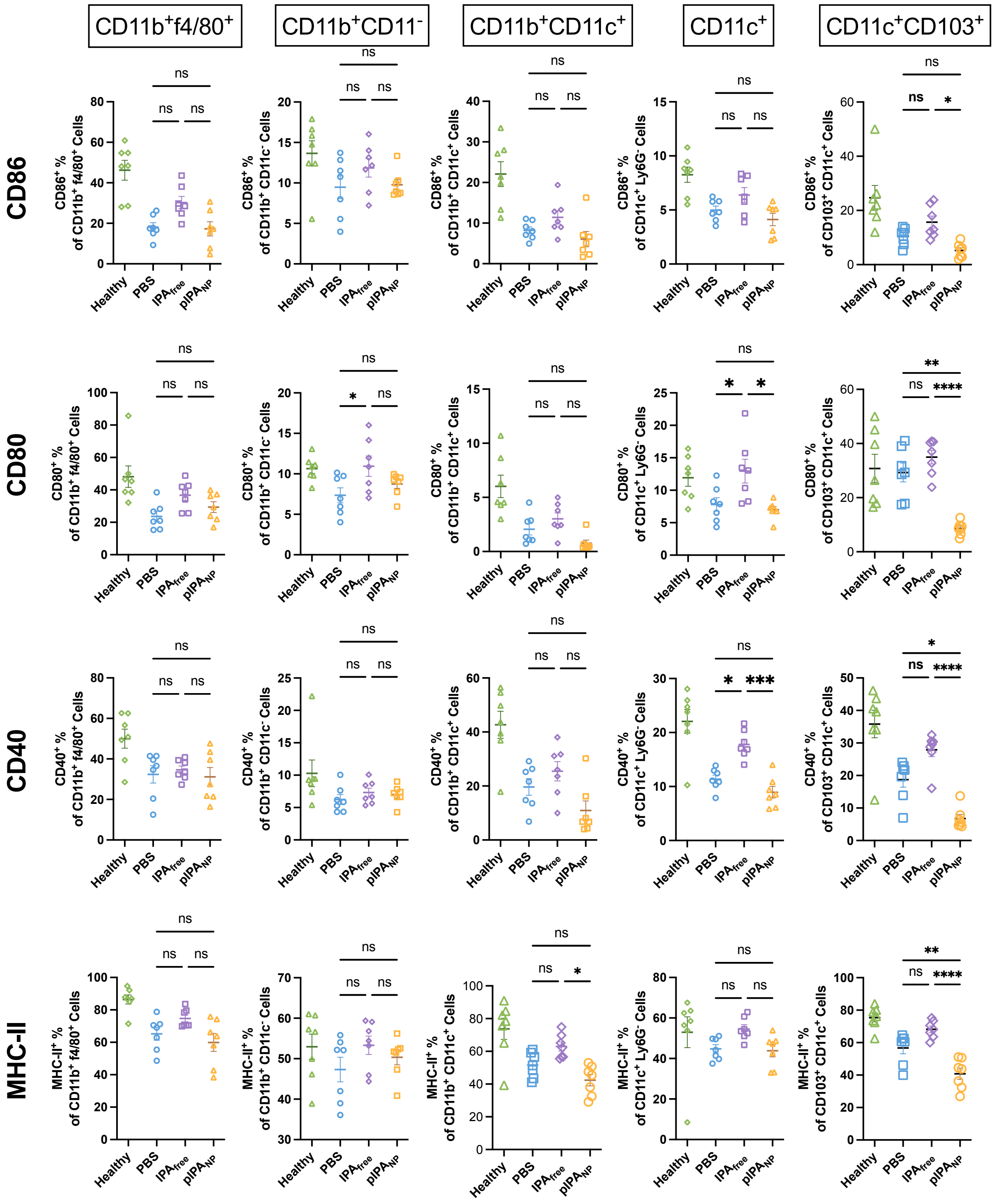


**Supplementary Figure S9. Percentages of costimulatory marker- and MHC-II-positive cells among inguinal lymph node myeloid subsets in the prophylactic CAIA study.** Percentage of $CD86^+$, $CD80^+$, $CD40^+$ and $MHC\text{-}II^+$ cells within $CD11b^+F4/80^+$, $CD11b^+CD11c^-$, $CD11b^+CD11c^+$, $CD11c^+CD11b^-$ and $CD103^+CD11c^+$ cells (n=7). The data correspond to the heatmap in Figure 3a. Data are presented as mean ± SEM. Groups were compared by one-way ANOVA with Tukey's multiple comparisons test. *$p < 0.05$, **$p < 0.01$, ***$p < 0.001$, ****$p < 0.0001$; ns, not significant.

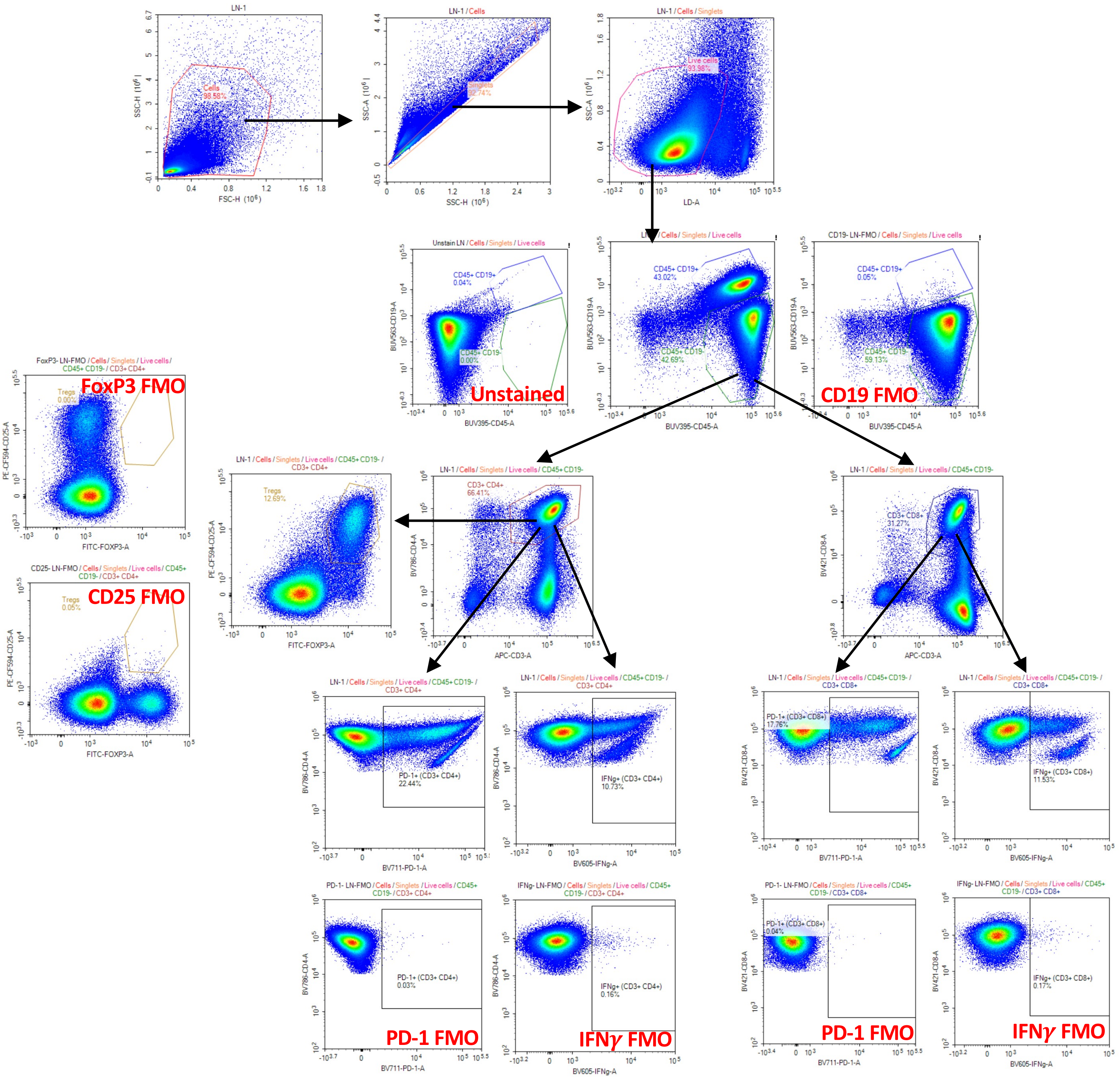


**Supplementary Figure S10. Flow cytometry gating strategy for T cell subsets.** Percentage of PD-1$^{+}$ and IFN$\gamma$+ cells among CD3$^{+}$CD4$^{+}$ and CD3$^{+}$CD8$^{+}$ subsets for Figure 3c.

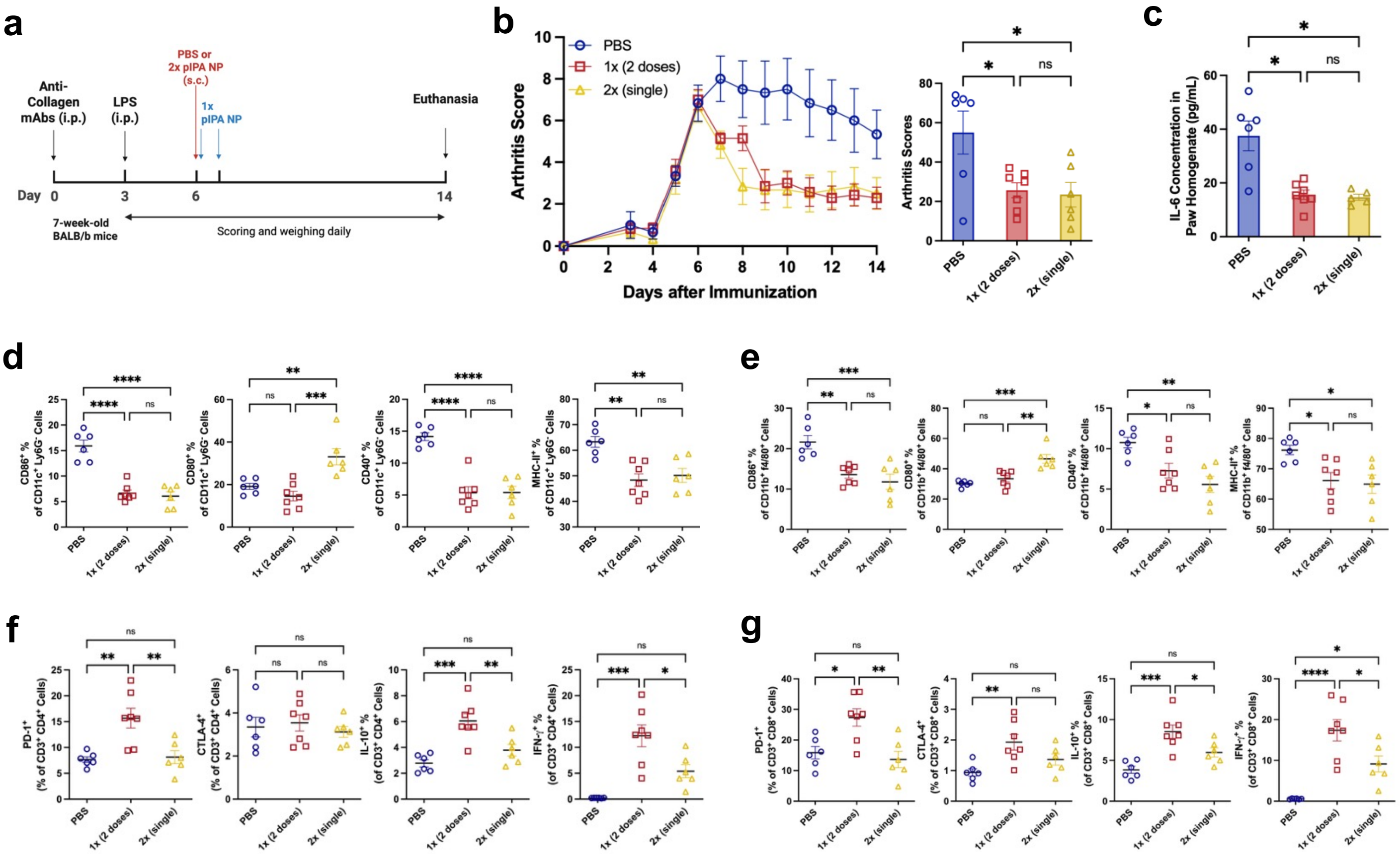


**Supplementary Figure S11. pIPA$_{NP}$ Administration in a Therapeutic CAIA Model Across Two Dosing Regimens. a.** Experimental timeline of the study. **b.** Clinical arthritis scores of mice from Day 0 to Day 14, along with the cumulative arthritis scores from Day 6 to Day 14. **c.** IL-6 concentrations in mouse paw homogenates. **d.** Percentage of CD86$^{+}$, CD80$^{+}$, CD40$^{+}$, and MHC-II$^{+}$ in CD11c$^{+}$Ly6G$^{-}$ cells and in **e.** CD11b$^{+}$f4/80$^{+}$ cells in hdLNs. **f.** Percentage of PD-1$^{+}$ and CTLA-4$^{+}$ cells among CD4$^{+}$ and **g.** CD8$^{+}$ T cells in hdLNs. Data are presented as mean±SEM (PBS: n = 6; 2x pIPA NP: n = 6; 1x pIPA NP: n = 7). Statistical comparisons were performed using ordinary one-way ANOVA with Tukey's multiple comparisons test for Fig b and c, and with the Kruskal-Wallis test with Dunn's post hoc test for d-g, depending on data distribution. *p < 0.05, **p < 0.01, ***p < 0.001, ****p < 0.0001; ns, not significant.

### PBS

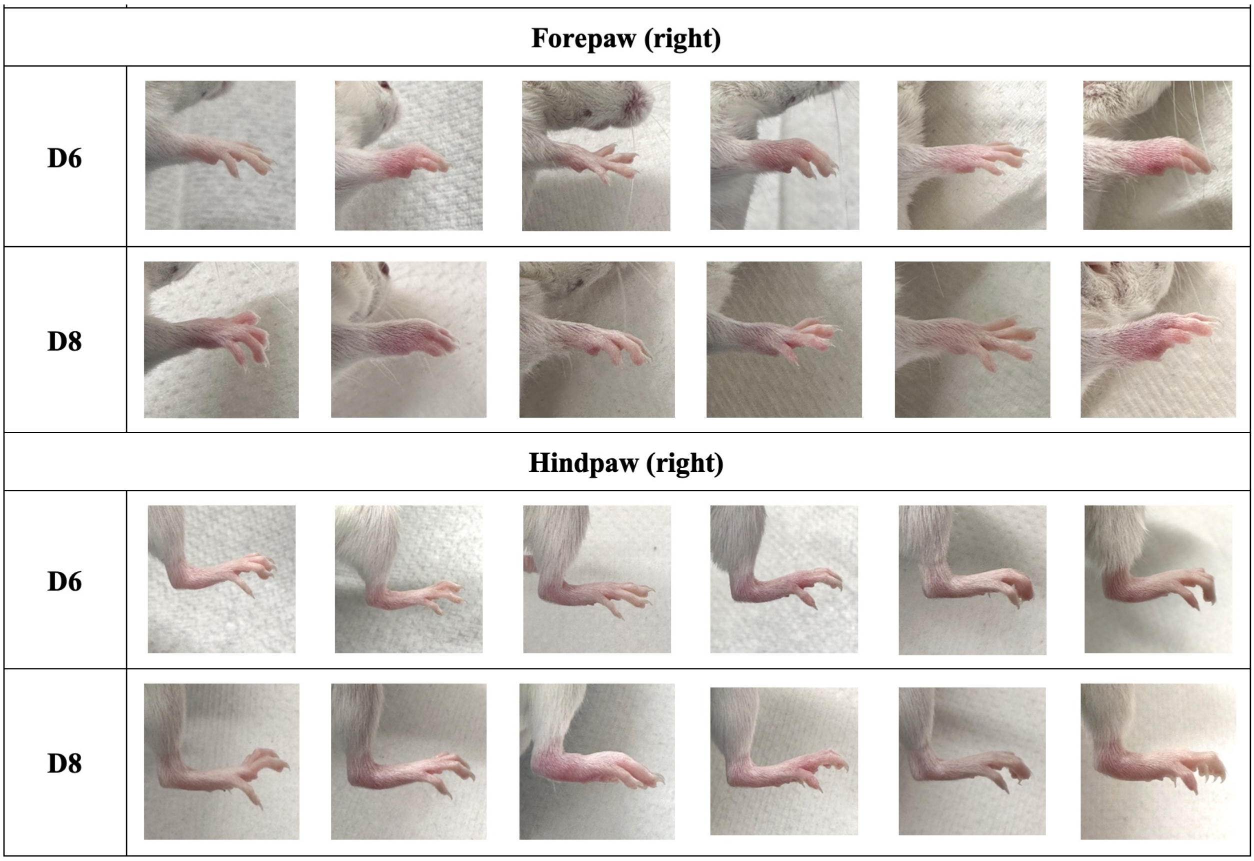


### $pIPA_{NP}$

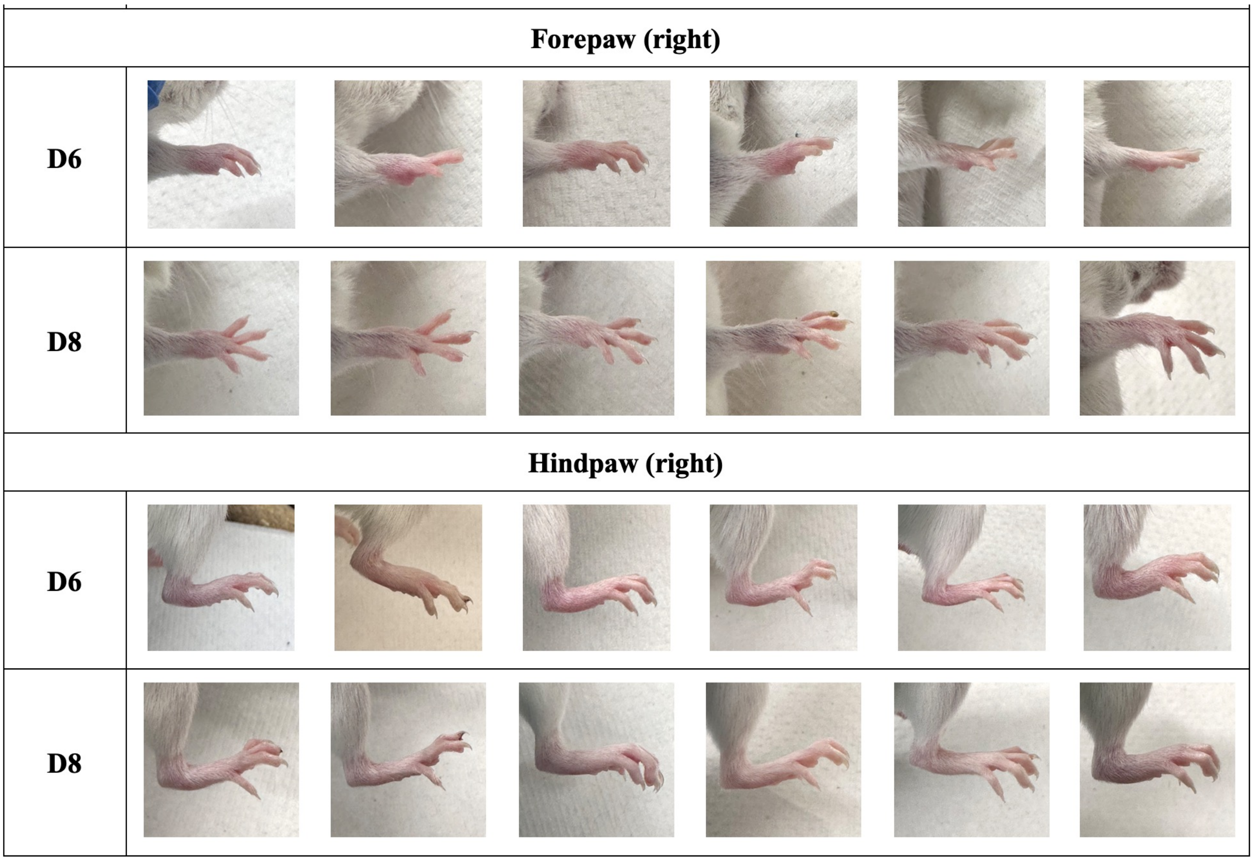


**Supplementary Figure S12.** Representative photographs of mouse paws from PBS-treated animals before (Day 6) and after (Day 8) treatment in the interventional CAIA study in Figure 4.

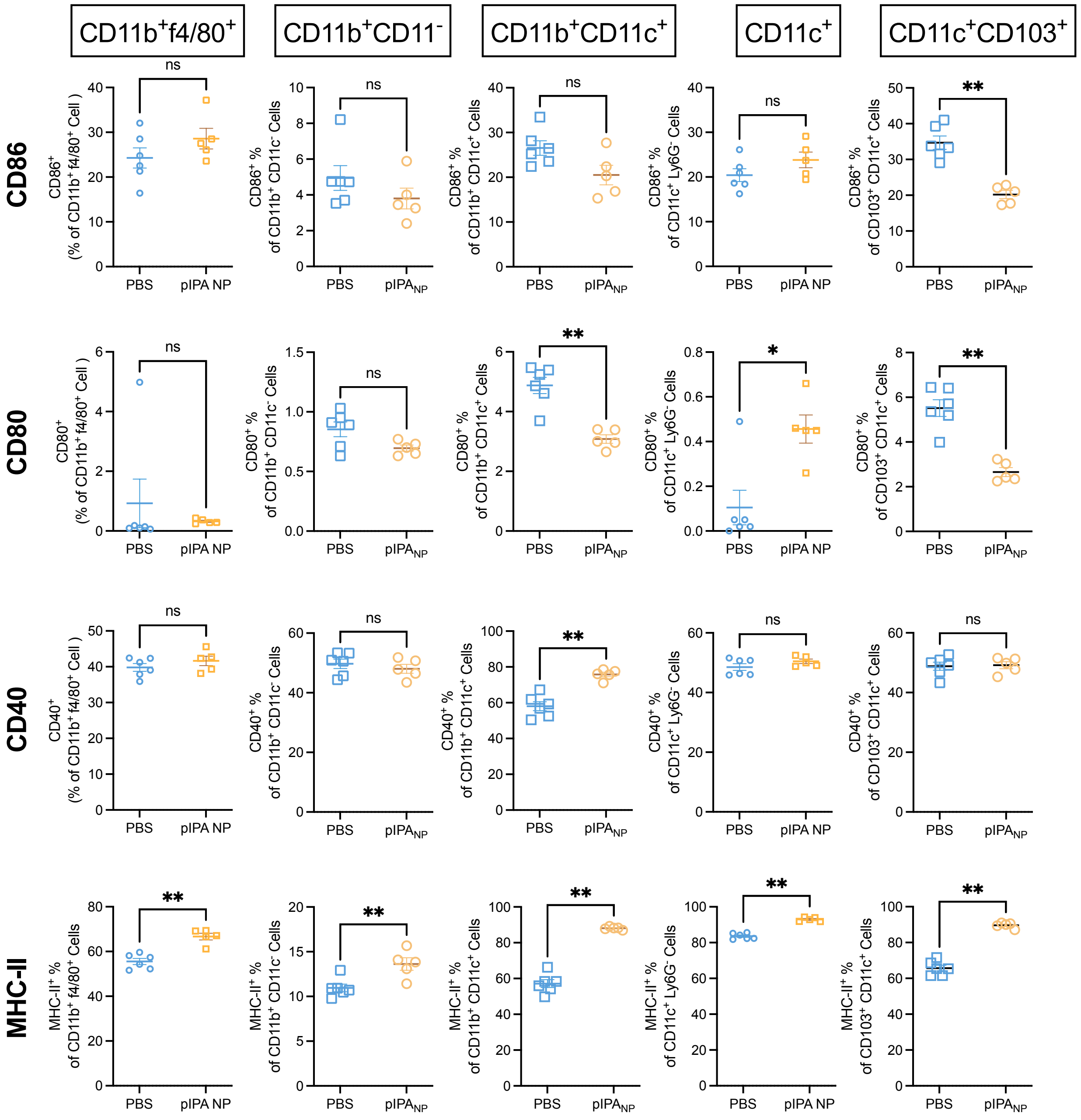


**Supplementary Figure S13. Percentages of costimulatory marker- and MHC-II-positive cells among spleen myeloid subsets in the therapeutic CAIA study.** Percentage of $CD86^+$, $CD80^+$, $CD40^+$ and MHC-$II^+$ cells within $CD11b^+F4/80^+$, $CD11b^+CD11c^−$, $CD11b^+CD11c^+$, $CD11c^+$ and $CD103^+CD11c^+$ cells in spleen (n=6). The data correspond to the heatmap in Figure 4h. Data are presented as mean ± SEM. Groups were compared by a two-tailed Mann-Whitney test. *$p < 0.05$, **$p < 0.01$, ns, not significant.

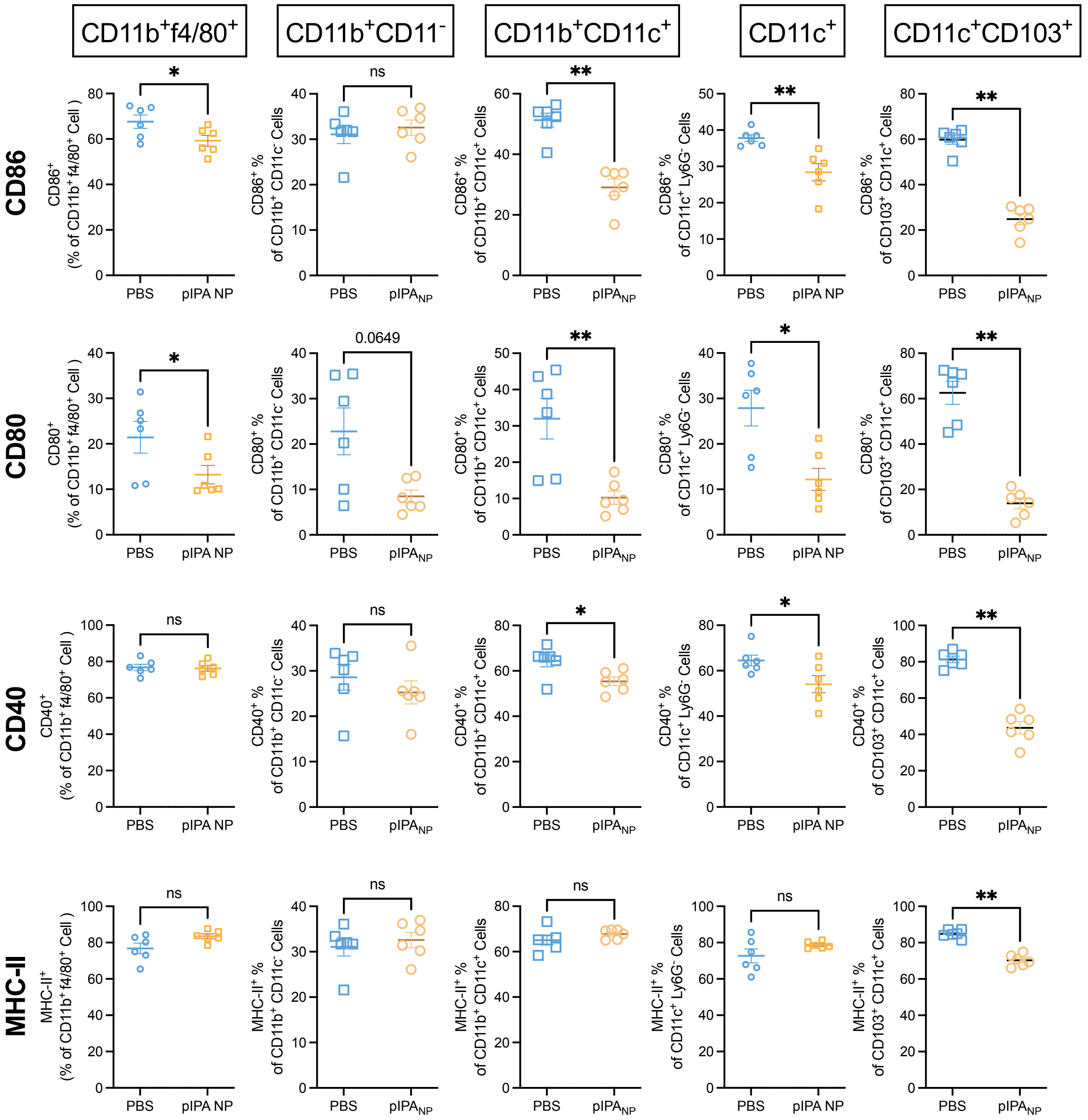


**Supplementary Figure S14. Percentages of costimulatory marker- and MHC-II-positive cells among hock-draining lymph node myeloid subsets in the therapeutic CAIA study.** Percentage of CD86$^+$, CD80$^+$, CD40$^+$ and MHC-II$^+$ cells within CD11b$^+$F4/80$^+$, CD11b$^+$CD11c$^-$, CD11b$^+$CD11c$^+$, CD11c$^+$ and CD103$^+$CD11c$^+$ cells in spleen (n=6). The data correspond to the heatmap in Figure 4h. Data are presented as mean ± SEM. Groups were compared by a two-tailed Mann-Whitney test. *p < 0.05, **p < 0.01, ns, not significant.

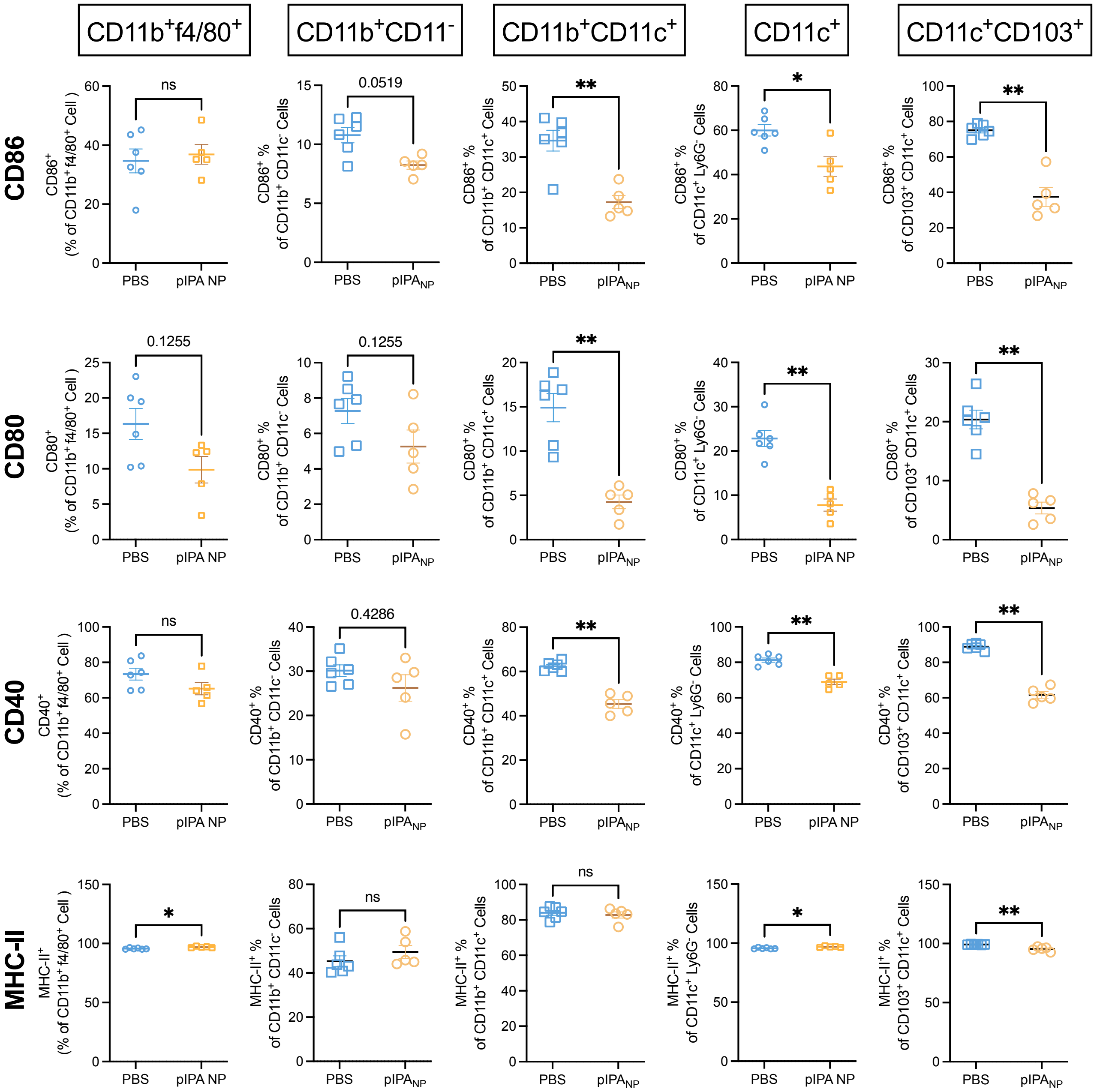


**Supplementary Figure S15. Percentages of costimulatory marker- and MHC-II-positive cells among mesenteric lymph node myeloid subsets in the therapeutic CAIA study.** Percentage of CD86⁺, CD80⁺, CD40⁺ and MHC-II⁺ cells within CD11b⁺F4/80⁺, CD11b⁺CD11c⁻, CD11b⁺CD11c⁺, CD11c⁺ and CD103⁺CD11c⁺ cells in spleen (n=6). The data correspond to the heatmap in Figure 4h. Data are presented as mean ± SEM. Groups were compared by a two-tailed Mann-Whitney test. *p < 0.05, **p < 0.01, ns, not significant.

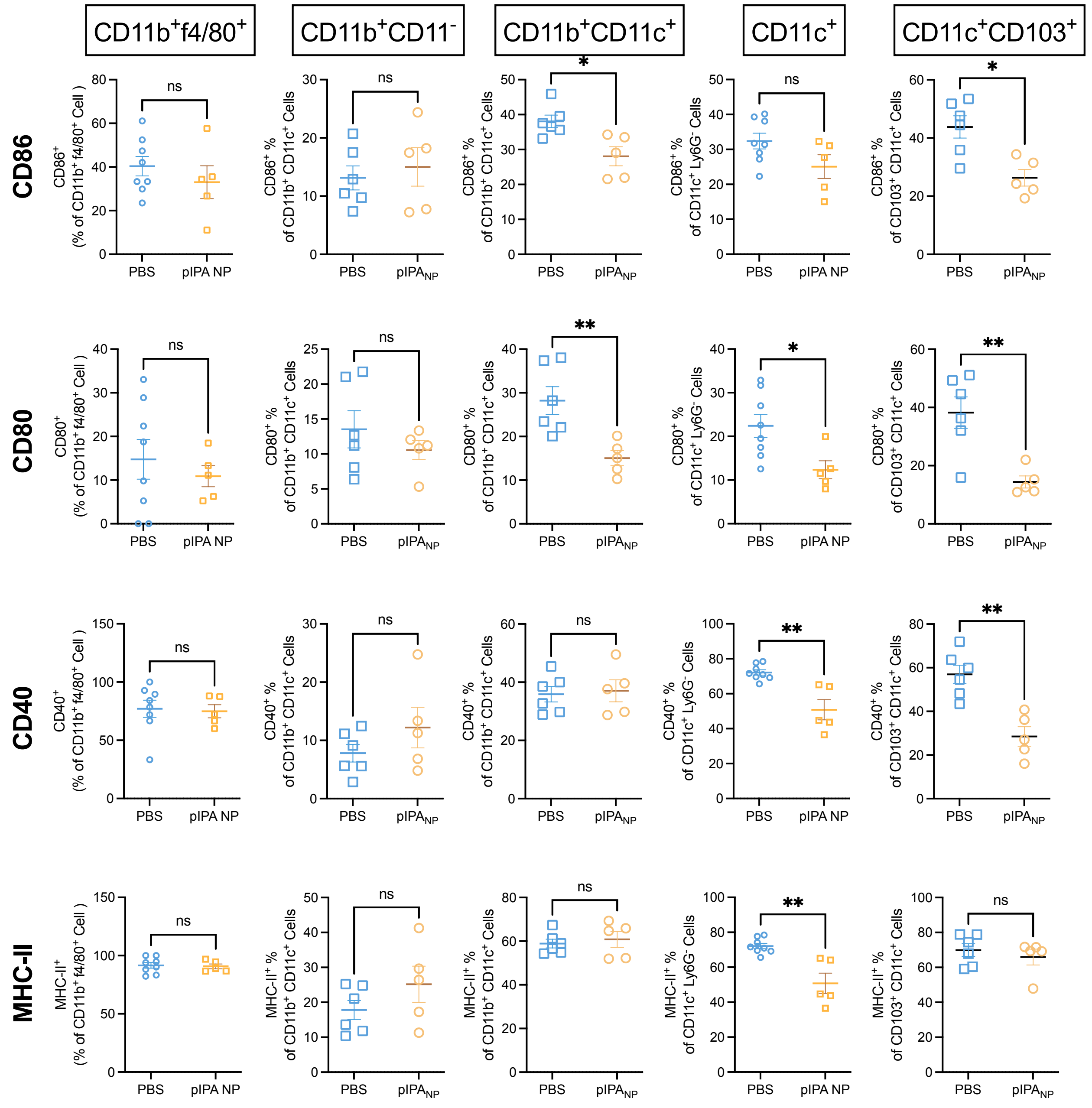


**Supplementary Figure S16. Percentages of costimulatory marker- and MHC-II-positive cells among inguinal lymph node myeloid subsets in the therapeutic CAIA study.** Percentage of $CD86^+$, $CD80^+$, $CD40^+$ and MHC-$II^+$ cells within $CD11b^+F4/80^+$, $CD11b^+CD11c^-$, $CD11b^+CD11c^+$, $CD11c^+$ and $CD103^+CD11c^+$ cells in spleen (n=6). The data correspond to the heatmap in Figure 4h. Data are presented as mean ± SEM. Groups were compared by a two-tailed Mann-Whitney test. *p $< 0.05$, **p $< 0.01$, ns, not significant.

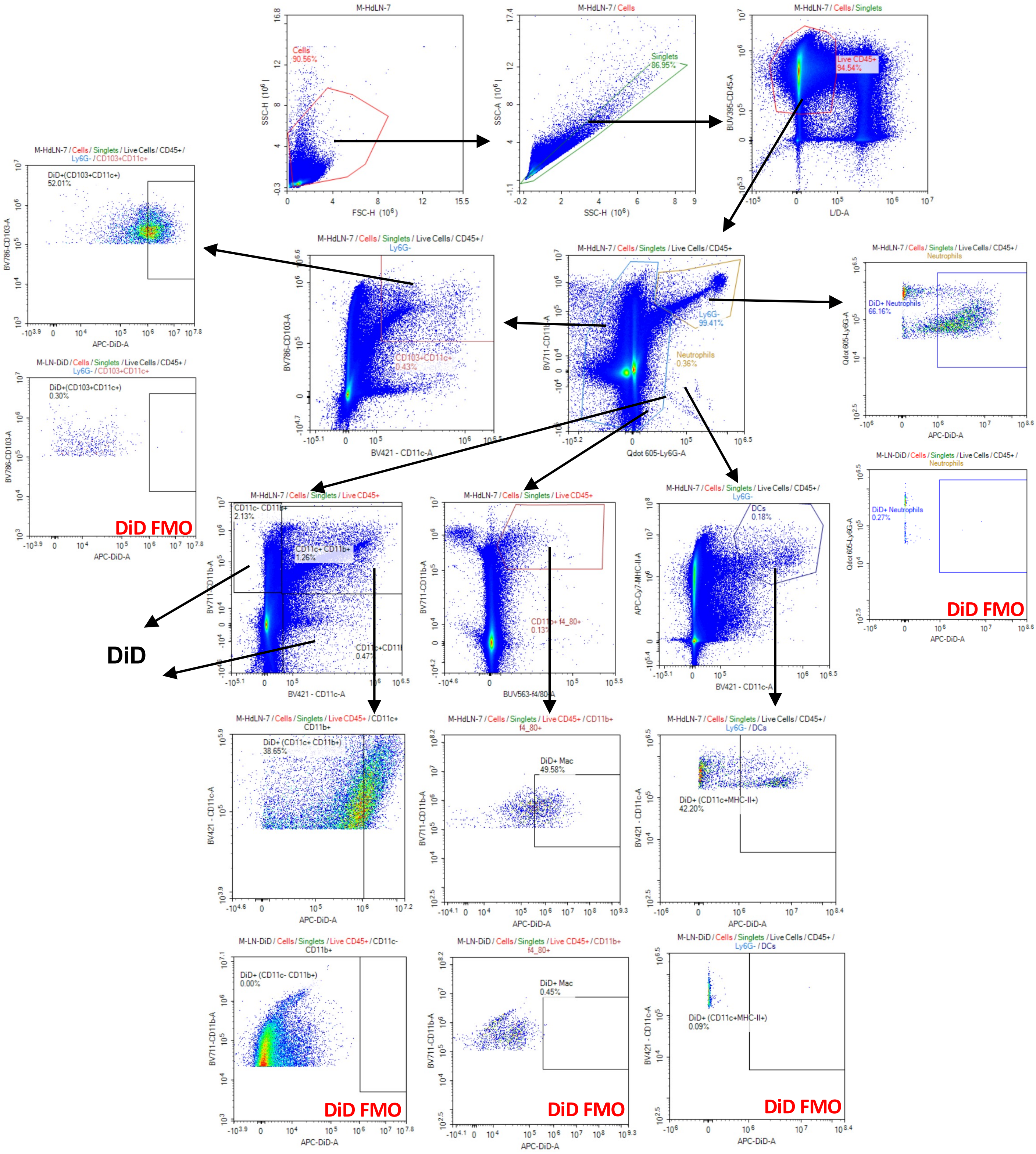


**Supplementary Figure S17. Flow cytometry gating strategy for the myeloid compartment in the cellular biodistribution study.** Myeloid subsets were defined as CD11b$^{+}$F4/80$^{+}$, CD11b$^{+}$CD11c$^{-}$, CD11b$^{+}$CD11c$^{+}$, CD11c$^{+}$CD11b$^{-}$, and CD103$^{+}$CD11c$^{+}$ cells, within which the percentages of DiD$^{+}$ cells were determined. This gating strategy applies to Figures 5a and 5b.

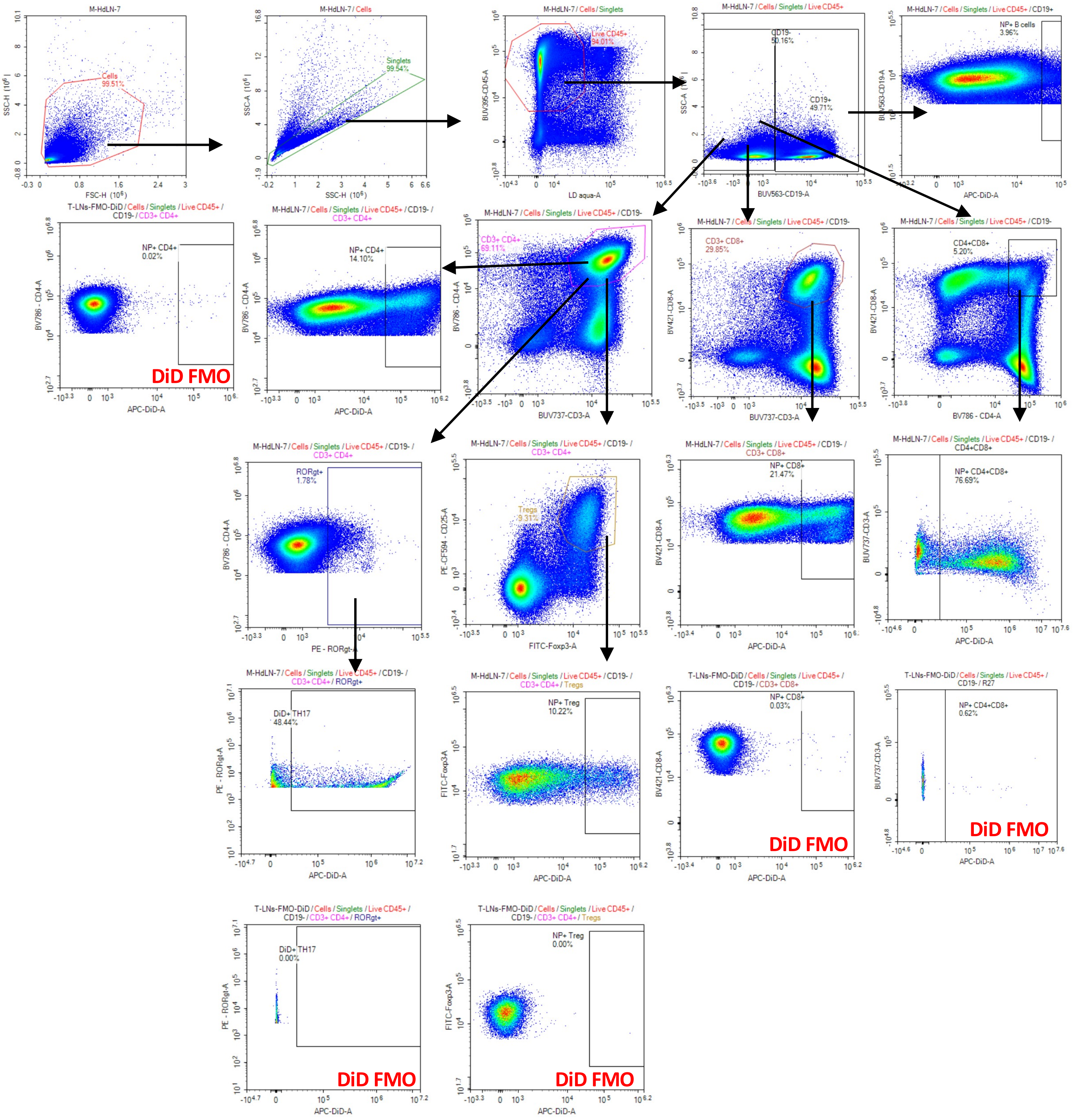


**Supplementary Figure S18. Flow cytometry gating strategy for the lymphocyte compartment in the cellular biodistribution study.** T cell and subsets were defined as CD19$^{+}$, CD4$^{+}$, CD8+, Treg, CD4$^{+}$CD8$^{+}$, and ROR$\gamma t^{+}$ cells, within which the percentages of DiD$^{+}$ cells were determined. This gating strategy applies to Figures 5a and 5b.

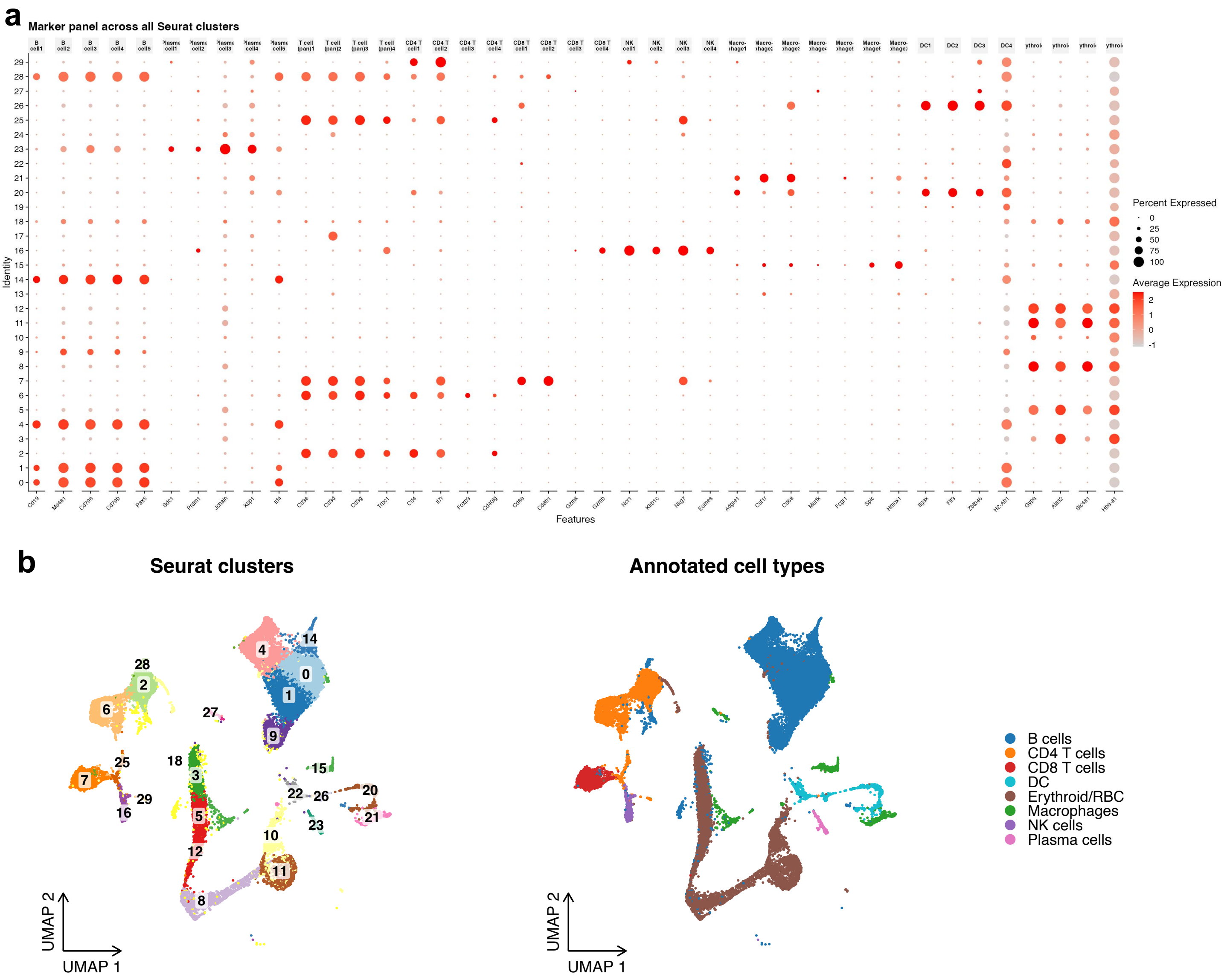


**Supplementary Figure S19. Cell-type annotation of splenic scRNA-seq data from CAIA mice.** Splenocytes from PBS- and pIPA NP-treated mice were analyzed by scRNA-seq (data pooled from two mice per group). **a.** Dot plot of canonical marker-gene expression across Seurat clusters. Dot size indicates the percentage of cells expressing each gene within a cluster, and color indicates average expression. **b.** UMAP representations colored by Seurat cluster (left) and broad cell-type annotation (right). Cell types were assigned manually using canonical marker genes and included B cells, plasma cells, CD4 T cells, CD8 T cells, NK cells, macrophages, dendritic cells, and erythroid cells.

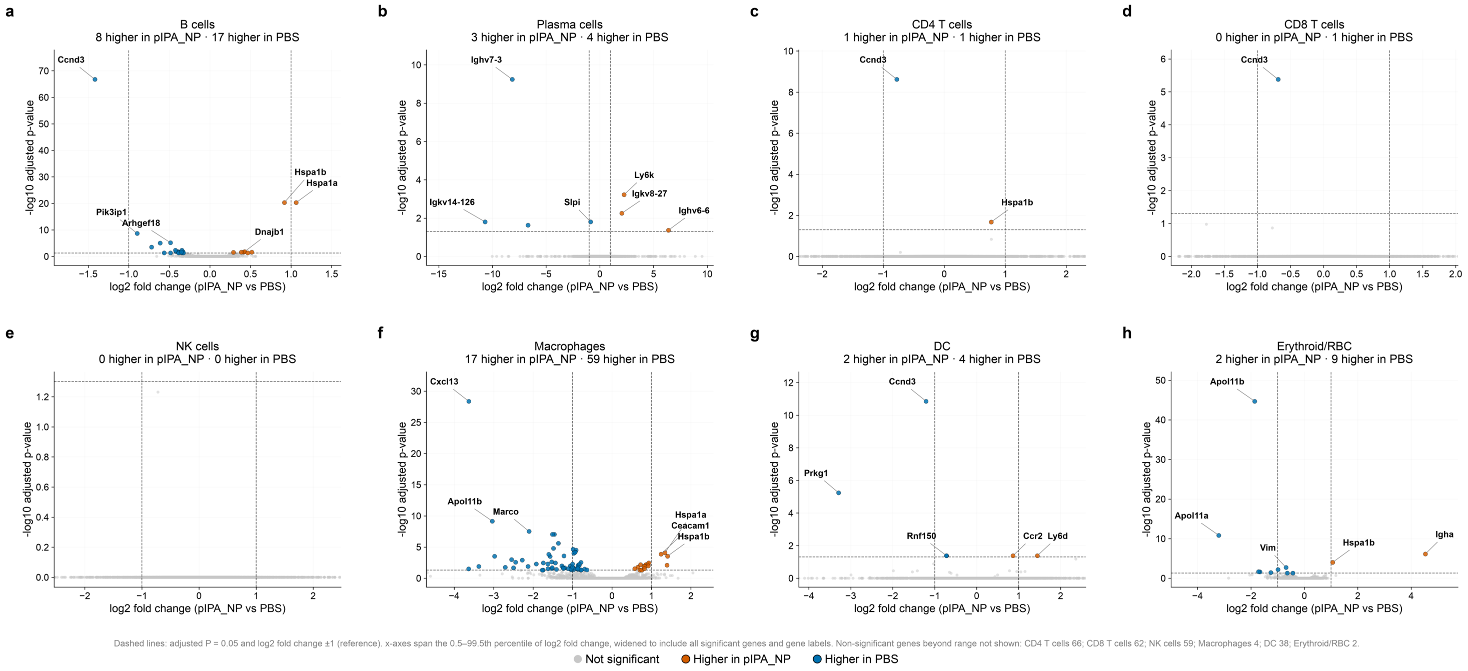


**Supplementary Figure S20. Cell-type-specific differential gene expression in spleens from pIPA NP- and PBS-treated CAIA mice.** Volcano plots show differential gene expression in B cells (**a**), plasma cells (**b**), CD4 T cells (**c**), CD8 T cells (**d**), NK cells (**e**), macrophages (**f**), dendritic cells (**g**), and erythroid cells (**h**). Counts were aggregated per mouse within each cell type and analyzed using DESeq2 (Wald test with Benjamini–Hochberg correction; two mice per group). Each point represents a gene. Orange and blue indicate significantly higher expression in the pIPA NP and PBS groups, respectively, at adjusted $P < 0.05$; gray indicates nonsignificant genes. Dashed lines indicate adjusted $P = 0.05$ and $\log_2$ fold change = ±1, with the latter shown as reference lines. Significant gene counts are indicated in each panel, and up to three significant genes in each direction are labeled.

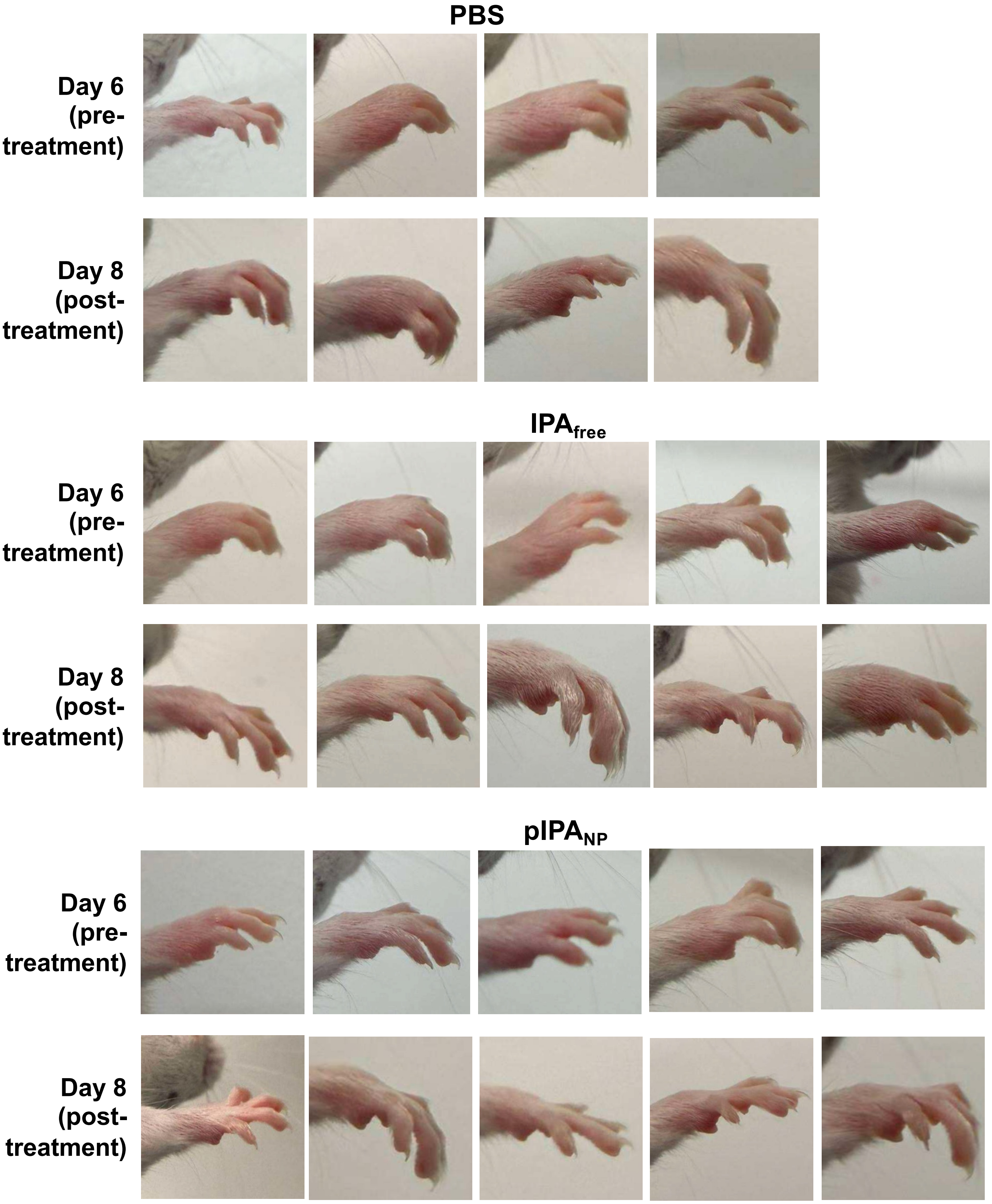


**Supplementary Figure S21. Representative photographs of mouse front paws from PBS-treated mice before and after treatment in the hock-injection study.** Photographs were taken on Day 6 (before treatment) and Day 8 (after treatment) with PBS administered subcutaneously near the hocks (Figure 6).

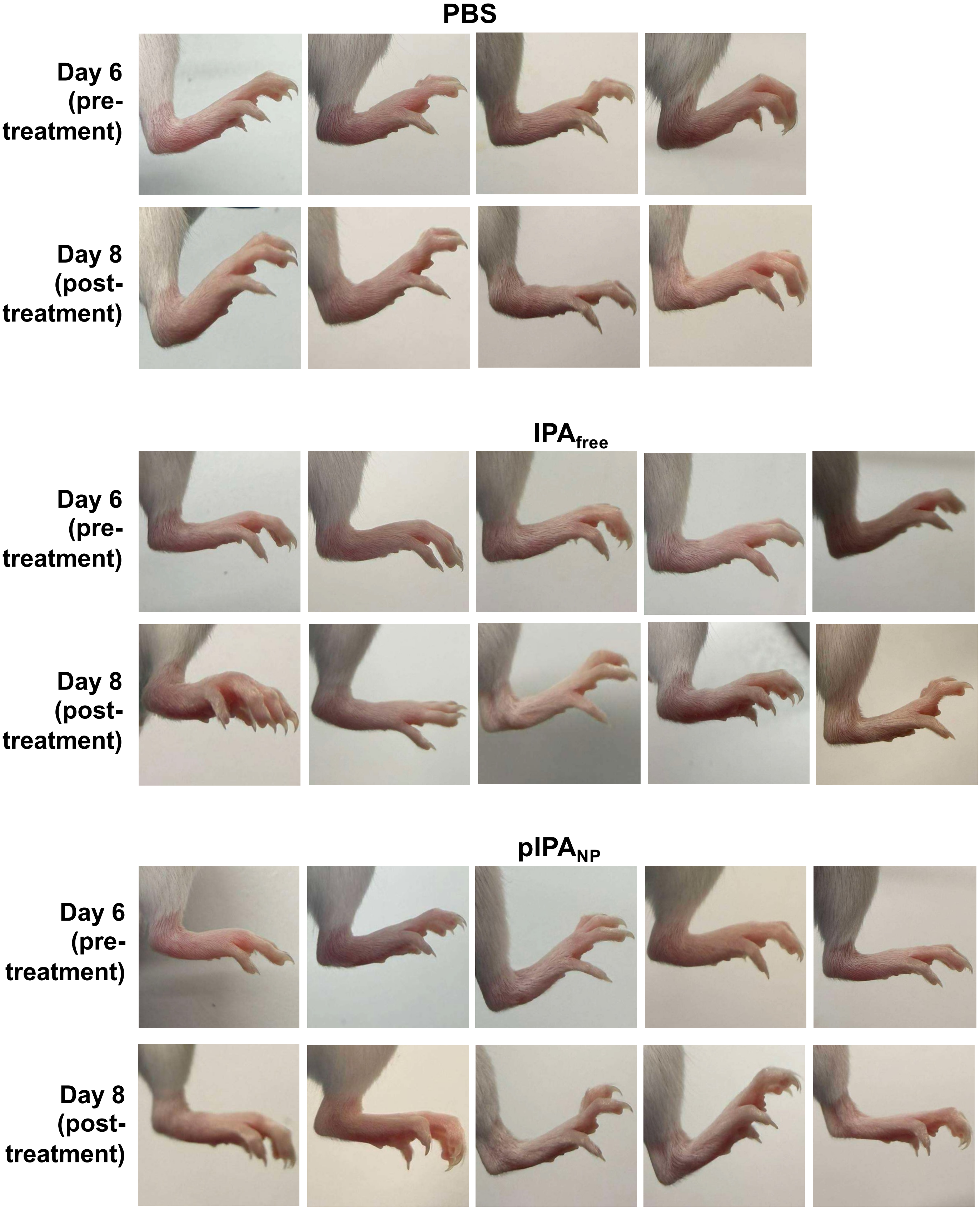


**Supplementary Figure S22. Representative photographs of mouse back paws from PBS-treated mice before and after treatment in the hock-injection study.** Photographs were taken on Day 6 (before treatment) and Day 8 (after treatment) with PBS administered subcutaneously near the hocks (Figure 6).

**Supplementary Table 1. Clinical arthritis scoring criteria for CAIA.** Each of the four paws is scored individually (0–4) based on the degree of swelling, and the total arthritis score is calculated as the sum of all four paw scores (range: 0–16). Adapted from Chondrex, Inc. https://www.chondrex.com/.

| Arthritis score | Condition |
|---|---|
| **0** | No swelling; paw appears healthy |
| **1** | Mild but definite swelling of a single joint type (e.g., digit, wrist, or ankle) |
| **2** | Mild but definite swelling of two joint types, or moderate swelling of a single joint type |
| **3** | Severe swelling involving multiple joint types |
| **4** | Maximal swelling of the entire paw, with the mouse unable to grip |

**Supplementary Table 2. Differentially expressed genes in splenic macrophages following pIPA NP treatment.** Counts were aggregated per mouse within cells annotated as macrophages and analyzed using DESeq2 (Wald test with Benjamini-Hochberg correction; Data pooled from two mice per group). The table lists genes with adjusted P < 0.05 in ascending order of adjusted P. Positive and negative $log_2$ fold changes indicate higher expression in the pIPA NP and PBS groups, respectively.

| Cell type | Gene | $log_2$ fold change | Adjusted P value | Higher in |
|---|---|---|---|---|
| Macrophages | *Cxcl13* | -3.63054321565905 | 4.15157357263072E-29 | PBS |
| Macrophages | *Apol11b* | -3.0343267400622800 | 6.92873953225163E-10 | PBS |
| Macrophages | *Marco* | -2.1013689749887300 | 3.07216174615081E-08 | PBS |
| Macrophages | *Slc40a1* | -1.4598626134404800 | 9.2816275667575E-08 | PBS |
| Macrophages | *C1qa* | -1.5110965983464800 | 9.2816275667575E-08 | PBS |
| Macrophages | *AW112010* | -1.3599994829747600 | 2.56133737617485E-06 | PBS |
| Macrophages | *Fcna* | -1.4862406527985000 | 1.66740625662884E-05 | PBS |
| Macrophages | *Hbb-bs* | -0.986648503674491 | 2.2710273490583E-05 | PBS |
| Macrophages | *Hba-a1* | -0.9116251387396790 | 3.16915469926221E-05 | PBS |
| Macrophages | *Hba-a2* | -0.9322025549973260 | 6.49109860340795E-05 | PBS |
| Macrophages | *Alas2* | -0.9367818301727140 | 6.81634433486821E-05 | PBS |
| Macrophages | *Hspa1a* | 1.3400174233777200 | 8.39757842759366E-05 | pIPA_NP |
| Macrophages | *Ccnd3* | -0.9681999497575940 | 9.88212867472827E-05 | PBS |
| Macrophages | *Ceacam1* | 1.243604760003220 | 0.00013595489737201600 | pIPA_NP |
| Macrophages | *Cadm1* | -1.6051602948571100 | 0.00014893780070745800 | PBS |
| Macrophages | *Vcam1* | -1.2173247553665800 | 0.00027026625424431500 | PBS |
| Macrophages | *Hspa1b* | 1.407821798600100 | 0.00028933569941435300 | pIPA_NP |
| Macrophages | *Gm47782* | -2.976903386259820 | 0.0002972722899787520 | PBS |
| Macrophages | *Itgad* | -1.5701685491593500 | 0.0003197350719399130 | PBS |
| Macrophages | *Hs3st2* | -2.5449627159259800 | 0.0010107841185364400 | PBS |
| Macrophages | *Gbp2b* | -2.2754125832346900 | 0.0012207946119144800 | PBS |
| Macrophages | *Arap2* | -1.527113861298420 | 0.0022897082632480000 | PBS |
| Macrophages | *Serpina3g* | -2.4424538112976400 | 0.0027651617748039900 | PBS |
| Macrophages | *Mkrn1* | -0.7887681092324420 | 0.002835851817246300 | PBS |
| Macrophages | *Itgal* | 0.933869521010597 | 0.0035544073930323500 | pIPA_NP |
| Macrophages | *Timd4* | -1.4373897975936000 | 0.0035544073930323500 | PBS |
| Macrophages | *Ccr3* | -1.7423371392690000 | 0.0035544073930323500 | PBS |
| Macrophages | *Itga9* | -1.6272270640996700 | 0.005007725513869190 | PBS |
| Macrophages | *Slc25a37* | -1.0008294935186300 | 0.005060732270685610 | PBS |
| Macrophages | *Fosb* | 0.9112434427980930 | 0.005218123720880210 | pIPA_NP |
| Macrophages | *Prkg1* | -1.9174821828751900 | 0.005218123720880210 | PBS |
| Macrophages | *Hbb-bt* | -0.8332627620339950 | 0.0060200437407635800 | PBS |
| Macrophages | *Klf4* | 0.857162981129754 | 0.006494176620041330 | pIPA_NP |
| Macrophages | *Adgre5* | 0.7249768218934250 | 0.006494176620041330 | pIPA_NP |
| Macrophages | *Gm56830* | 1.3928653235387100 | 0.007849084820376600 | pIPA_NP |
| Macrophages | *Il1b* | 0.8482156459325880 | 0.008468657693685630 | pIPA_NP |
| Macrophages | *Bpgm* | -0.8298536788297720 | 0.00849015423883214 | PBS |
| Macrophages | *Ube2l6* | -0.8620360780784120 | 0.00849015423883214 | PBS |
| Macrophages | *Cx3cr1* | 0.9171593763082790 | 0.010303852307392700 | pIPA_NP |
| Macrophages | *Clec4n* | -1.0561087790104600 | 0.010303852307392700 | PBS |
| Macrophages | *Bach2* | -1.2730639631010000 | 0.010402085622452100 | PBS |
| Macrophages | *Nfkbiz* | 0.8282424549132480 | 0.01274989138765940 | pIPA_NP |
| Macrophages | *Ccl6* | 0.8063059591711970 | 0.012809839081924700 | pIPA_NP |
| Macrophages | *Mfge8* | -2.1276524119511300 | 0.012809839081924700 | PBS |
| Macrophages | *C4b* | -3.3748993992681700 | 0.012809839081924700 | PBS |
| Macrophages | *Slc4a1* | -1.0054224868761800 | 0.015360467195383700 | PBS |
| Macrophages | *Rsad2* | -0.860669954018846 | 0.01692937698761860 | PBS |
| Macrophages | *Jun* | 0.6562837732950280 | 0.017012681791059400 | pIPA_NP |
| Macrophages | *Ubd* | -2.70637807840803 | 0.01715243669430290 | PBS |
| Macrophages | *Sash1* | -1.245709798336030 | 0.02001199482805430 | PBS |
| Macrophages | *Epb41l3* | -1.5138412139671900 | 0.020293835762929400 | PBS |
| Macrophages | *Cd55* | -1.2011854850484900 | 0.02085738640980830 | PBS |
| Macrophages | *P2ry14* | -2.4989707543712100 | 0.02085738640980830 | PBS |
| Macrophages | *Hebp1* | -0.9717220242297920 | 0.022289878791235800 | PBS |
| Macrophages | *Gfra2* | -1.5265062437036300 | 0.027113625474120400 | PBS |
| Macrophages | *Igf1* | -1.2303263161309100 | 0.028508733634573400 | PBS |
| Macrophages | *Fech* | -0.8667264340083900 | 0.029479938647800100 | PBS |
| Macrophages | *Sat1* | 0.5760697435851810 | 0.029650863310238200 | pIPA_NP |
| Macrophages | *Mbp* | 0.8203438801471470 | 0.030168885045170800 | pIPA_NP |
| Macrophages | *Axl* | -1.1083316174889600 | 0.030168885045170800 | PBS |
| Macrophages | *Cd5l* | -1.620473968803870 | 0.030244012706274500 | PBS |
| Macrophages | *Cd24a* | -0.8845549835180250 | 0.031195756611409100 | PBS |
| Macrophages | *Spic* | -1.0412460385043200 | 0.031195756611409100 | PBS |
| Macrophages | *Iigp1* | -3.6321845279692400 | 0.031195756611409100 | PBS |
| Macrophages | *Epb41* | -0.6925069720056060 | 0.03318856587137500 | PBS |
| Macrophages | *Cd81* | -0.859044524866335 | 0.03747678696069250 | PBS |
| Macrophages | *Pax5* | -1.4071086284895600 | 0.03934549196675840 | PBS |
| Macrophages | *Frmd4b* | -0.9749435853310850 | 0.04089100958037120 | PBS |
| Macrophages | *Sorbs2* | -1.0346198850702500 | 0.042359763428055000 | PBS |
| Macrophages | *Bnip3l* | -0.6342520491100940 | 0.04786474702081550 | PBS |
| Macrophages | *Slc25a39* | -0.7837520608530680 | 0.04786474702081550 | PBS |
| Macrophages | *Ccl9* | 0.7543583896063940 | 0.048553344946727000 | pIPA_NP |
| Macrophages | *Rap1gap2* | 0.7111378061496850 | 0.048553344946727000 | pIPA_NP |
| Macrophages | *Tent5c* | -0.7867772403530330 | 0.048553344946727000 | PBS |
| Macrophages | *Rapgef4* | -1.7480012451708100 | 0.048553344946727000 | PBS |
| Macrophages | *Sox5* | -1.7693777042703100 | 0.04967332988219640 | PBS |
